# Quantifying spectral information about source separation in multisource odour plumes

**DOI:** 10.1101/2024.01.14.575605

**Authors:** Sina Tootoonian, Aaron C. True, John P. Crimaldi, Andreas T. Schaefer

**Affiliations:** Sensory Circuits and Neurotechnology Laboratory, The Francis Crick Institute, London, UK; Department of Civil, Environmental and Architectural Engineering, University of Colorado, Boulder, USA; Department of Neuroscience, Physiology and Pharmacology, University College London, UK

## Abstract

Odours released by objects in natural environments can contain information about their spatial locations. In particular, the correlation of odour concentration fields produced by two spatially separated sources contains information about the distance between the sources. Mice are able to distinguish correlated and anti-correlated odour fluctuations at frequencies up to 40 Hz. Can this high-frequency acuity support odour source localization? Here we answer this question by quantifying the spatial information about source separation contained in the spectral constituents of correlations. We used computational fluid dynamics simulations of multisource plumes in two-dimensional chaotic flow environments to generate temporally complex, covarying odour concentration fields. By relating the correlation of these fields to the spectral decompositions of the associated odour concentration timeseries, and making simplifying assumptions about the statistics of these decompositions, we derived analytic expressions for the Fisher information contained in the spectral components of the correlations about source separation. We computed the Fisher information for a broad range of frequencies and source separations and found that high frequencies were more informative than low frequencies when sources were close relative to the sizes of the large eddies in the flow. We observed a qualitatively similar effect in an independent set of simulations with different geometry, but not for surrogate data with a similar power spectrum to our simulations but in which all frequencies were *a priori* equally informative. Our work suggests that the high-frequency acuity of the murine olfactory system may support high-resolution spatial localization of odour sources. We also provide a model of the distribution of the spectral components of correlations that is accurate over a broad range of frequencies and source separations. More broadly, our work establishes an approach for the quantification of the spatial information in odour concentration timeseries.

## 1 Introduction

Olfactory signals are crucial to the behaviours of many animals, whether for seeking mates, nurturing young, avoiding predators, finding food, or navigating complex environments, to list a few important examples. Odours are often conveyed to animals by turbulent flows. Odours released into turbulent flow environments are advected downstream with the mean flow. Concurrent distortion of odour streams by chaotic flow structures spanning a range of sizes (a defining characteristic of fluid turbulence) produces discrete odour filaments whose structure is subsequently altered by the turbulent mixing field. The associated stretching and folding of filaments sharpens odour concentration gradients, enhancing molecular diffusion in response to these flow-sharpened gradients and acting to broaden filaments. The net effect of these competing processes, collectively constrained by odour mass conservation, is broadly referred to as chaotic advection, turbulent diffusion, or simply *mixing* [1, 2, 3], and the resultant product is a complex, spatiotemporally dynamic concentration landscape. We refer to these as *odour landscapes* or *plumes*.

Statistics describing concentration fluctuations of passive scalar plumes (composed of odours that do not modify the local flow field) and their spatial variation are set by a number of dimensionless parameters. These include a Reynolds number (Re) describing the ambient flow environment (setting the statistics of the turbulent flow field) and a Péclet number (Pe) describing the relative importance of advective and diffusive odour transport (or alternatively a Schmidt number describing the relative diffusivities of fluid momentum and odour mass). Higher Re and Pe numbers are associated with more turbulent flows producing plume statistics dominated by turbulent diffusion. Additionally, a number of ratios describe the *source configuration*. These include its size relative to the largest, energy-containing eddies (characterized by an integral length scale), its proximity to solid boundaries, and its injection momentum relative to the flow. Source configuration effects are well-known (e.g. geometry [4, 5, 6], source momentum relative to the mean flow [7], and proximity to surfaces [8, 9, 10]). The variety of experimentally-measured odour landscapes shown in Fig 1 illustrates the effects of some of these important factors.

**Fig 1.**
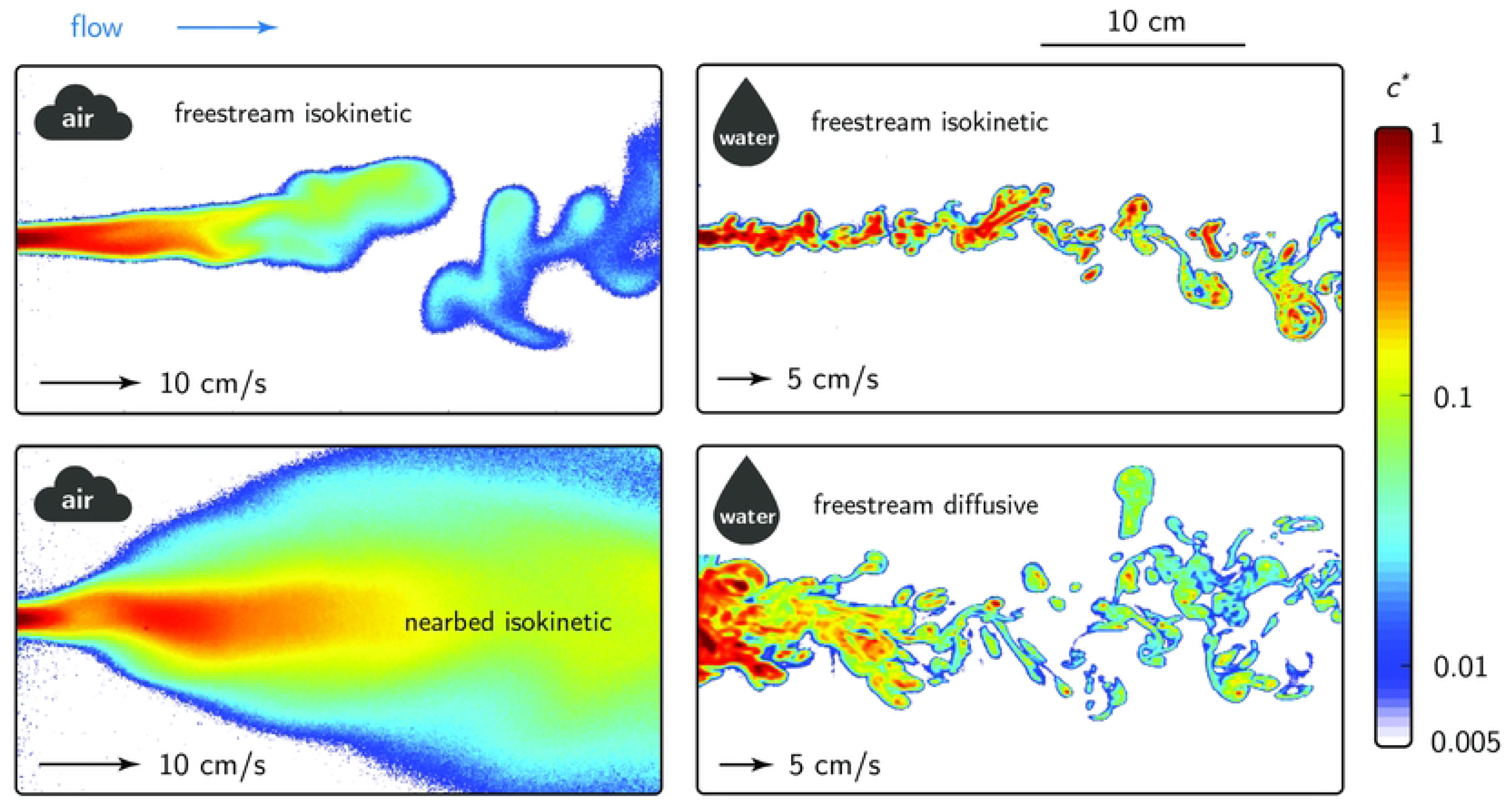
Variety of odour landscapes. Diverse single odour landscapes in air and water (adapted from [11]) with varying source release configurations and flow, fluid, and odourant properties. Animals may have to operate in multiple landscapes. For example, the odour landscape relevant to a mouse may be similar to the nearbed isokinetic condition (lower left) when it is searching the soil, and more like the freestream isokinetic (upper left) when it is rearing. The plumes were measured experimentally using planar laser-induced fluorescence (PLIF) and are described in detail in [10] (air) and [5] (water).

Plume statistics exhibit characteristic variations with distance from the source — including with streamwise and cross-stream location, parallel and normal to the mean flow vector, respectively — owing to transitions between phenomenological regimes defined by differing sources of concentration variance. For example, for a small relative source size, near the source where the plume width is small relative to the large eddies, concentration variance is primarily driven by meandering of the whole plume driven by those large eddies (e.g. [12]). As the plume width grows with distance from the source, the large eddies produce less meandering and instead a range of eddy sizes are effective at mixing odour-laden and ambient (clean) fluid — a turbulent diffusive regime with enhanced relative dispersion of the plume locally around its meandering centroid. A number of studies have examined statistical structure with distance from the source [13, 14, 15, 16, 17]. Broadly speaking, source configuration effects influence near-field plume statistics (close to the source), whereas the far-field structure is largely self-similar, being set by the physics of turbulent flows and the linearly coupled advection-diffusion equations governing odourant transport and diffusion.

When multiple non-reactive odours are released from initially-distant locations into turbulent flows, odour-specific plumes evolve independently and uniquely under the influence of chaotic coherent flow structures. The nature of the cumulative odour landscape describing all local odour concentrations and associated gradients is then simply the superposition of these odour-specific plumes, information that is potentially exploitable by a navigating sensor or organism. The relevance of various instantaneous and statistical plume properties as exploitable cues for organisms during odour source localization (odour navigation) is an important and active area of research in a number of disciplines, including olfactory neuroscience [15, 18, 19, 20, 21, 22, 23].

Of particular relevance to the work presented here is that, in addition to the factors setting the overall statistics of each individual odour plume, pointwise concentration correlations between odour species will depend strongly on the initial source separation distance. [24, 25, 26, 27, 28]. In particular, the correlations of concentration fluctuations can vary non-monotonically with distance from the source, and they show strong source separation effects with weak Reynolds number effects [27, 29, 30]. This contributes to the observations of zero, negative, and positive correlation regimes (no interaction, destructive and constructive interference, respectively) [24]. As an analogue to the single-source case where the local plume width relative to the large eddy sizes is important in driving local production of concentration variability, the source separation relative to the large eddy sizes in the multisource case drives these complex correlation behaviors.

Many statistical models describing concentration fluctuations for the single-source case have been proposed in the literature, and most leverage the simplicity of the exponential family to describe the one-point probability density functions [31, 32]. While the best match for a particularly distance from the source and plume realization (Re, Pe, source configuration) varies, there is recent evidence that the flexibility of the Gamma distribution can robustly account for much of this variability [33], where the shape parameter is related to the concentration fluctuation intensity [4]. Intuitively, good statistical descriptors for concentration fluctuations must also capture the intermittent nature of concentration dynamics in turbulent flows manifesting as high probabilities of zero concentration events. [32]. Extensions of these models to describe the concentration statistics of multiple interacting sources are less frequent in the literature, but the exponential family was also shown to provide a good description of the distributions for the sum of concentrations [25, 34]. Even fewer studies have examined the spectral properties of correlations; however, recent work showed that for a single source separation, correlations increase with distance from the source and become more spectrally uniform [27].

As noted early by Hopfield [35], the modulation of correlation by intersource distance can help animals locate odour sources in the environment i.e. perform olfactory scene segmentation. But do animals use such correlations to perform odour source localization? Recently it was shown that mice can discriminate fast odour fluctuations at frequencies of up to 40 Hz, much higher than expected for a so-called ‘slow’ modality like olfaction [36]. What is the function of such high bandwidth sensitivity? Odour plumes in naturalistic environments contain fluctuations at a wide range of frequencies, modulated by distance to the source, proximity to solid boundaries, odour source characteristics, and the ambient flow environment. One possibility is that information about intersource distance is better encoded in different frequency bands depending on the olfactory context, and that the sensitivity of the mouse olfactory system to rapidly fluctuating odours allows it to use this information when performing odour source localization. Thus the question we aim to answer in this study is: *What is the spectral distribution of the information that correlations carry about source separation?*

## 2 Results

To provide a testing ground for our approach to quantifying information in odour plumes we performed computational fluid dynamics (CFD) simulations of two-dimensional fluid flowing past equally spaced cylindrical obstacles in a rectangular wind tunnel, as shown in Fig 2. The interaction of the flow with the obstacles generated complex flow patterns that we used as a proxy for turbulent advection. A snapshot of such a flow pattern is shown in Fig 3A. We placed multiple odour sources in the simulated wind tunnel at a fixed distance from the flow inlet and at various locations from the midline of flow and measured the correlation of the resulting concentration profiles — henceforth referred to as *plumes* — at a fixed location down wind. In Fig 3B we show some example plumes from sources at three different intersource distances. We express all distances in terms of the spacing between the obstacles, called the ‘pitch,’ and indicate this by suffixing all distances with ‘*ϕ*’. This is a natural length scale for distance normalization since it approximates the sizes of the largest eddies.

**Fig 2.**
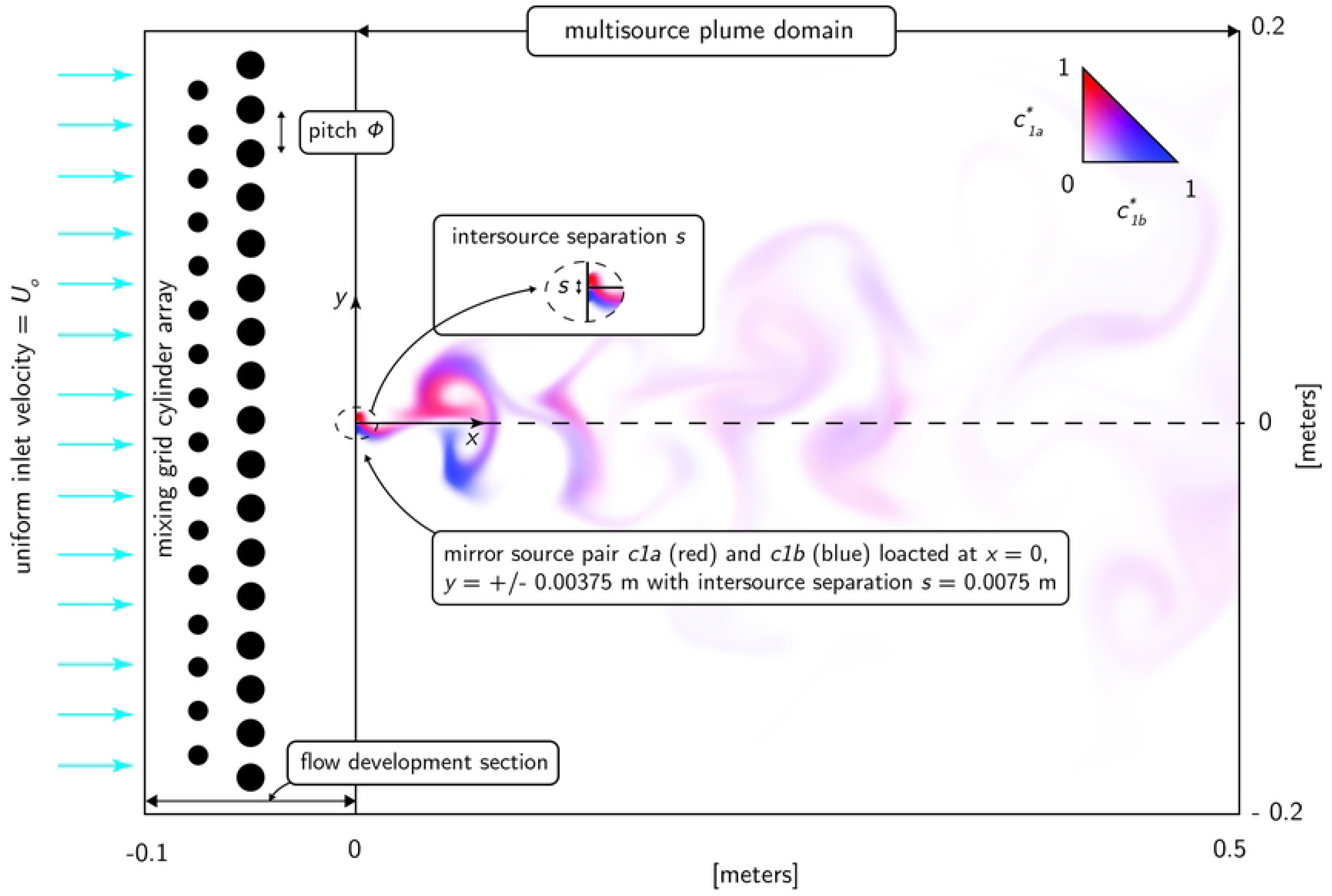
CFD simulation domain. The wind tunnel domain used to simulate multispecies odour landscapes. The total domain consists of an inlet flow development section followed by a plume domain section, beginning at *x* = 0 from where an array of mirrored source pairs are distributed about the domain centerline at *y*=0. Geometric, flow, fluid, and odourant properties are summarized in Table 1.

**Fig 3.**
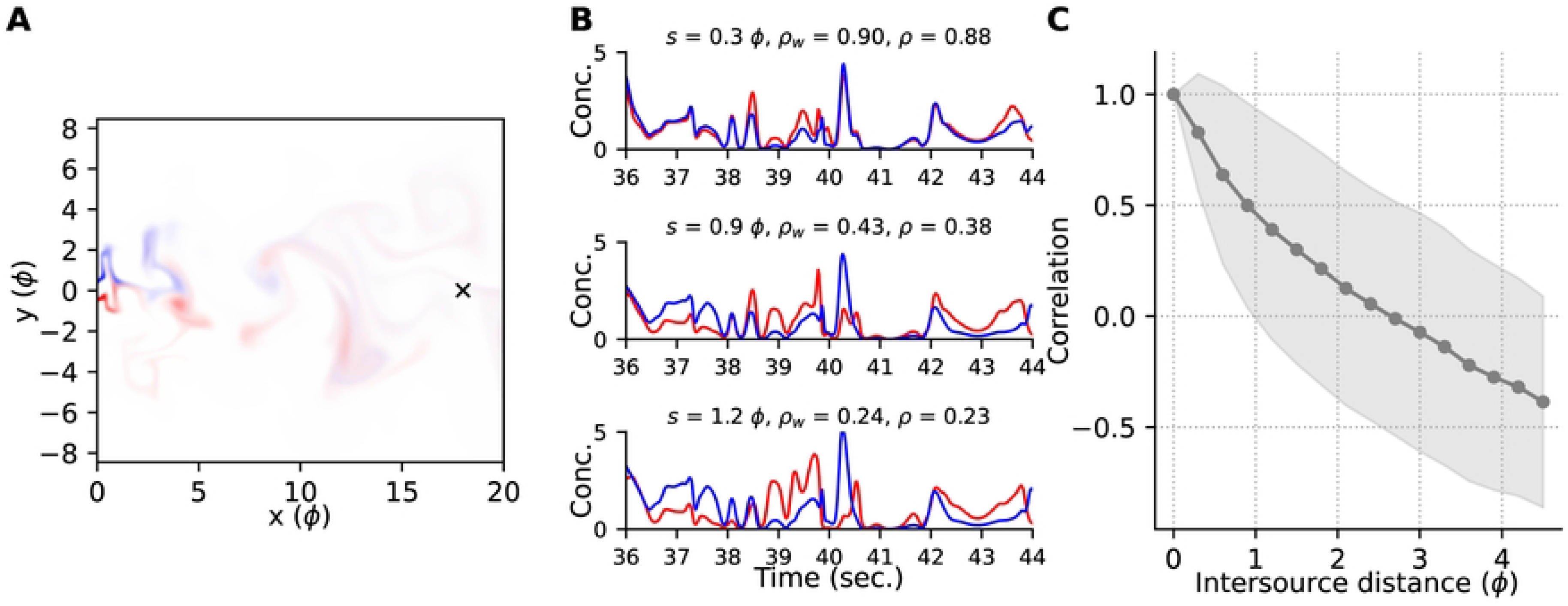
Example plumes and correlations. **(A)** Snapshot of the simulated flow patterns advecting two sources (colours). The plumes are sampled downwind at the probe location marked ‘*×*’. **(B)** Concentration profiles of the two odours at the probe location, scaled by the larger of their two standard deviations computed over the window shown. Sources were placed at the same downwind (horizontal) location and separated by crosswind (vertical) distances indicated by *s*. Pearson correlations computed for the time windows shown, and for the entire simulation duration, are indicated by *ρ*_*w*_ and *ρ*, respectively. **(C)** Mean *±* standard deviation of the Pearson correlations computed over 1-second boxcar windows overlapping by 500 msec., for all sources at the intersource distances indicated. All distances are measured in pitch (*ϕ*).

We used the Pearson correlation to measure the cofluctuations of two plumes. For the examples shown in Fig 3B correlations decrease with increasing intersource distance. We confirmed that this effect holds for the rest of our data in Fig 3C, similar to previous observations in the literature, e.g. [37].

**Table 1:**
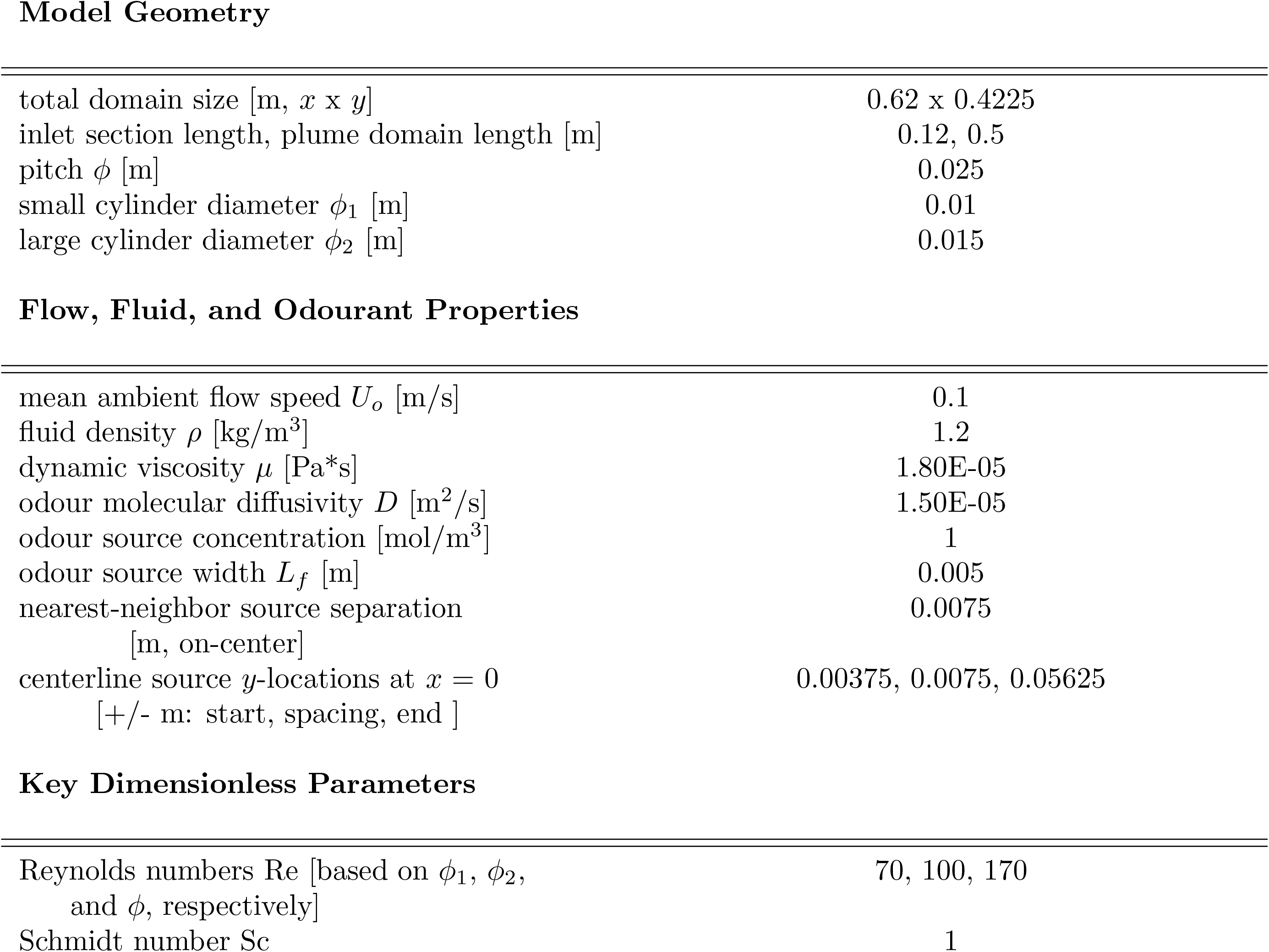
Summary of model geometry, flow, fluid, and odourant properties used for multisource plume simulations. Properties are for air and common airborne odourants at *≈* 20^*o*^C.

Model Geometry
|  |  |
| --- | --- |
| total domain size [m, $x \times y$] | 0.62 x 0.4225 |
| inlet section length, plume domain length [m] | 0.12, 0.5 |
| pitch $\phi$ [m] | 0.025 |
| small cylinder diameter $\phi_1$ [m] | 0.01 |
| large cylinder diameter $\phi_2$ [m] | 0.015 |
| Flow, Fluid, and Odourant Properties |  |
| mean ambient flow speed $U_o$ [m/s] | 0.1 |
| fluid density $\rho$ [kg/m<sup>3</sup>] | 1.2 |
| dynamic viscosity $\mu$ [Pa*s] | 1.80E-05 |
| odour molecular diffusivity $D$ [m<sup>2</sup>/s] | 1.50E-05 |
| odour source concentration [mol/m<sup>3</sup>] | 1 |
| odour source width $L_f$ [m] | 0.005 |
| nearest-neighbor source separation<br>[m, on-center] | 0.0075 |
| centerline source $y$-locations at $x = 0$<br>[+/- m: start, spacing, end ] | 0.00375, 0.0075, 0.05625 |
| Key Dimensionless Parameters |  |
| Reynolds numbers Re [based on $\phi_1$, $\phi_2$,<br>and $\phi$, respectively] | 70, 100, 170 |
| Schmidt number Sc | 1 |

**Table 2:**
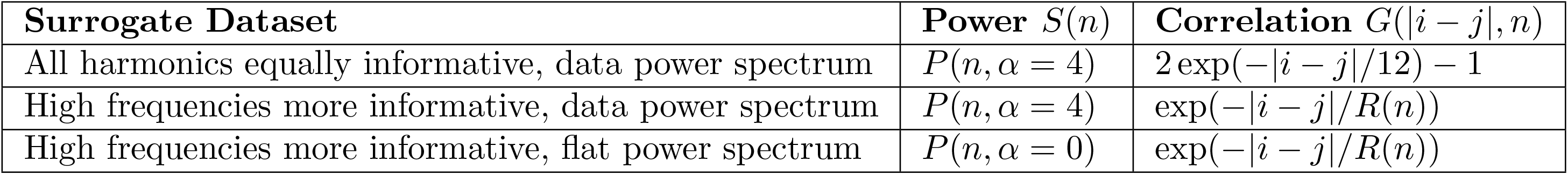
Summary of surrogate datasets and the power and correlation functions used. The kernel for each dataset was the product of the power and correlation functions: *k*(*i, j, n*) = *S*(*n*)*G*(|*i − j*|, *n*).

The Pearson correlation of two plumes *x*(*t*) and *y*(*t*) in a time window of width *T* seconds is

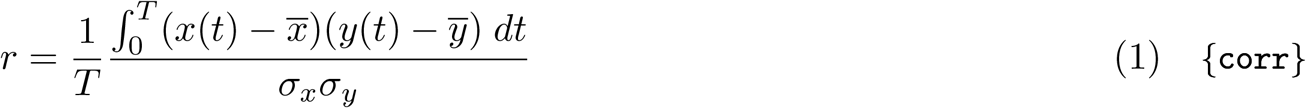

where 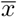 and *σ*_*x*_ are the mean and standard deviation of *x*(*t*) over this window, and similarly for *y* and *σ*_*y*_. A key property of the Pearson correlation for our purposes is that it can be decomposed as a sum of correlations computed for each harmonic of the fundamental frequency 1*/T*

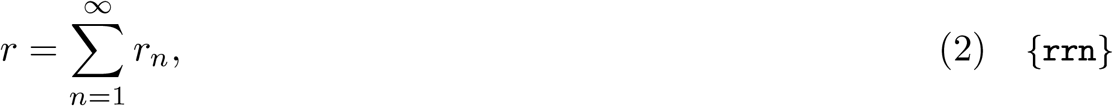

where *r*_*n*_ is the contribution from the *n*’th harmonic,

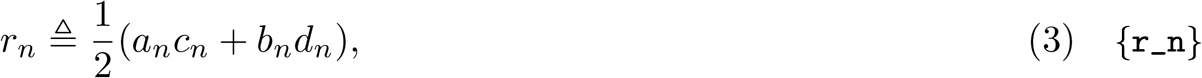

and (*a*_*n*_, *b*_*n*_) and (*c*_*n*_, *d*_*n*_) are the cosine and sine coefficients of the Fourier decomposition of *x*(*t*) and *y*(*t*), respectively, scaled by the standard deviations of these two signals (see Eqn 68 in Methods).

The spectral decomposition of plume correlations clearly depends on the corresponding decomposition of the plumes themselves. Observation of long duration signals such as plumes over short time windows introduces uncertainty about their spectral content called *spectral leakage* [38], wherein the spectral content of the observed signal at a given frequency is not just that of the original signal at that frequency, but includes weighted contributions from all other frequencies. The effects of spectral leakage can be ameliorated by judicious weighting of the observations, called ‘windowing’. Unless stated otherwise, the results below use a 1-second Hann window. In Fig 4 we have plotted the distribution of some of these correlation components as a function of distance.

**Fig 4.**
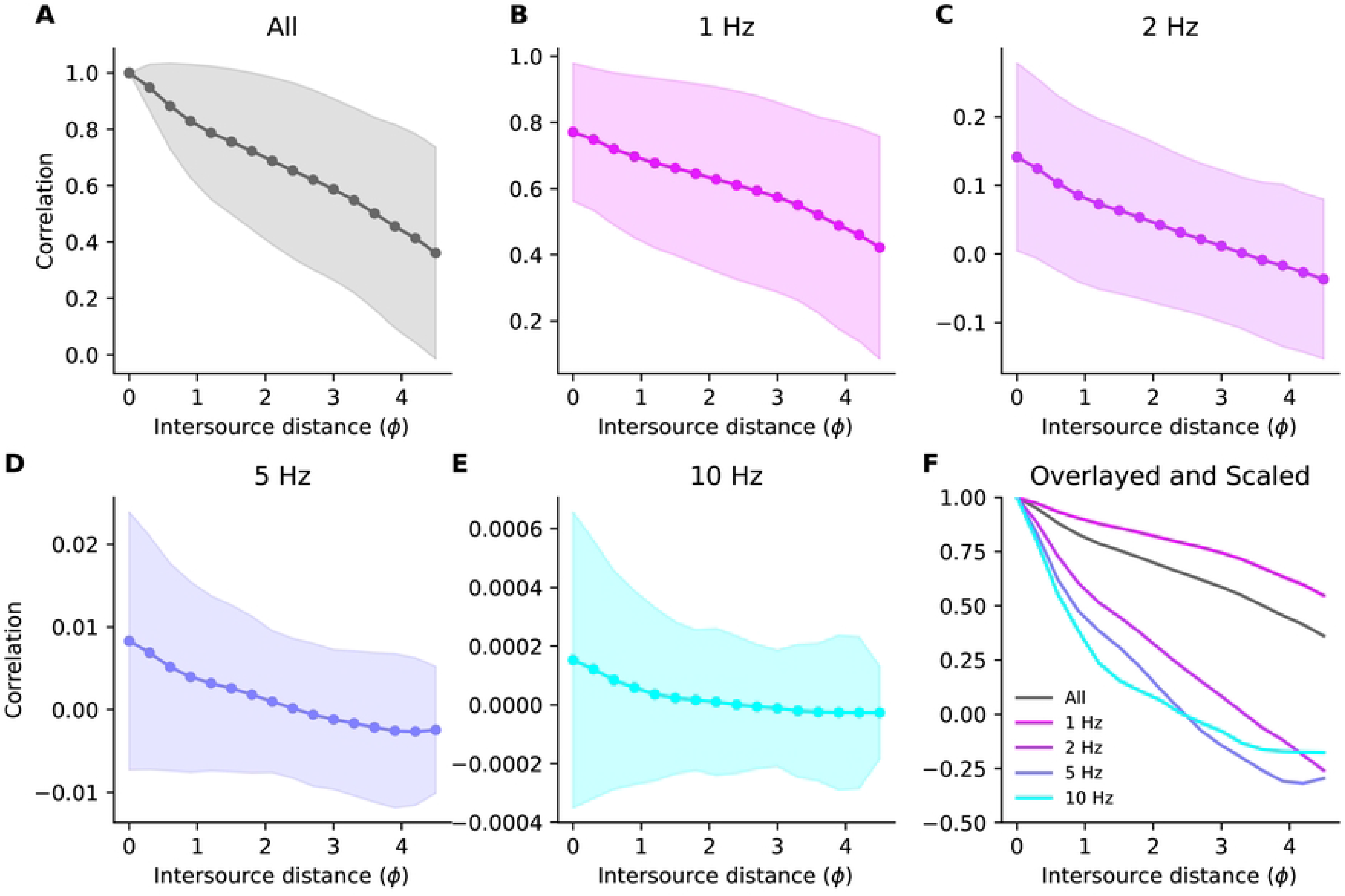
Decomposition of correlations. **(A)** Mean *±* S.D. of Pearson correlations between the plumes emitted by two sources the specified distance apart, computed using 1-second Hann windows (compare to Fig 3C which used 1-second boxcar windows). **(B–E)** The components of the correlation at the frequencies stated above each panel. **(F)** The means of the data in panels (B–E) scaled to have the same value at intersource distance zero and overlayed for ease of comparison.

To measure the amount of information that correlations *r*_*n*_ provide about intersource distance *s*, we used the Fisher information [39], defined as

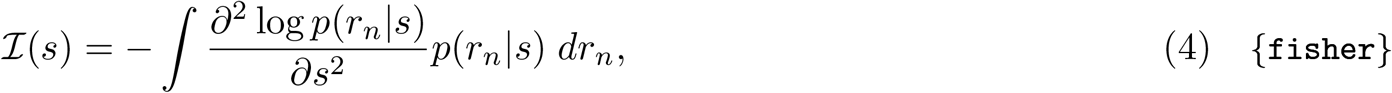

where *p*(*r*_*n*_|*s*) is the distribution of correlations *r*_*n*_ for harmonic *n* for an intersource separation *s*. Informally, the Fisher information measures the dependence of our estimate of *s* on the observed correlations *r*_*n*_. The more sensitive this estimate is to correlations, the more informative the correlations. The Fisher information also bounds the precision of unbiased estimation of intersource distance from correlations. Thus the higher the Fisher information, the higher the maximum precision with which intersource distance can be estimated from correlations. Note also that the Fisher information is a function of the intersource distance *s*. Thus the information provided by correlations will in general depend on the (true) intersource distance.

### 2.1 Outline of our approach

Our approach to determining the spectral distribution of information in the correlations is as follows:

1. Use simplifying assumptions about the statistics of the Fourier coefficients of the component waveforms to determine analytically tractable approximations to the distribution *p*(*r*_*n*_|*s*) needed to compute the Fisher information (Sec 2.2, Eqn 24, and Fig 7);
2. Use the distribution of correlations thus derived to compute analytic expressions for the Fisher information (Sec 2.3 and Eqn 32);
3. Fit tractable distributions for the quantities required in the analytic expressions of Fisher information to our simulation data (Eqn 33, Fig 8 and Fig 9);
4. Compute the Fisher information using fitted parameters for each harmonic and intersource distance (Eqn 35 and Fig 10).

To determine the extent to which our results generalized to other flows we performed a second set of simulations to those presented in the Main Text using similar flow parameters but slightly different geometry. In Supporting Information Sec S3 we describe these simulations and present some of the corresponding results of the analyses presented in the Main Text.

### 2.2 The distribution of correlations

To determine an analytically tractable expression for *p*(*r*_*n*_|*s*), we began by using the rules of probability to relate the correlation *r*_*n*_ to the Fourier coefficients *a*_*n*_, *b*_*n*_, *c*_*n*_ and *d*_*n*_,

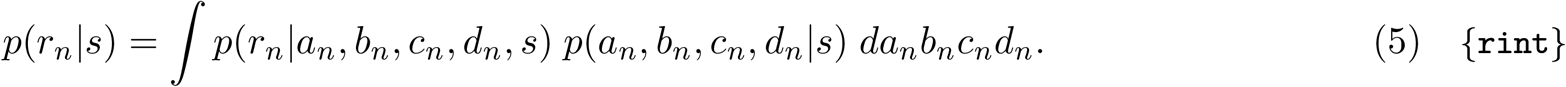

The first term in the integrand describes the dependence of correlations *r*_*n*_ on the Fourier coefficients and the intersource distance. From Eqn 3 we know that *r*_*n*_ is determined entirely by the Fourier coefficients. We can express this fact probabilistically by concentrating all of the probability density at the value in Eqn 3 using the Dirac *δ* function,

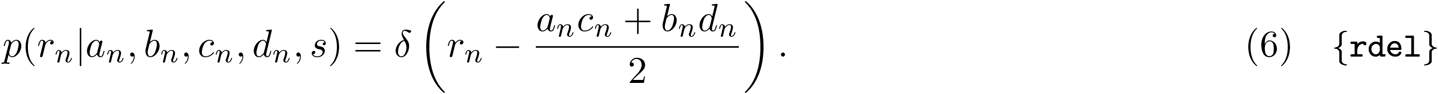

The second term in the integrand of Eqn 5 above is the joint distribution of coefficients from two sources *p*(*a*_*n*_, *b*_*n*_, *c*_*n*_, *d*_*n*_|*s*). It is the more interesting one because it describes the dependence of the Fourier coefficients on the intersource distance *s*. To emphasize the relationship of coefficients from one source, *a*_*n*_, *b*_*n*_, to those from the other, *c*_*n*_, *d*_*n*_ we decompose the joint distribution as,

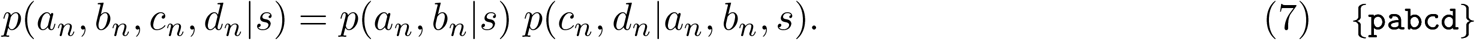

The first term on the right-hand side of Eqn 7 is the probability of observing the pair of coefficients *a*_*n*_, *b*_*n*_ from one source, given intersource distance *s*. To arrive at analytically tractable solutions, we made our first assumption:

1. Concentration profiles are Gaussian processes. This assumption implies that the joint distribution of coefficients from a given source, *p*(*a*_*n*_, *b*_*n*_|*s*), is a two-dimensional Gaussian distribution. In principle, the mean and covariance of this Gaussian can depend on the locations of the sources. To simplify further, we assumed that sources were close enough together relative to the animal and to the geometry of the flow such that the distribution of sine and cosine coefficients from each source is independent of its location. That is,
2. The distribution of coefficients from each source is the same for all sources,

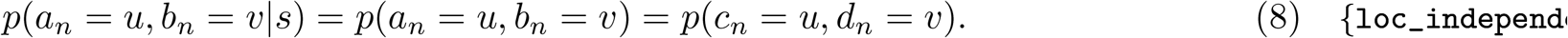

This assumption allowed us to replace *p*(*a*_*n*_, *b*_*n*_|*s*) in Eqn 7 with *p*(*a*_*n*_, *b*_*n*_). To specify the mean and covariance of this distribution, we made our next assumption,
3. Concentration profiles are temporally stationary. This means that the statistical properties of plumes do not change with time. Temporal stationarity has the following important implications for the Fourier coefficients (see Sec S5.1 for details). First, it implies that the marginal distribution of the sine and cosine coefficients must be the same, that is

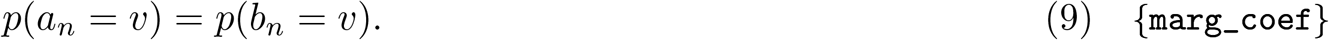

Second, it implies that the coefficients at non-zero frequency have mean zero, that is

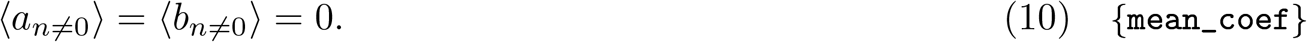

Third, it implies that the coefficients are uncorrelated, so The expectations in Eqn 10 and Eqn 11 are over time windows.

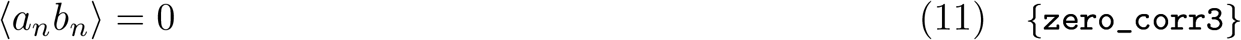

By combining our Gaussian process assumption with our assumptions of location independence and temporal stationarity we can completely specify the first term in Eqn 7:

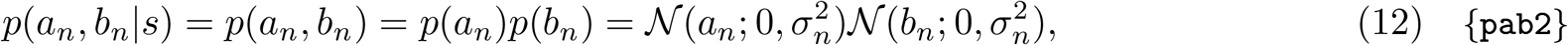

where 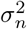 is the marginal variance of the coefficients for harmonic *n*. Location independence implies that the joint distribution of the coefficients *c*_*n*_ and *d*_*n*_ from the second source have the same form,

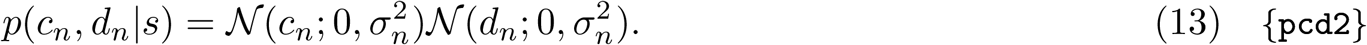

The second term in Eqn 7 is *p*(*c*_*n*_, *d*_*n*_|*a*_*n*_, *b*_*n*_, *s*) and expresses the probability of observing the pair of coefficients (*c*_*n*_, *d*_*n*_) from the second source given the observed coefficients (*a*_*n*_, *b*_*n*_) from the first source located at an intersource distance of *s*. To specify it, we first represented the decomposition of Eqn 7 as the graphical model in Fig 5A. The marginal independence of the sine and cosine coefficients expressed in Eqn 12 is reflected in the network through the lack of connections between *a*_*n*_ and *b*_*n*_, and between *c*_*n*_ and *d*_*n*_. Importantly, the network indicates that the coefficients *c*_*n*_ and *d*_*n*_ are conditionally independent given the observed values of *a*_*n*_ and *b*_*n*_. This is because the only routes by which *c*_*n*_ and *d*_*n*_ can influence each other are through *a*_*n*_ and *b*_*n*_, which serve as common input to *a*_*n*_ and *b*_*n*_, and these are observed [40]. Therefore,

**Fig 5.**
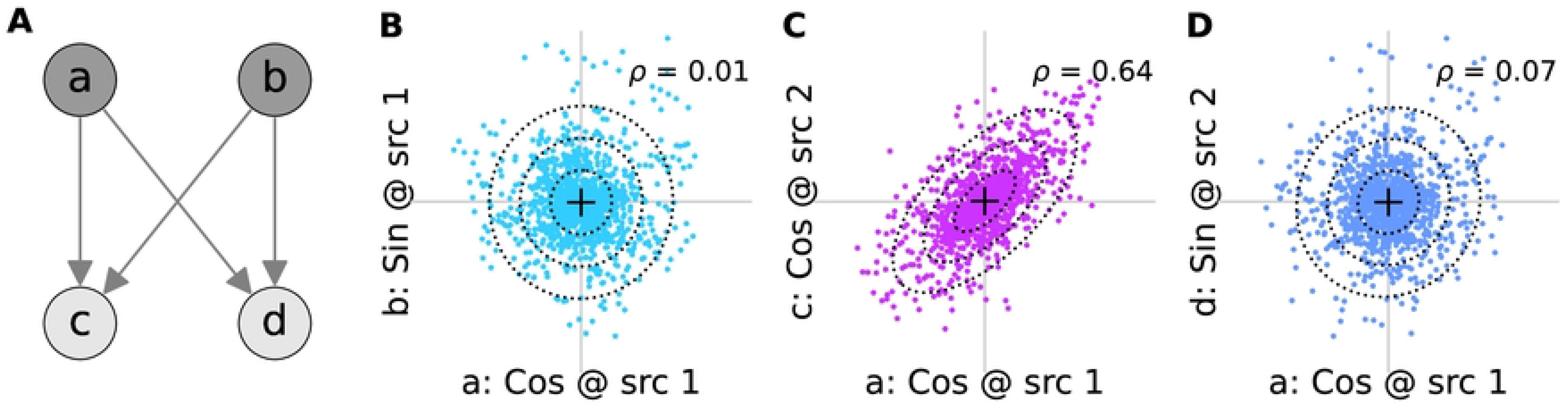
Decomposition of phase relationships. **(A)** Knowing the cosine (*a*) and sine (*b*) coefficients at one source can inform about the corresponding coefficients (*c* and *d*, respectively) at another. The lack of arrows between *a* and *b*, and between *c* and *d*, indicates their marginal independence. **(B)** Cosine (*a*) vs. sine (*b*) coefficients from the same source at 5 Hz, pooled across all sources. **(C)** Cosine coefficients from one source (*a*) vs. those (*c*) from the closest source in the positive vertical direction, pooled across all such pairs. **(D)** As in panel C but plotted against the sine coefficient from the neighbouring source. The Pearson correlation of the coefficients in each panel is indicated with *ρ*.

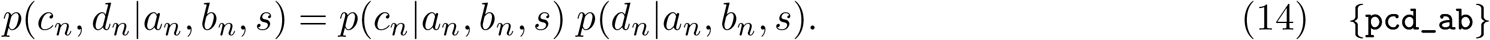

To specify a form for the conditional distributions on the right-hand side of Eqn 14, we must determine the uncertainty that remains about the coefficients *c*_*n*_ and *d*_*n*_ of the component waveform at the second source when we know the corresponding coefficients *a*_*n*_ and *b*_*n*_ at the first. To do so, we expressed the component waveform at the second source as a scaled and phase-shifted version of the first, plus a residual. Doing so, and making our final assumption,
4. The conditional distribution *p*(*c*_*n*_, *d*_*n*_|*a*_*n*_, *b*_*n*_, *s*) of coefficients at one source (*c*_*n*_ and *d*_*n*_) given those at another (*a*_*n*_ and *b*_*n*_), is bivariate Gaussian,
we concluded that the conditional distribution of the individual coefficients is

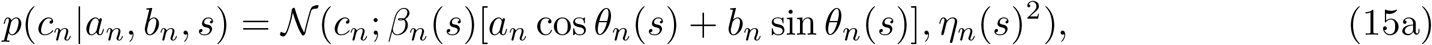

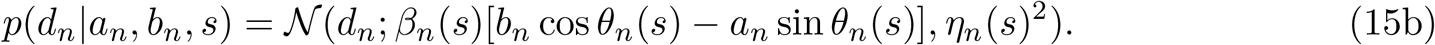

Here *β*_*n*_(*s*) and the *θ*_*n*_(*s*) are the best-fit scaling and phase-shifts, and *η*_*n*_(*s*)^2^ is the variance of the residual (see Methods Sec 5.3.2).

Substituting these equations into Eqn 14 and combining with Eqn 12 we arrive at the form of the joint distribution of the coefficients

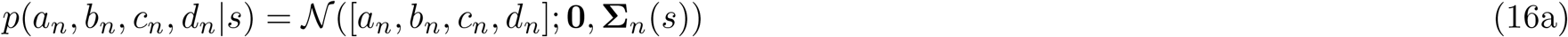

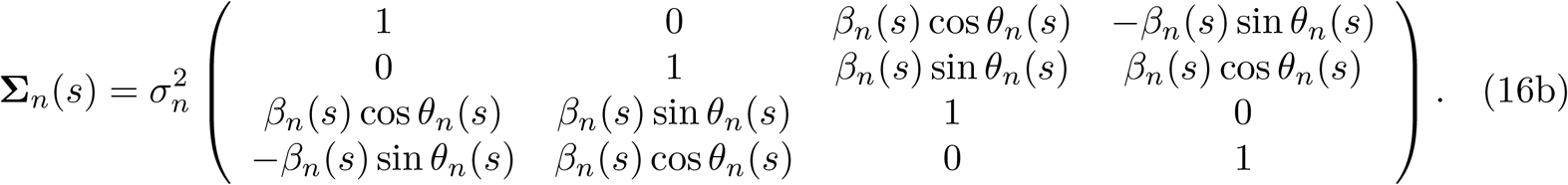

Note that by the location independence assumption, the marginal variance 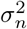of the coefficients does not depend on *s*.

The covariance in Eqn 16 specifies two types of relationship between the coefficients from the two sources. Those that are ‘in-phase’ relate coefficients of the same type, for example the cosine coefficients *a*_*n*_ and *c*_*n*_. Those that are ‘out-of-phase’ or ‘quadrature’ relate coefficients of different types, for example the cosine coefficient *a*_*n*_ at the first source and the sine coefficient *d*_*n*_ at the second. Determining the spatial information that correlations provide requires incorporating both types of relationship by using the full set of covariances specified in Eqn 16.

However, for our data the out-of-phase contribution is very small, and the in-phase relationship dominates. This can be seen, for example, in the high in-phase correlation of the data in Fig 5C vs. the near zero correlation of the out-of-phase data in Fig 5D (see also Fig S10). Therefore, it will be convenient to focus on the marginal distribution relating only the in-phase coefficients. For the pair of cosine coefficients this is

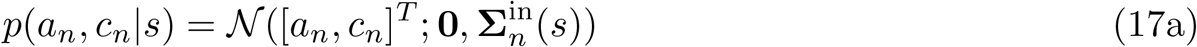

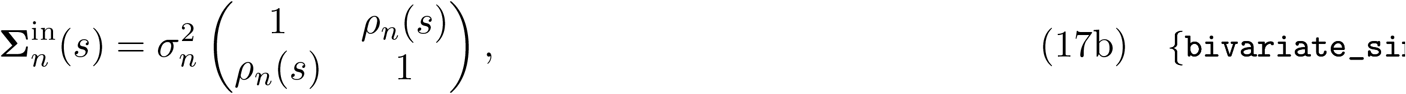

where we’ve defined the in-phase correlation

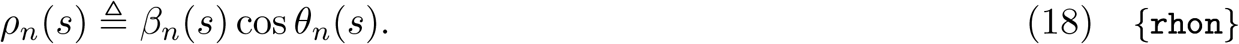

The relationship relating the pair of sine coefficients *b*_*n*_ and *d*_*n*_ has the same form as Eqn 17. Since the off-diagonal element of 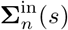 is the covariance of pairs of sine or cosine coefficients, we can use Eqn 3 to relate the in-phase correlations to the average component correlations as

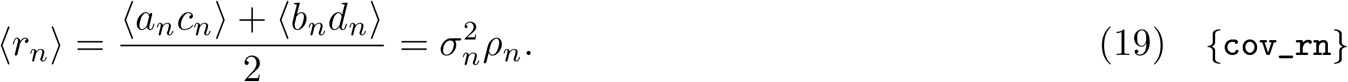

The relationship of the out-of-phase coefficients can be described in a similar way. For example,

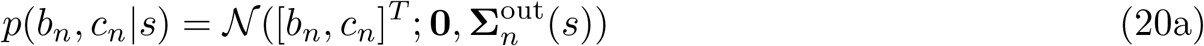

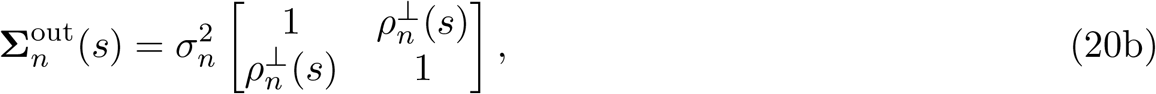

where the out-of-phase correlation is defined as

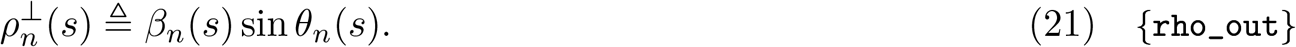

The distribution *p*(*a*_*n*_, *d*_*n*_|*s*) has the same form, but with negative the covariance between the two variables. In Fig 6 we have plotted examples of the joint distribution of in-phase (panels A-D) and out-of-phase (panels E-H) coefficients. These plots confirm that the out-of-phase correlations, while not zero, are much smaller than the in-phase correlations.

**Fig 6.**
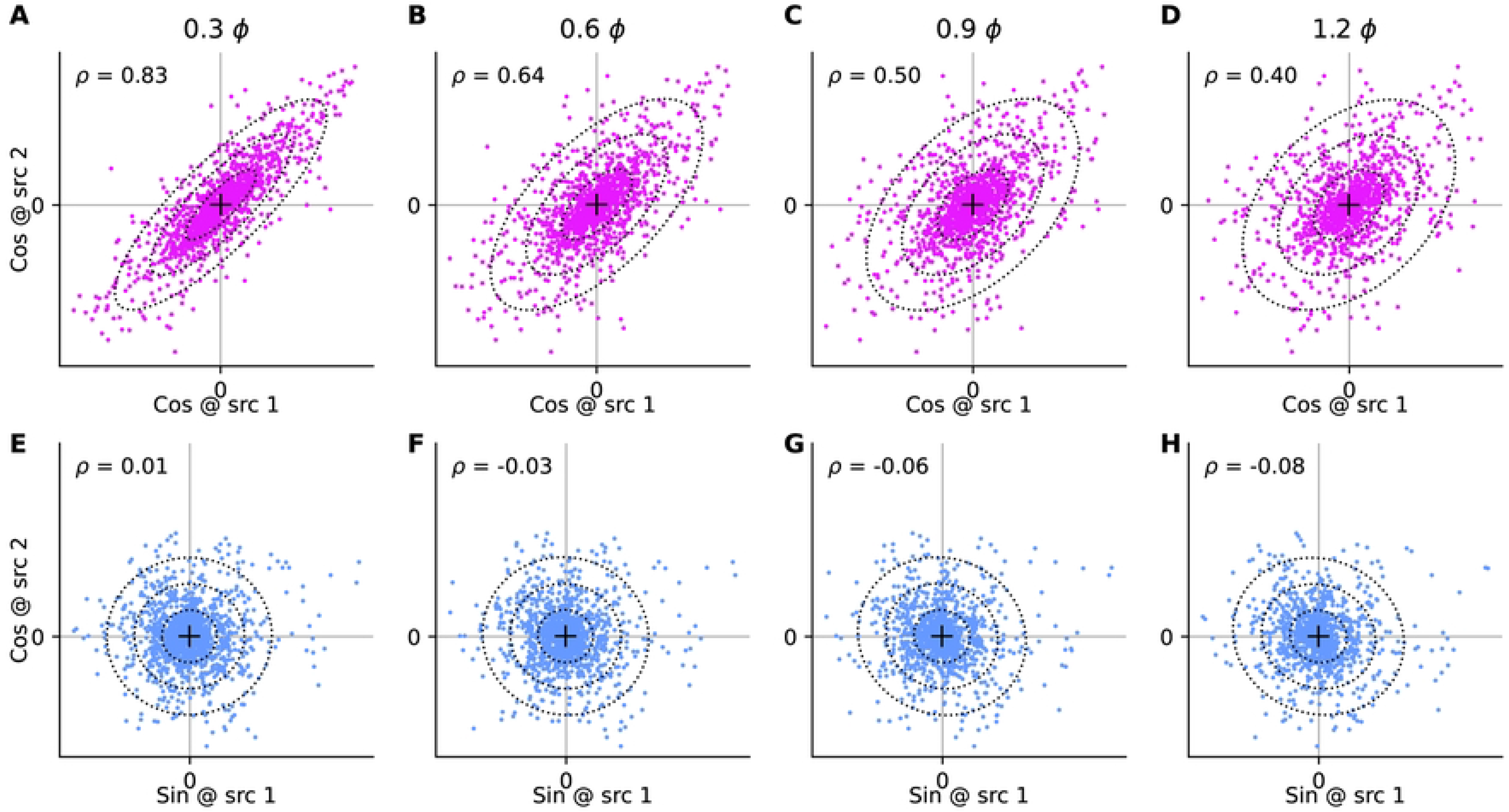
Fourier coefficients vs. distance. Distribution of the indicated coefficients for the 5 Hz component of the plume from one source vs. the indicated coefficient computed for the second source, computed for all pairs of sources separated by the intersource distance stated above panels A-D. Crosses indicate the mean of each dataset; ellipses indicate first three standard deviations of bivariate normal fit to the data; Pearson correlations (*ρ*) of the data in each plot are indicated in the top-left corner. **(A-D)** Cosine coefficient at source 1 vs. cosine coefficient at source 2. **(E-H)** Sine coefficient at source 1 vs. cosine coefficient at source 2.

Armed with the joint distribution of Fourier coefficients given intersource distance specified in Eqn 16, we can return to Eqn 5 and derive the distribution of correlations to be the asymmetric Laplacian (see Methods Sec 5.3.3)

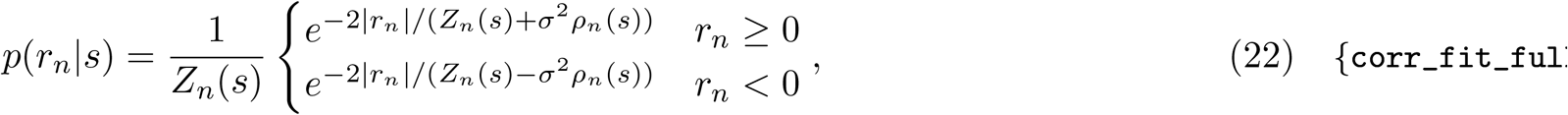

where the normalizing constant is defined as

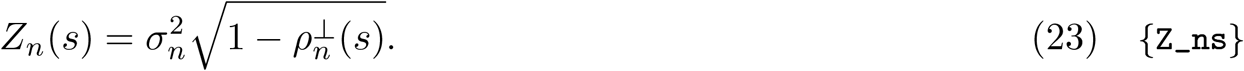

#### A simplifying assumption

Because the out-of-phase correlations of our data are typically very close to zero, in what follows we will assume that these correlations are zero. In that case, 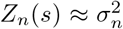 and Eqn 22 simplifies to

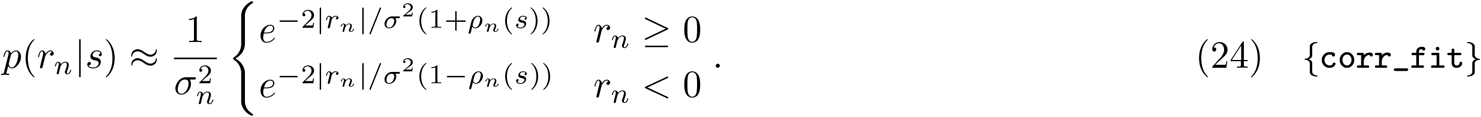

To evaluate the agreement of the asymmetric Laplacian in Eqn 24 with the observed distribution of correlations we compare their cumulative distribution functions (CDFs). In Fig 7A-C we make the comparison for the 5 Hz correlation data at three different intersource distances. We quantified the agreement between the CDFs as 1 minus the largest absolute different between them. A value of 1 would indicate perfect agreement, while, a value of 0, the smallest possible would indicate non-overlapping distributions. A heatmap of the match over the full range of intersource distances and frequencies (Fig 7G) reveals a match of ∼ 0.7 and higher over most of this range.

**Fig 7.**
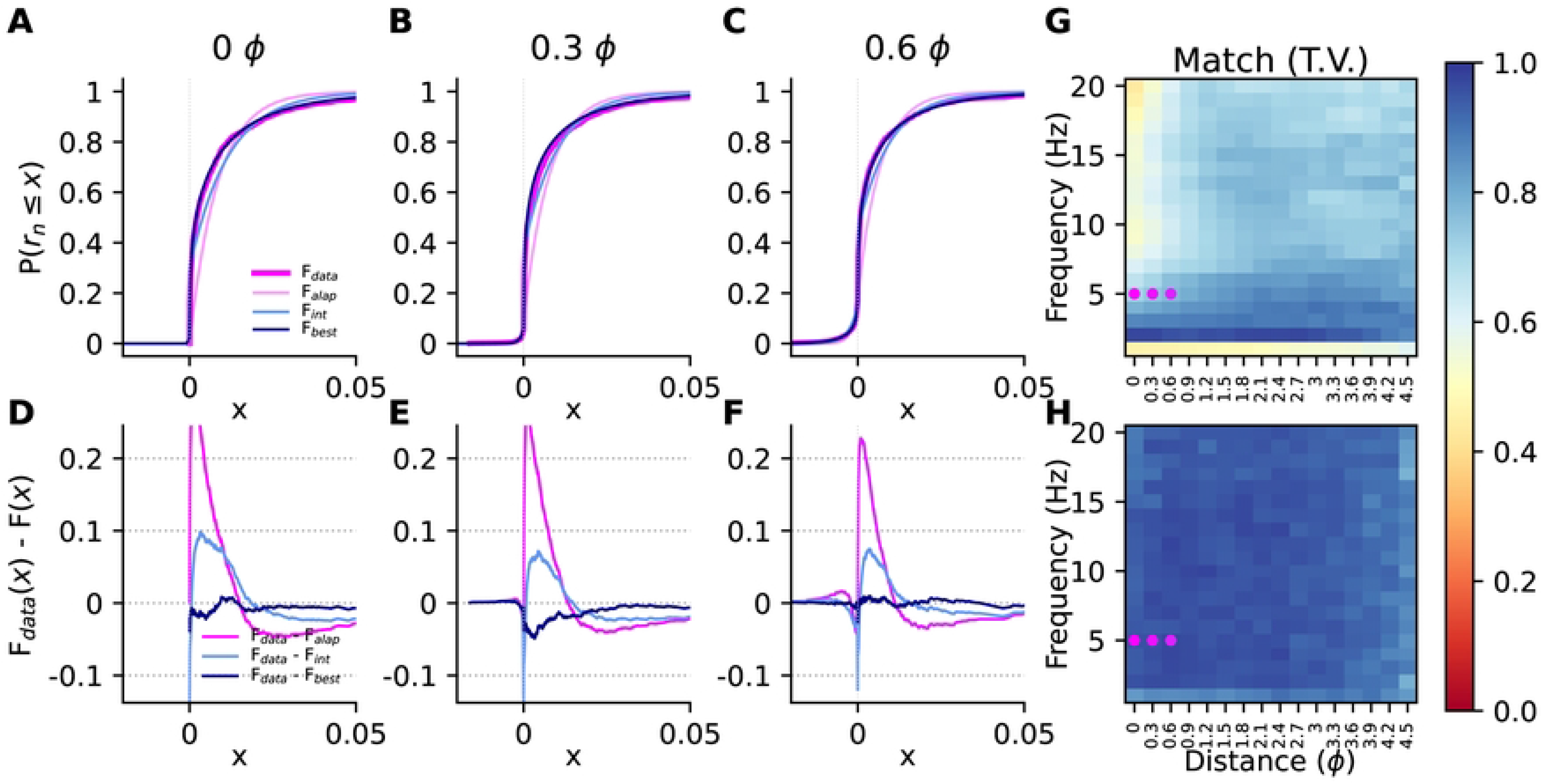
Modeling the distribution of observed correlations. **(A–C)** Examples of observed (F_*data*_, thick pink) and predicted cumulative distribution function for the 5 Hz component of correlations, for three different intersource distances (indicated above each panel). Predictions were for the asymmetric Laplacian distributions (F_*alap*_, pink) in Eqn 24, asymmetric Laplacian with intermittency (F_*int*_, light blue), and the best-fitting model when testing both intermittent and non-intermittent versions of the asymmetric Laplacian, Gamma, and generalized inverse Gaussian (F_*best*_, navy). The parameters of F_*alap*_ were derived from Gaussian fits to the distributions of Fourier coefficients, the parameters for the remaining fits were found by directly fitting the correlation data. **(D–F)** Absolute difference between the predicted and observed cumulative distribution functions in the corresponding panels in the first row. Data in A-F may be optimistic because fits were computed using some (though not all) of the data they are being qualitatively evaluated against in those panels. See panel H for performance on held-out data. **(G)** Fit quality for the asymmetric Laplacian model, measured as 1 minus the largest absolute difference between the predicted and observed CDFs, computed for all distances and frequencies listed. Higher values indicated better fits. Values may be optimistic because parameters were both trained and evaluated on the same (full) dataset. Coloured dots correspond to data points shown in the left three panels. **(H)** As in panel G, but for the best fitting model when testing both intermittent and non-intermittent versions of the asymmetric Laplacian, Gamma, and generalized inverse Gaussian, fitted directly to the correlations, and evaluated on unseen data. See Methods Sec 5.2.2.

#### 2.2.1 Improving the model of correlations

Although there is good qualitative agreement between the data distributions and the corresponding asymmetric Laplacians, the plots in Fig 7A-C also suggest that there are systematic differences between the two. In Fig 7D-F we have plotted the differences between the data distribution and the asymmetric Laplacian fits. These plots have similar shapes, consisting of a narrow positive lobe followed by a broader negative lobe, eventually decaying to zero as both CDFs approach 1. This reveals that the data had more correlation values concentrated near zero, and correspondingly fewer large correlation values, than the asymmetric Laplacians.

To achieve a better fit to the observed distribution of correlations we elaborated on the asymmetric Laplacian model to better capture the small correlations. We reasoned that since turbulent flows produce intermittent signals [41], a portion of the very small correlations may be attributed to the absence of actual correlations and attributable instead to noise. Similar to previous approaches used to incorporate intermittency into single-source concentration distribution models [32], our correlation model captures intermittency by introducing a binary random variable *z* that determined whether the observation *y*_*n*_ was of a correlation *r*_*n*_, or noise *w*. That is,

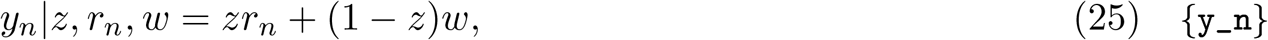

so that when the masking variable *z* = 1, we observe a ‘true’ correlation *r*_*n*_, and when *z* = 0, we observe an instance of noise instead. We assume that the masking variable *z* obeys a Bernoulli distribution,

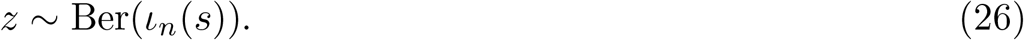

The intermittency parameter *ι*_*n*_(*s*) ∈ [0, 1] determines the fraction of observations we deem to be true correlations, and can depend both on the harmonic *n* and the intersource distance *s*. The true correlations *r*_*n*_ are drawn from the asymmetric Laplacian distribution of correlations in Eqn 24,

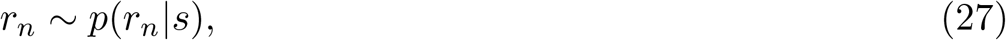

while the noisy observations *w* come from a zero-mean Gaussian

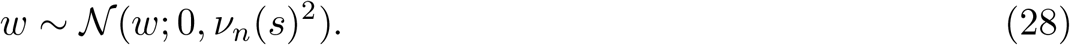

Like the intermittency parameter, the variance 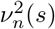 of the noise can depend on both the harmonic and intersource distance. The predicted distribution of correlations for our ‘intermittent asymmetric Laplacian’ model is then a linear combination of the ‘true’ correlation distribution *p*(*r*_*n*_|*s*) and the noise distribution determined by the intermittency parameter *ι*_*n*_(*s*)

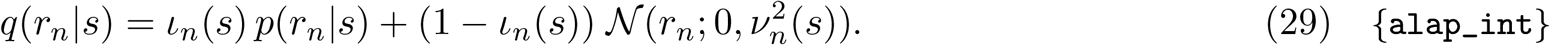

Setting *ι*_*n*_(*s*) to 1 corresponds to our initial asymmetric Laplacian model, while setting it zero represents a fully intermittent case where all correlations are in fact noise. The amount of intermittency and the variance of the noise were determined from the data (see Methods, Sec 5.2.2).

To demonstrate the effect of including intermittency in our model we have added the CDFs for the distributions in Eqn 29 fitted to the data in Fig 7A-C to the corresponding panels in those plots (light blue traces), revealing an improved qualitative agreement. In Fig 7D-F we have added the deviations between the data CDFs and those of the new model that includes intermittency. The panels show that both the positive and negative deviations have been reduced, as expected, but not eliminated.

The fact that both positive and negative deviations remain in Fig 7D-F indicates that the data have more small values than even augmentation with intermittency can capture. Therefore, we extended our model by replacing the Exponential distributions in Eqn 24 that constitute the positive and negative halves of the asymmetric Laplacian with distributions that had higher densities near zero, but that included the Exponential as special cases. One candidate is the Gamma distribution, *p*(*r*) ∝ *r*^*k*−1^*e*^*r/λ*^. Just like the Exponential distribution, the Gamma distribution has a scale parameter, *λ* for the positive correlations above. In addition, it introduces a shape parameter, *k >* 0. When *k* = 1, the Gamma distribution reduces to the Exponential distribution. When *k <* 1, the density approaches infinity as *r*→ 0. This singularity at 0 aids in modeling the large number of low correlation values we observe. Analogously to Eqn 24, we ‘sandwich’ two Gamma distributions together so that we can also cover negative values (see also Methods Sec 5.2.2), and arrive at our correlation distribution

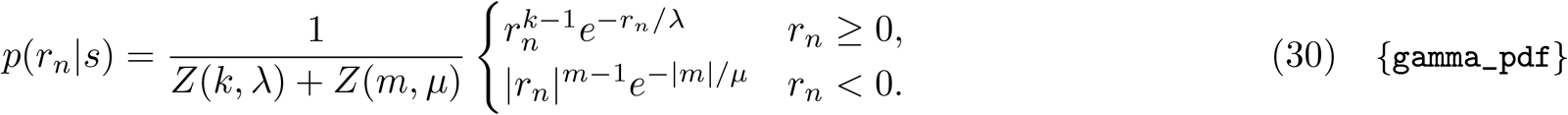

Here *Z* is the normalizing function of the Gamma distribution, *k* and *λ* are the shape and scale parameters for the positive correlations, and *m* and *μ* are the corresponding parameters for the negative correlations. We have omitted the dependence of these parameters on the harmonic *n* and the intersource distance *s* for clarity.

Another probability distribution, which includes the Gamma (and therefore the Exponential) as a special case, is the generalized inverse Gaussian. The parameterization of this distribution that we use gives the probability density function *p*(*r*) ∝ *r*^*k*−1^*e*^−*α*(*r/λ*+*λ/r*)*/*2^. This distribution introduces a further shape parameter *α >* 0 and a term proportional to 1*/r* in the argument of the exponential that provide additional flexibility in modeling small observations. The scale of the distribution can be taken to be the length constant of the *r* term in the exponential, 2*λ/α*. Keeping this constant while reducing *αλ* towards zero, one approaches the Gamma distribution. As before, we sandwich two such distributions together to arrive at our distribution of correlations,

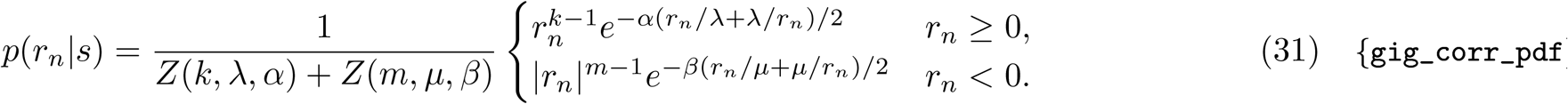

Here *Z* is the normalizing function of the generalized inverse Gaussian distribution, *k, λ*, and *α* are the parameters for the positive correlations, and *k* and *μ*and *β* are the corresponding parameters for the negative correlations. As before, we have omitted the dependence of these parameters on the harmonic *n* and the intersource distance *s* for clarity.

Our elaborated models of the correlations were thus of the same form as Eqn 29, with *p*(*r*_*n*_ | *s*) set to either asymmetric Laplacian, or sandwiched versions of the Gamma, or generalized inverse Gaussian distributions. We fit both intermittent and non-intermittent versions of these models to the correlations at each frequency and intersource separation separately. Examples of such fits are shown in Fig 7, showing improved agreement with the observed correlations. The performance of the best-fitting models on unseen data over the full range of frequencies an intersource distances are shown Fig 7H, demonstrating a very good fit over the entire range.

### 2.3 Computing the Fisher information

Having now determined expressions for the distribution of correlations given intersource distance, we can use Eqn 4 to determine the Fisher information. Evaluating that expression requires not just a form for the correlation distributions themselves, but also for how they change with intersource distance. These changes in turn are determined by how the various parameters of our correlation model change with intersource distance.

The three models we have considered, the asymmetric Laplacian, Gamma and generalized inverse Gamma, have 2, 4, and 6 parameters, respectively. The intermittent version of each model requires an additional two parameters to capture the intermittency. To avoid the complexity of modeling how large numbers of parameters change with intersource distance, we will use our simplest model, the non-intermittent asymmetric Laplacian of Eqn 22, to estimate Fisher information.

The general expression using Eqn 22 is complex but simplifies significantly when the out-of-phase correlation is zero, which is approximately the case for our data. Therefore in what follows, we will use the simplified expressions. The Fisher information is then (see Methods Sec 5.3.4)

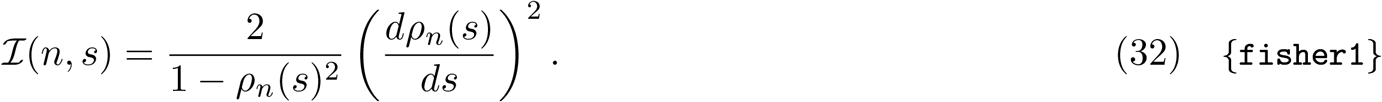

As basic checks of this expression we observe that

- It’s non-negative since 0 ≤ *ρ*_*n*_(*s*)^2^ ≤ 1;
- It depends on how the correlations change with distance, via the *dρ*_*n*_(*s*)*/ds* term, so that when this dependence is zero, there is no information in the correlations, as expected;
- For a fixed amount of distance dependence *dρ*_*n*_(*s*)*/ds*, information is least when there is no correlation (*ρ*_*n*_(*s*) = 0).

Armed with Eqn 32, we still require the distance dependence of correlations *ρ*_*n*_(*s*) to compute the Fisher information. To motivate a parametric form for this dependence, we require that at the very least it should peak at an intersource distance of zero and decrease with distance. A simple parametric form that meets this requirement is exponential decay with distance *s*.

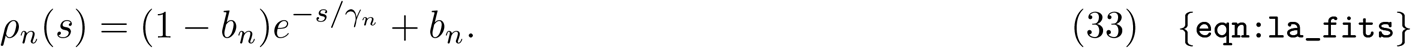

The two parameters of the model are the length scale of the decay *γ*_*n*_, and the constant offset *b*_*n*_ that determines the value at very large intersource distances. In Fig 8 we demonstrate the fit of this parametric form to some example correlation data.

**Fig 8.**
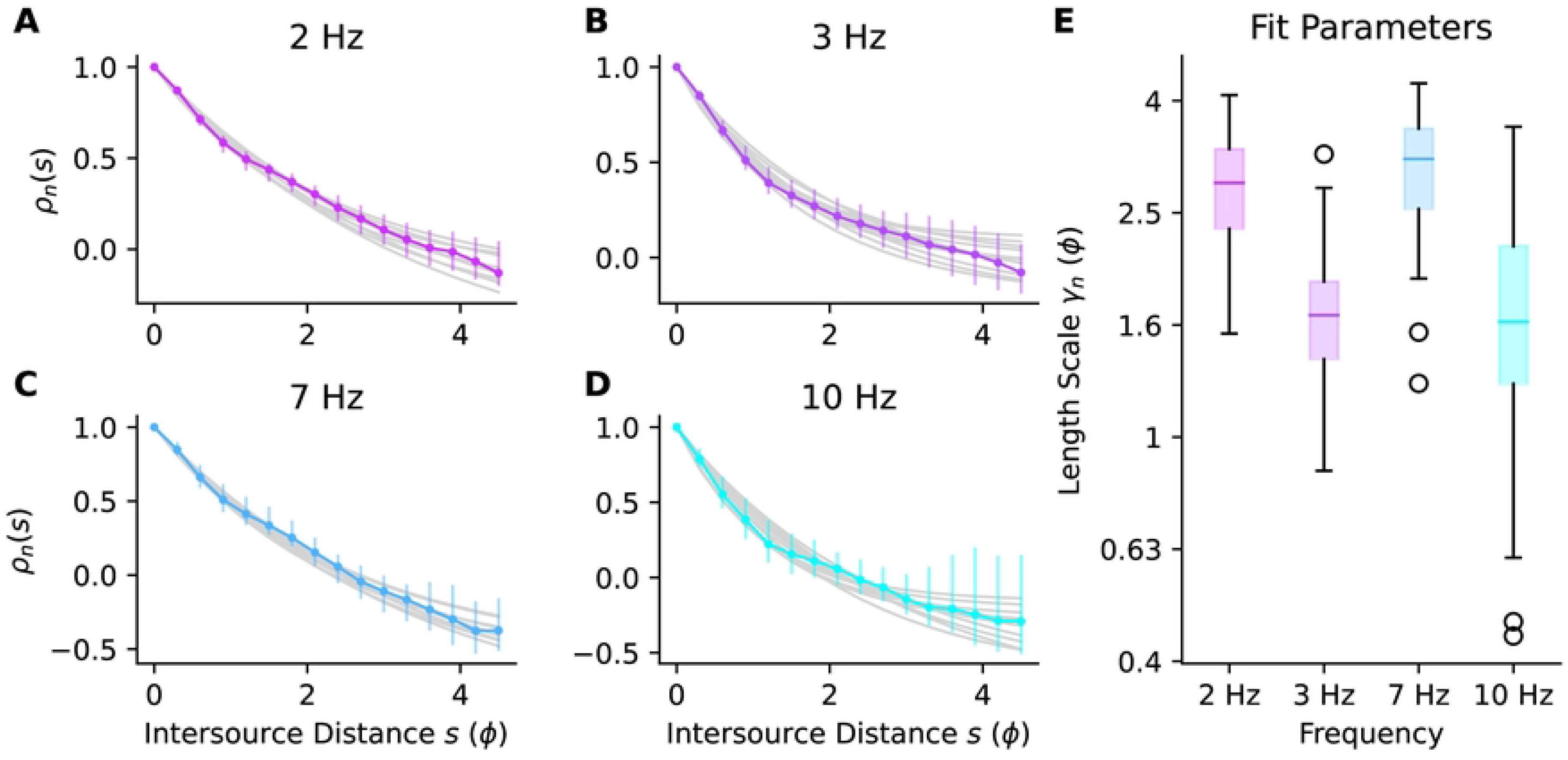
Parametric fits to *ρ*_*n*_(*s*). **(A–D)** Bootstrap medians (coloured points) and 5–95th percentiles (coloured bars) of *ρ*_*n*_(*s*) at each intersource distance, and a few example fits using bootstrap parameters (gray traces), for a few example frequencies. **(E)** Length scale parameter (on a logarithmic axis) for the fits to the data in (A–D). Boxes indicate inter-quartile range (IQR), central lines are medians, whiskers extend to last data points within 1.5 IQR, circles are outliers.

Interestingly, when we examined the decay of correlation with distance, we observed that higher frequencies decayed faster than lower frequencies (Fig 9A). We plotted the length constants of the generalized exponential fits to the decays against frequency (Fig 9B) and observed that they decreased at about one pitch every 10 Hz (Fig 9E). According to Eqn 32, Fisher information depends on the rate of change of correlations with intersource distance. Therefore, the differences in decay rates that we observed suggest that the amount of information carried by correlations at different frequencies varies. As a comparison, we generated surrogate data with a similar power spectrum to our simulated plumes, but for which all frequencies were equally informative (see Methods Sec 5.2.7). We observed very little change in length constants with frequency in this surrogate data (Fig 9C,D,E).

**Fig 9.**
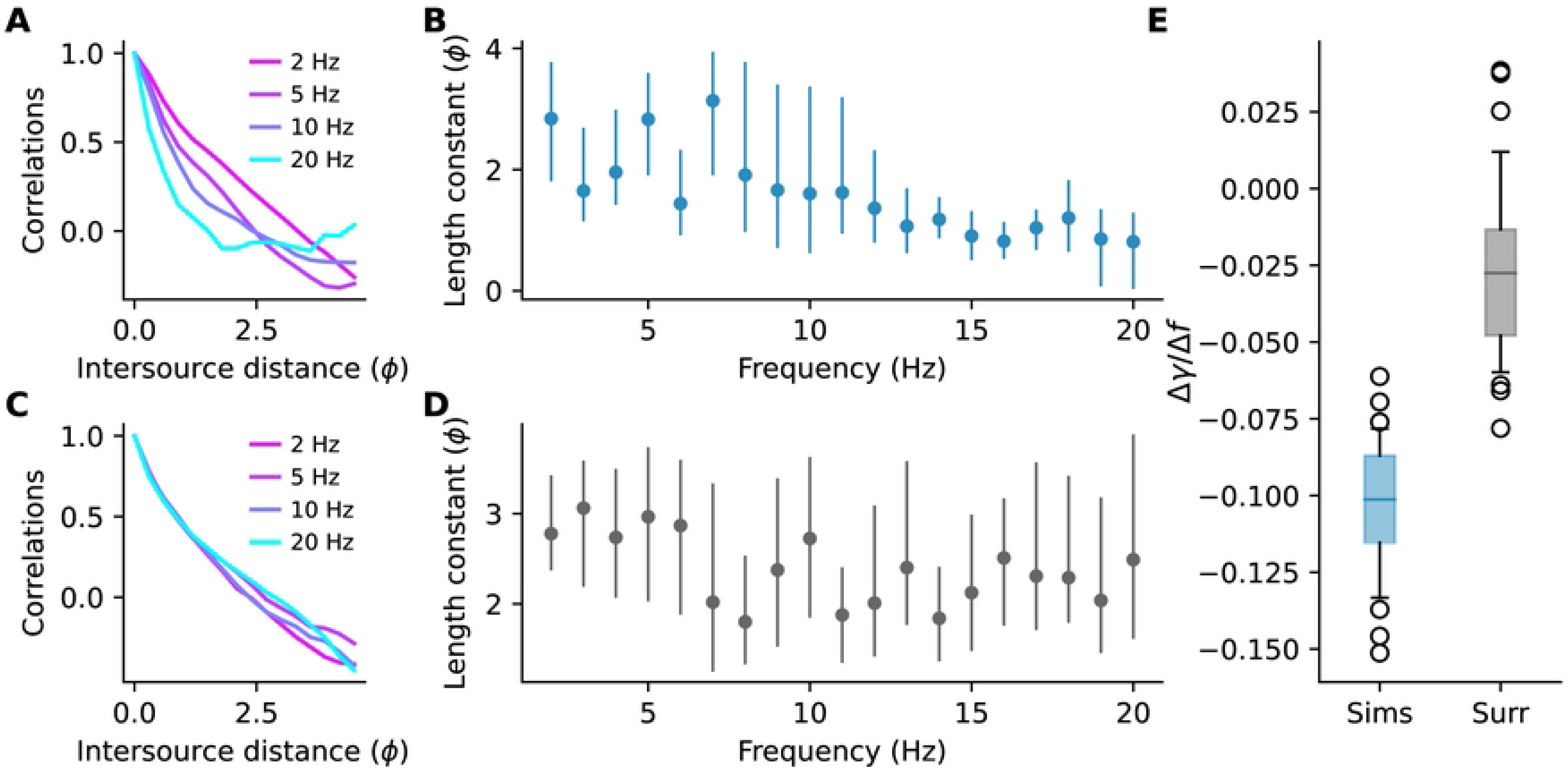
The decay of correlations with frequency. **(A)** Normalized correlations at different frequencies. **(B)** Length constants *γ* of exponential fits to the decay of correlations for different frequency components. Dots and lines are the bootstrap median and 5th-95th percentiles, respectively **(C**,**D)** As in panels A and B but for surrogate data where all frequencies are equally informative. **(E)** Bootstrap distribution of coefficients when regressing the length constants in **(B**,**D)** on frequency. Negative values indicate length constants shorten with frequency. Central lines are medians, boxes are interquartile range, whiskers are [5th-95th] percentiles, circles are outliers. The simulation data (but not the surrogate data) had very large time constant at 1 Hz so only the data from 2 - 20 Hz are shown above and used in the analysis.

Given the parametric form Eqn 33 of the distance dependence we derived an analytic expression for the Fisher information. To simplify the expression we first normalized distance *s* by the length-scale parameter *γ*_*n*_ and defined

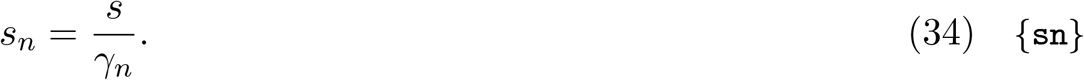

In Methods Sec 5.3.4 we show that the Fisher information about intersource distance provided by the *n*’th harmonic component of correlations can be expressed in terms of normalized distance *s*_*n*_ as

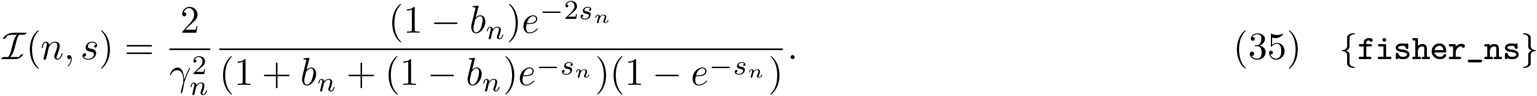

In Fig 10A we’ve plotted the bootstrap median and 5th-95th percentiles of Fisher information at a range of distances for a few example harmonics. The heatmap in Fig 10B shows the Fisher information computed using all the data (i.e. not bootstrapped) at a broad range of frequencies and intersource distances.

**Fig 10.**
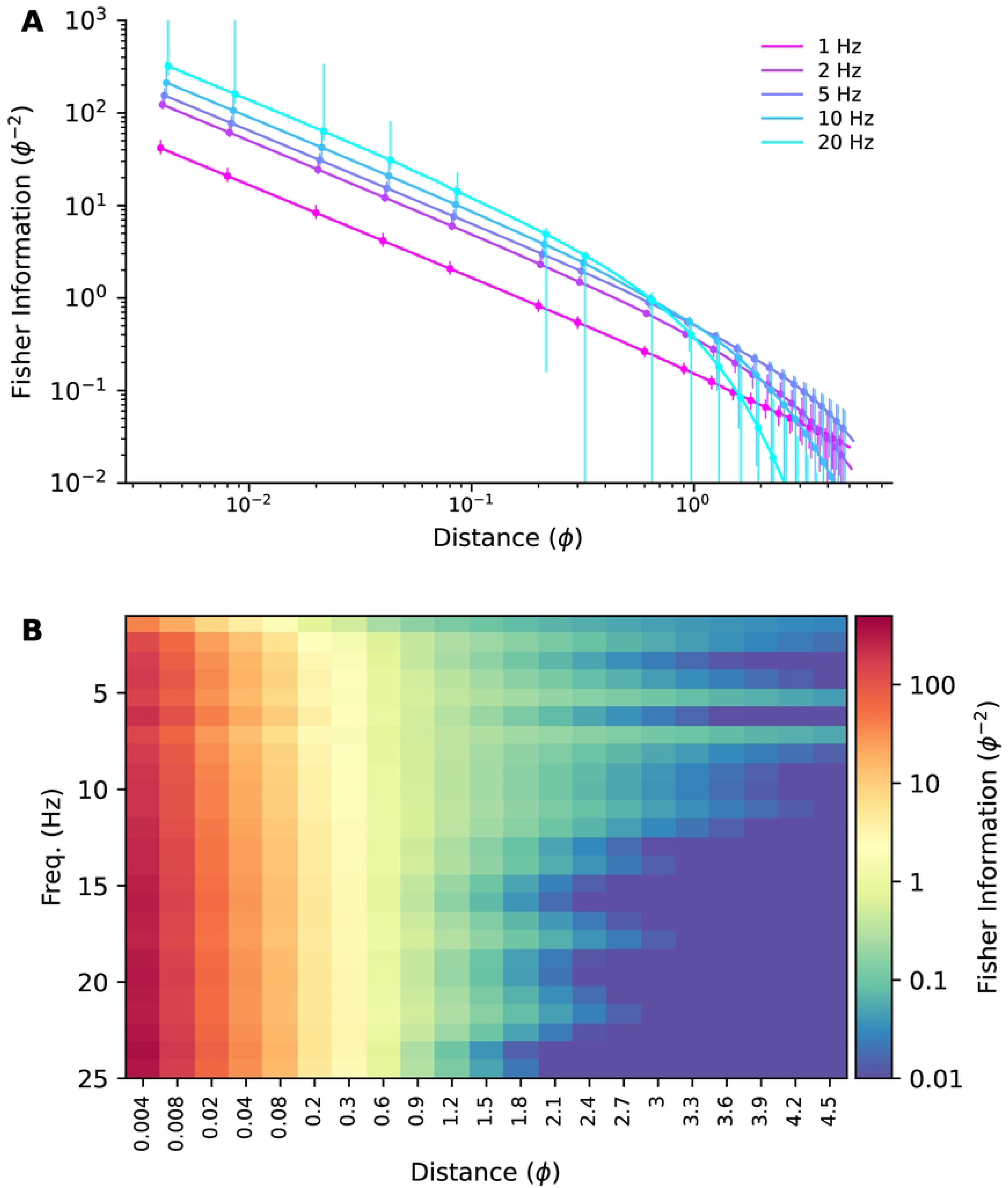
Fisher information vs. intersource distance. **(A)** Bootstrap medians (points) and 5–95th percentiles (bars) of Fisher information vs. intersource distance for five example frequencies. **(B)** Fisher information heatmap, computed using all the data (i.e. not bootstrapped). Note the nonlinear scaling of the x-axis.

#### 2.3.1 Which frequencies are more informative?

The Fisher information heatmaps for our CFD plumes suggest that high frequencies are more informative when sources are close together. To see what such heatmaps would look like when the ground-truth distribution of information over frequencies is known, we generated surrogate plume datasets with similar power spectra to our CFD plumes, but with different distributions of information across frequencies (see Methods Sec 5.2.7). In the first such dataset, all frequencies were set to be equally informative. In Fig 11B we have plotted the Fisher information heatmap for this dataset. The information content varies with intersource distance, but is homogeneous with frequency, as expected, and unlike that of our CFD plumes (reproduced in Fig 11A).

**Fig 11.**
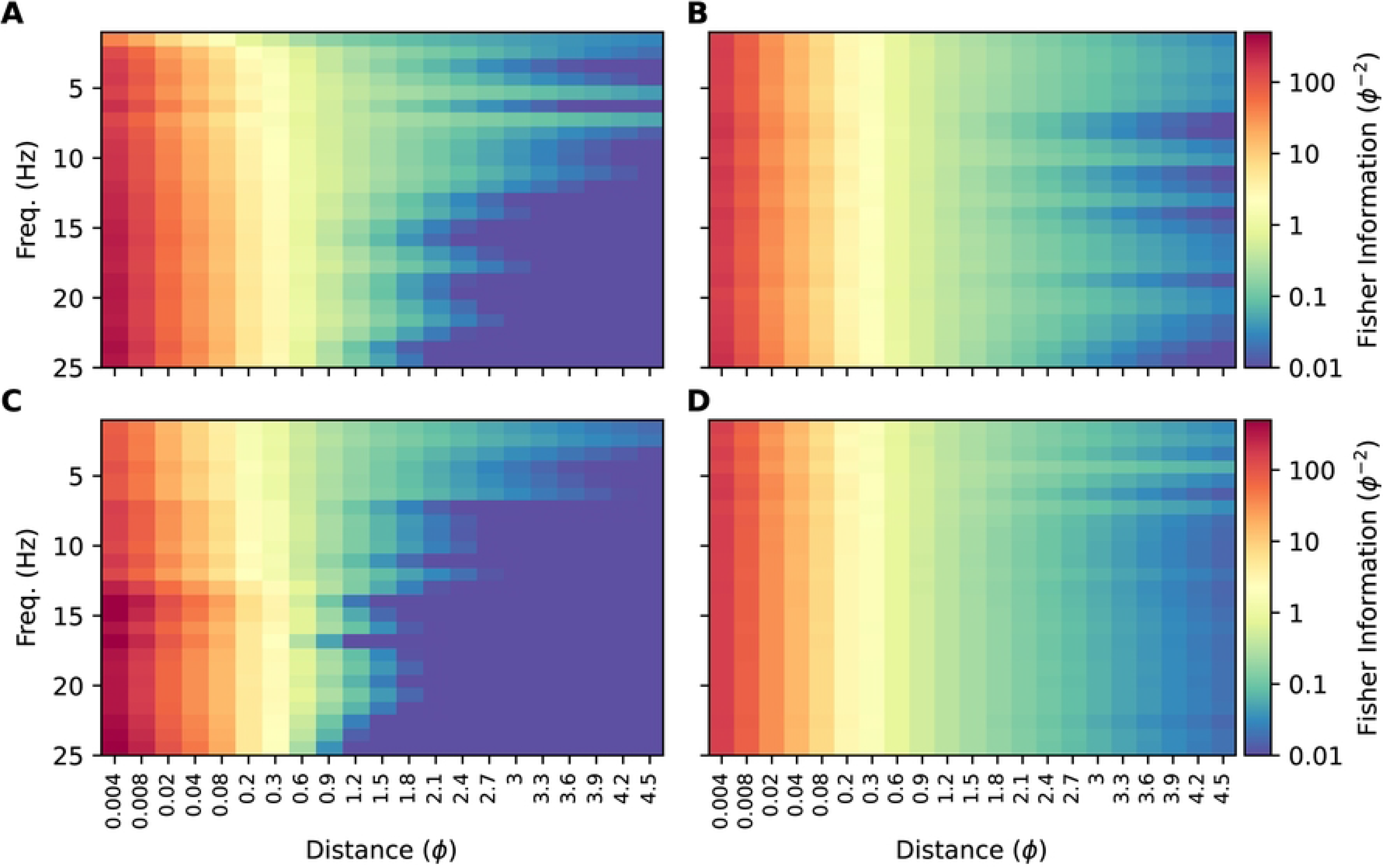
Fisher information for simulations and surrogate data. **(A)** Fisher information heatmap for our simulated plumes (reproduced from Fig 10B). **(B)** Heatmap for surrogate data with power spectrum similar to our simulated data and where all frequencies are equally informative. **(C)** Heatmap for surrogate data with power spectrum similar to our simulated data and where frequencies *>* 12.5 Hz have the same, higher value of Fisher information than the lower frequencies. **(D)** Fisher information for our simulated plumes when using a 1-second boxcar window.

Next, we generated a second surrogate dataset in which correlations decayed with intersource distance six times faster for the higher half of the frequency range than the lower half. This should result in the high frequencies being more informative at small intersource distances. At large intersource distances, high frequency correlations will have decayed to zero due to their fast decay rate, yielding little information. Low frequencies will not have decayed completely to zero and will remain informative. Thus we expected to see high frequencies being more informative at small intersource distances, and low frequencies more informative at large intersource distances. This is indeed what we saw when we plotted the Fisher information heatmap for this dataset in Fig 11C.

The similarity of the information heatmap for our CFD plumes (Fig 11A) to that of the surrogate dataset where high-frequencies were *a priori* more informative (Fig 11C), and its dissimilarity to the heatmap of the dataset where all frequencies were *a priori* equally informative, supports the conclusion that for our CFD plumes high frequencies are more informative when sources are close together.

To emphasize the importance of windowing to these results, in Fig 11D we plot the Fisher information heatmap for our CFD plumes, but when analyzed with a 1-second boxcar window. The boxcar window, also known as a rectangular window, weights all samples equally. The poor spectral leakage properties of this window coupled with the power-law power spectrum of our data (Fig S7) results in the low frequency information masking that of the higher frequencies. This then results in an heatmap where information appears to be homogeneously distributed across frequencies, similar to Fig 11B.

From Eqn 32, Fisher information depends on the rate at which correlations change with distance. The fact that correlations decay faster for high frequencies than for low frequencies means that when sources are closer together, the correlations at high frequencies will be more informative. This is shown in the left-most panel of Fig 12A, where we have plotted the information available at each frequency for an intersource distance of ∼ 0.1 pitch. At that intersource distance information shows a positive trend with frequency. As sources move farther apart, high frequency correlations will have decayed, eventually changing at the same rate with distance as the slower decaying low frequency correlations. This will result in all frequencies being similarly informative, as shown in the middle panel of Fig 12A for an intersource separation of ∼ 1 pitch. As sources move even farther apart, the lower frequencies will become more informative, as shown in the right panel of Fig 12A for an intersource separation of ∼ 2 pitches. The overall change in the trend as intersource distance increases is shown in Fig 12C. For surrogate data in which all frequencies are equally informative, trends in information with frequency are minimal at all intersource separations; see Fig 12B,C. Note that the exponential decay in correlations with frequency means that regardless of which frequencies are more informative, the absolute amount of information available will decrease with intersource distance. This is because information is derived from the rate of change of correlation with intersource distance, and this decreases as sources move farther apart. This effect is shown by the decrease in the down-scaling applied to the data in the panels of Fig 12A and B.

The analyses above relied on the use of windows that reduced spectral leakage when the amount of power in different frequency bands is different, as is the case in real plumes and in our simulations. By using the Hann window to reduce leakage we were able to distinguish the different rates at which correlations decay in different frequency bands, and therefore the different amounts of information that they contain. As we show in Fig S1 and Fig S2 these effects were not limited to the 1-second Hann window used above, and we also observed them when we used other windows that similarly reduced spectral leakage.

## 3 Discussion

### 3.1 Quantifying information in plume correlations

The concentration timeseries from two odour sources measured at a downwind location tend to become more correlated when the sources move closer together. Therefore, correlations contain information about the spatial separations of the odour sources that generated them. Correlations can be decomposed as the sum of component correlations at different frequencies. Recently it was shown that mice can discriminate correlated vs. anti-correlated concentration timeseries at up to 40 Hz [36]. Could this high bandwidth sensitivity be caused by additional spatial information those frequencies might carry? Motivated by this question we set out to quantify the information contained in the component correlations about the spatial location of odour sources, to determine whether high frequencies contain more spatial information than lower frequencies.

Our approach is based on parametric modeling of how component correlations decay with intersource distance. We first express the component correlation at each frequency in terms of the corresponding Fourier decomposition coefficients of the odour timeseries being correlated. We then fit parametric models to these correlations and how they change with intersource distance. Using these parametric models we then derived a closed form expression for the Fisher information contained in the component correlations about the intersource distance in terms of the correlation of the Fourier coefficients and how these change with intersource distance (Eqn 32).

We applied our approach to two computational fluid dynamics simulations of two-dimensional grid turbulence to determine which frequency bands were most informative about relative source locations. To verify our approach we also applied it to several surrogate datasets which we constructed to contain different patterns of spatial information in their component correlations. Analyzing our data using a 1-second Hann window to reduce spectral leakage, we observed that high frequency correlations decayed faster than those at low frequencies (Fig 9). This meant that high frequencies were more informative when sources were less than ∼ 1 pitch apart, and low frequencies when sources were farther apart than this Fig 12. We saw this effect for other similar leakage-reducing windows (Fig S1 and Fig S2), and not in surrogate data in which all frequencies were equally informative.

**Fig 12.**
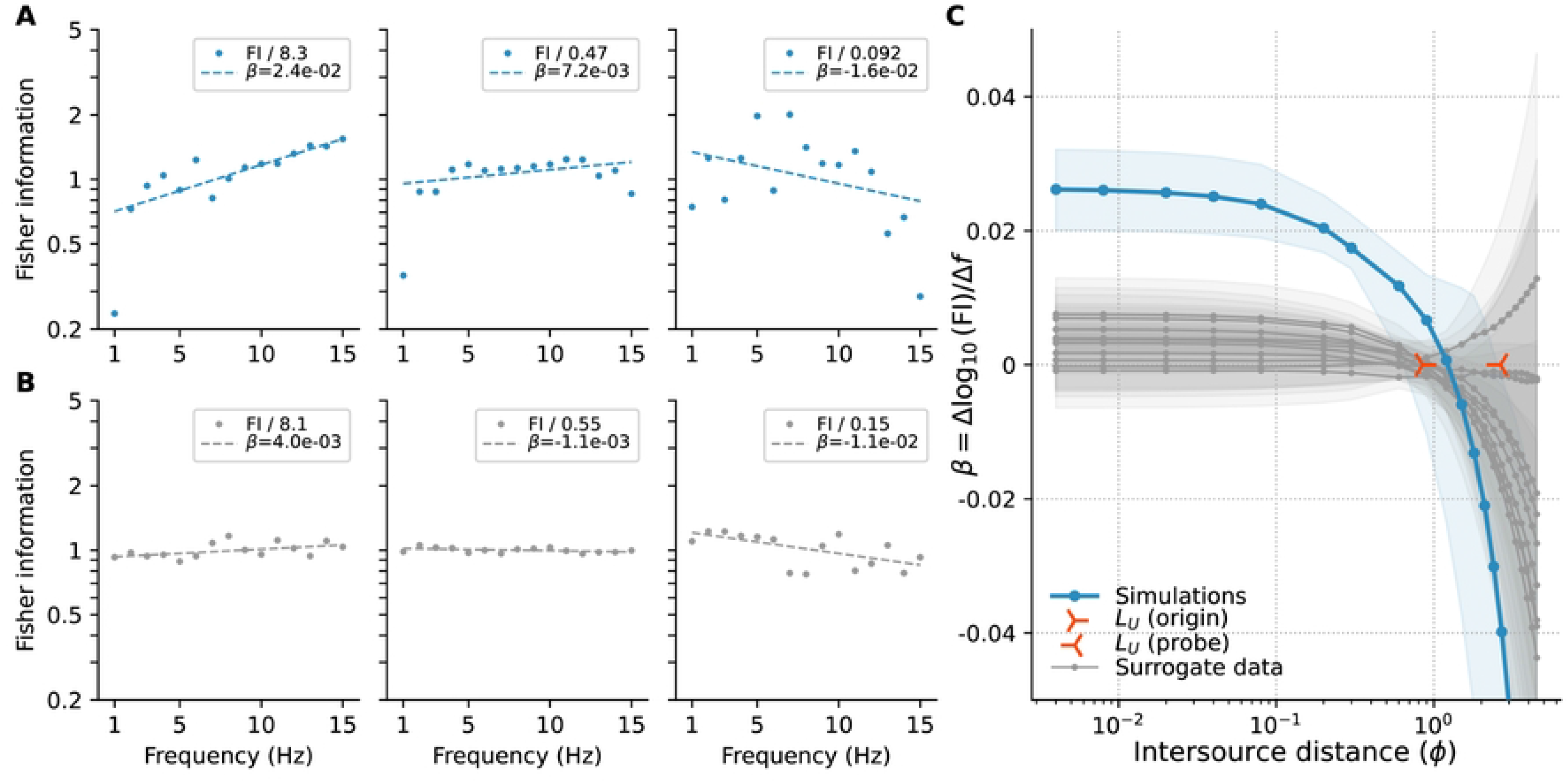
Fisher information vs. frequency. **(A)** Base 10 logarithm of the Fisher information at each frequency component (dots) and linear fits (dashed lines), for intersource separations of *∼*0.1, 1, and 2 pitches (left, middle and right panel, respectively), for the simulated plumes using a 1-second Hann window. Coefficients *β* of the linear fits are stated in the legends. Data have been scaled by the amount shown in each legend to ease comparison. **(B)** As in panel A but for surrogate data where all frequencies are equally informative. **(C)** Bootstrap median (dots) and 5th-95th percentiles (bands) of the coefficients *β* of linear fits to the logarithm of Fisher information regressed on frequency for the CFD plumes (blue) and 10 surrogate datasets where all frequencies are equally informative (gray). Positive coefficients indicate information increases with frequency, negative coefficients that it decreases. Centres of orange markers indicate the integral length scale (*L*_*U*_, see Methods Sec 5.2.8) in the vertical direction (orthogonal to the mean flow) of the vertical velocity field, computed at the midline where the odour sources were located (‘origin’, *L*_*U*_ = 0.85*ϕ*) and at the probe location (‘probe’, *L*_*U*_ = 2.6*ϕ*). Linear fits were computed using robust regression, see Methods Sec 5.2.5.

To determine the Fisher information contained in correlations about source separation, we required a probabilistic model of these correlations and how they vary with source separation. In our initial approach, we made simplifying assumptions about the Gaussianity of the plumes and their interactions and derived the predicted correlation distributions to be an asymmetric Laplacian (Eqn 22). While this distribution fit the data well (Fig 7), we found that it deviated from the observations systematically due to its inability to describe the high probabilities of low correlation events (i.e. intermittency). This is perhaps unsurprising given that instantaneous plume distributions are known to be non-Gaussian [32] with the exception of large distances from the source, exhibiting transitions from exponential-like behaviors near the source through a log-normal transition to the far-field Gaussian regime [4]. To address these deficiencies, we then modeled the correlations directly using the asymmetric Laplacian as a template. We firstly extended it to incorporate intermittency by attributing very small correlations to noise. Secondly, in addition to the asymmetrical Laplacian, we also evaluated the ability of the Gamma and generalized inverse Gaussian distributions to describe the distribution of observations flagged as true correlations. We found that the extended models were able to capture the observed correlation distributions quite well across all source separations and frequencies (Fig 7H). The improved fits also indicate that the joint distribution of coefficients is not mulitvariate Gaussian. An interesting avenue for future investigation would be to determine the joint distribtion of Fourier coefficients that would predict the same parameteric form for the distribution of correlations as our model when we fitted the correlations directly.

We aimed to determine whether high frequencies contained more spatial information than low frequencies. We found that this was the case when sources were close together. At intermediate source separations, all frequencies were equally informative. Low frequencies were more informative when odour sources were far apart. However, the absolute amount of information in these latter cases was much lower, and presumably more easily obscured by noise. Thus, when correlations contain enough spatial information to rise above the noise floor, it may be contained mainly in high frequencies. Therefore the sensitivity to high frequency correlations recently observed in mice [36] may endow animals with fine spatial resolution when locating odour sources in the environment. We note, however, that diffusion acts to smear out high frequency information more quickly than low frequency information, given the sharper concentration gradients associated with these fluctuations. The distance that high frequency information can persist in a given olfactory context is therefore set by the mean wind speed, the strength of turbulent straining, and the odourant molecular diffusivity.

We observed that correlations dropped approximately exponentially with intersource distance, see e.g. Fig 3C. This exponential decrease with intersource distance has been previously observed in the fluid dynamics literature [42]. These correlations are composed of the component correlations at each frequency component. Therefore, there is considerable latitude in the way the component correlations may decay: they are bounded by the variance at each frequency and must sum at each intersource separation to the overall correlation at that separation. Nevertheless, we observed that the component correlations all had the same, nearly exponential form (Fig 9A). Explaining the reason why the component correlations decay exponentially would be an interesting topic of future work.

Our work highlights the importance of window shape when analyzing signals where information is distributed across the frequency spectrum and where spectral power spans a wide range. Our information measures were computed for 1-second windows. Longer windows would reduce leakage effects, but would lengthen the time needed for behvioural decisions. An interesting question is how the distribution of spectral changes depends on the time windows used — or in ethological terms, which frequencies the olfactory system should attend to if odour source localizations have to be made quickly. We leave the investigation of these important questions to future work.

Fisher information is a ‘local’ measure about how a given signal can distinguish a given parameter value from neigbouring values. Our expression for Fisher information Eqn 32 is accordingly local and depends on the value of component correlations *ρ*_*n*_(*s*) at a given intersource separation and how they change around that point. To actually evaluate the Fisher information we used parametric fits to these component correlations (Eqn 33). In contrast to the local nature of Fisher information, these parameteric fits are ‘global’ in that the fitted value at each intersource separation is influenced by all the data, not just those local to the separation. For example, deviations at large intersource separations can affect the length constant *γ* of the fit, which will in turn affect the value of Fisher information reported for small intersource distances (and all others). It will therefore be useful to investigate whether fitting approaches that are more local, such as splines, produce better estimates of the Fisher information.

We chose to compare the two plumes by computing their Pearson correlations. We chose the Pearson correlation because it is insensitive to the mean and scale of the signals being compared and only registers their covariation, and it is this covariation that reflects the relative separation of the odour sources. By our choice of Pearson correlation we do not mean to imply that it is the optimal measure for decoding intersource distance. It is, rather, a simple measure whose information content provides a lower bound for the total information present in the interacting plumes about source separation. Other measures, for example those based on event timings (see e.g. [43]) may be more effective for odour source localization.

We used Fisher information as our information measure. An obvious alternative is mutual information between the intersource distances and correlations. One advantage of Fisher information is that it is based on the likelihood function *p*(*r*_*n*_|*s*) only and does not require the specification of a prior distribution on intersource distances. Mutual information requires the joint distribution *p*(*r*_*n*_, *s*) of intersource distances and correlations and thus *does* require a prior on intersource distances – or alternatively a prior *p*(*r*_*n*_) and likelihood *p*(*s*| *r*_*n*_) for correlations, which seem harder to specify. However, this is not a major shortcoming as a prior on intersource distances would not be hard to motivate or to determine empirically for a given environment. The type of information provided by Fisher and mutual information are also different. Fisher information is ‘local’ in the unknown parameter (intersource distance) and indicates how discriminable a given value of that parameter is from another value infinitesimally close. Hence it is a function of the unknown parameter, as seen in Fig 10. It also bounds the variance of unbiased estimates of intersource distance from correlations, and would thus be particularly relevant if the animal needs to accurately localize the relative locations of odour sources. Mutual information is a ‘global’ measure in that its computations involves averaging over the full joint distribution of intersource distances and correlations. We leave the task of determining which of these two measures of information is most relevant for quantifying the spatial information in plumes to future work.

### 3.2 Dispersion, coalescence, and coherent flow structures in naturalistic plumes

Our multisource plume datasets were generated in two-dimensional, spatially-decaying grid turbulence. The statistics of these plumes are broadly consistent with those observed in diverse naturalistic olfactory contexts, regardless of interesting phenomenological differences between two-dimensional and three-dimensional turbulent flows [44]. These consistencies include the observation of positive correlation regimes far from the source with clear source separation effects (Fig 3), the importance of intermittency in describing correlation distributions (Fig 7), the success of exponential-like distributions in describing correlations (Eqn 22 and Fig 7), and the exponential decay of correlations with increasing source separation (Fig 8). Given these broad consistencies, the analytical toolbox we present for quantifying the Fisher information in the spectral components of correlations about source separations is likely well-suited for applications to many odor landscapes relevant to diverse olfactory contexts.

Given the evidence for spectral information contained in correlations about source separation, what are the dynamical processes that drive these correlations? Lagrangian coherent structures (LCS) provide an intuitive framework for understanding the physical processes that drive coalescence of initially-distant odour sources and thus the statistics of their correlations. LCS arise from a dynamical systems view of fluid turbulence, where stable and unstable manifolds arise in regions of exponential fluid straining [45].

These manifolds represent repelling and attracting regions of the flow, respectively. Attracting structures are regions of disproportionate importance in odour dispersion because they closely correspond to spatial regions in which single odours evolve and initially distant odours coalesce in chaotic flow environments [46]. A representative instantaneous attracting LCS computed using the backwards finite-time Lyapunov exponent (FTLE) is shown in Fig. 13 alongside the underlying flow-field. Ridge lines in the FTLE field show strong spatial correlations with odour pair coalescence.

**Fig 13.**
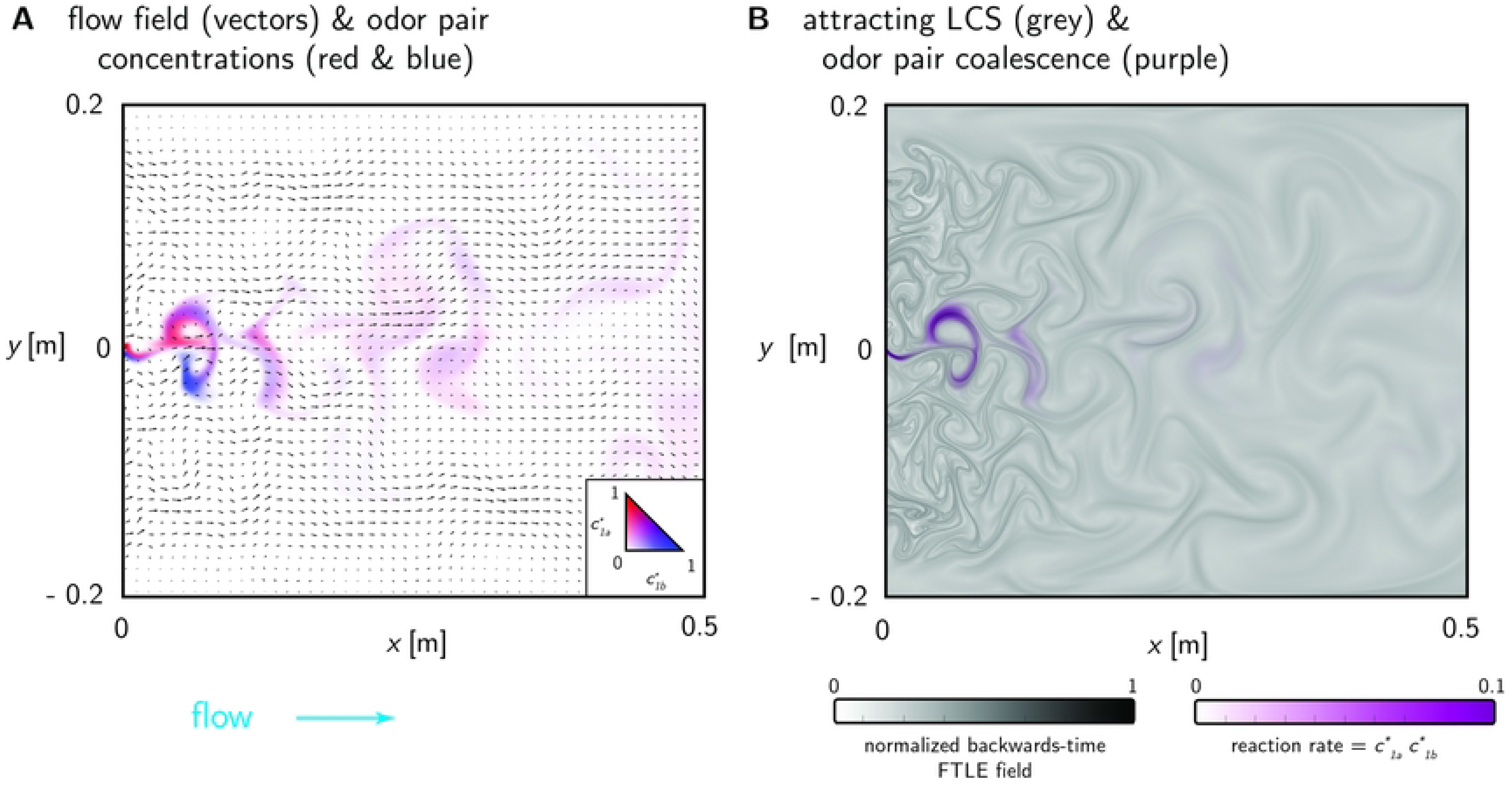
Lagrangian coherent structures. **(A)** An instantaneous snapshot of the flow field (vectors) and the concentration fields of a pair of odours (*c*_1*a*_ in red and *c*_1*b*_ in blue). **(B)** The corresponding attracting LCS quantified through the backwards-time FTLE field (grey) and”reaction rate” of the odour pair (the product of their concentrations, purple).

The exact structures of LCS and their evolution in space and time will vary among turbulent flows occurring in diverse odor landscapes. The LCS dynamics will largely determine the correlations between initially-separate odour sources, and therefore the specific flow signature inherent in the LCS field will manifest in any analysis of plume information. Thus, we suggest that LCS be used to properly contextualize our analyses and findings in order to better relate our findings to naturalistic plumes. A full quantitative linking between spectral plume information and statistical characteristics of the LCS field could be an intriguing subject for future work.

## 4 Conclusion

In conclusion, we have outlined an approach for quantifying the spectral information in odour plume correlations about odour source locations. We have tested our approach on two different simulated two-dimensional turbulent flows. Owing both to the variety of odour landscape characteristics in naturalistic plume environments and to the preeminent role of CFS in driving both dispersion and coalescence processes in natural as well as simulated plumes, we expect that the method outlined here will also be applicable to describing the spatial information contained in three-dimensional plumes.

### 5 Methods

All of our simulation data, and the code for our simulationes and analyses are provided at https://github.com/stootoon/fisher-plumes.

### 5.1 Computational fluid dynamics

We numerically modeled two-dimensional grid-turbulence in a wide wind tunnel and introduced an array of unique odour sources to generate chaotic multisource plumes (Fig 2). These data allowed us to i. investigate spectral correlations between odour sources as a function of source separation, ii. compute spectral Fisher information content encoding source separation and location, and iii. evaluate spatial structure in these entities. To test the robustness of the analyses to generalized plume environments, we conducted a second set of numerical simulations using different flow and odour parameters yielding an additional plume dataset with different spatiotemporal characteristics. Below we detail the CFD methods used for all analyses in the Main Text. We provide an alternative set of supplemental data derived from other numerical implementations in the Supporting Information Sec S3.

#### 5.1.1 Model domain, simulations, and meshing

The model domain (Fig 2) consists of a wide rectangular wind tunnel (0.62 m x 0.4225 m, *x* by *y*) with a mixing grid cylinder array distributed across the flow development inlet section (0.12 m in streamwise length). An array of 16 odour sources (8 source pairs mirrored about the domain centerline at *y*=0) of width *L*_*f*_ was distributed spatially across a transect a short distance downstream of the mixing grid at *x* = 0, covering a range of source separations *s* for analysis (Table 1). Source *y*-locations were constrained to the central 1/4 of the total inlet width to minimize walls effects, i.e. any impacts on cross-stream plume spread due to growth of the velocity boundary layer along the lateral walls. A uniform inlet velocity upstream of the mixing grid produces a chaotic flow field downstream through the nonlinear interactions of wake structures shed by the mixing grid. Simulations spanned a dimensional duration 70 s consisting of i. a 10 s start up period to establish fully chaotic flow conditions throughout the model domain and ii a subsequent 60 s analysis period containing dynamic multispecies plumes generated by the source array issuing into the chaotic flow environment.

Numerical simulations were performed via finite element discretization of the Navier-Stokes and continuity equations (Eqs. S1 and S2) governing fluid flow, and the coupled, non-reactive advection-diffusion equation (Eqn. S4) governing odour transport and diffusion (see Sec S2). The COMSOL Multiphysics package (ver. 6.0) was used to generate the mesh in the model domain described below and to solve the system of equations resulting from the weak-form discretization of the governing equations, subjected to prescribed initial and boundary conditions described below. Geometric, flow, fluid, and odourant properties for model runs in the domain depicted in Fig 2 are summarized in Table 1.

The unstructured finite-element mesh contained 289,607 total triangular and quadrilateral elements (110 elements/cm^2^ on average), locally refined in spatial regions with strong fluid velocity or odour concentration gradients, notably along the solid surfaces of the mixing grid cylinders and lateral walls and near the finite-width odour sources where strong initial odour gradients persist. The level of mesh refinement (increasing model degrees of freedom) was iterated to a final spatial resolution sufficient to resolve the smallest anticipated fluid velocity and odour concentration gradients. The final mesh implemented for all model runs had typical maximum and minimum element sizes (spatial resolution) of approximately 2 mm and 0.2 mm, respectively, with approximately 35 and 10 mesh elements per pitch *ϕ* and per source diameter *L*_*f*_, respectively.

#### 5.1.2 Initial and Boundary Conditions

Initial conditions were **u**\* = 0, *p*\* = 0 and *c*\* = 0 everywhere. The mixing grid and lateral walls were no-slip conditions, **u**\* = 0, with zero total odour flux normal to the surfaces, −**n** · (**J** + **u**\**c*\*) = 0, where the diffusive flux is **J** = −**n** · *D*∇*c*\* and **u**\**c*\* is the advective flux (**n** is the unit vector normal). The velocity inlet condition was normal, uniform flow with speed *U*_*o*_, **u** = *U*_*o*_**n** and the outlet condition on the downstream end of the domain was zero pressure *p*\* = 0. For solver stability we ramped the inlet velocity from zero to *U*_*o*_ over short interval in time. Odour sources of strength *c*\* =1 were introduced as constant concentration constraints from finite-width locations (table 1), where the source profile at the origins was a smoothed top hat. The inlet and outlet odour boundary conditions were no diffusive flux normal to the boundary −**n** · **J** = −**n** · *D*∇*c*\* = 0.

#### 5.1.3 Discretization, Solvers, and Convergence

Lagrangian shape functions were used for weak-form discretization of the fluid velocity, pressure, and odour concentration fields. The order of the integration scheme was matched to the element order for all dependent variable (first-order here). The time-dependent solver employs an implicit backward differentiation formula (BDF) method with maximum second order schemes, balancing numerical stability and damping tendencies for our smoothly varying velocity and scalar concentration gradients. BDF methods use variable-order, variable step-size backward differentiation and are known for their stability [47, 48]. The variable step size taken by the solver was informed by a prescribed absolute tolerance for the nonlinear solver and an implicit formulation of the mesh Courant-Friedrichs-Lewy (CFL) number. The solution sequence for the resulting systems of equations was fully-coupled in all dependent variables (fluid velocity and pressure, scalar concentration) for each solver iteration using an affine invariant form of the damped Newton method [49]. The nonlinear systems of equations were solved iteratively with specified convergence criteria using the direct PARDISO solver optimized for parallelized solutions of sparse systems of equations ([50, 51]). The discretization schemes and solvers detailed above on the described mesh yielded good numerical stability and solution convergence with acceptable memory and computation requirements.

#### 5.1.4 Lagrangian Coherent Structures

To investigate the dynamics of the Lagrangian flow structures responsible for odour mixing we computed backwards finite-time Lyaponov exponent (FTLE) fields per [45]. The computation followed the methods described in [46], and the resulting backwards time FTLE fields provided a proxy for the attracting LCS in the flow. The integration time *T*_*LCS*_ was set to 0.6 seconds based on preliminary investigations of the spatially-varying integral time scales. The temporal resolution of the LCS computation matched the resolution of the underlying velocity data (20 ms), and the initial spatial resolution the Lagrangian tracer grid was 250 *μ*m. These integration parameters yielded well-defined FTLE fields with strong ridge lines that were in good qualitative agreement with spatial regions odour pair coalescence, suggestive of convergence in the FTLE computation.

### 5.2 Numerical methods

#### 5.2.1 Computing plume correlations

##### Fourier decomposition and windowing of plumes

To compute the correlations between a pair of plumes, a short-time Fourier transform (STFT) was applied to each plume using the `scipy.signal.stft` [52]. The STFT was computed for a specified window length using rectangular (boxcar) windows, with an overlap of half the window length, and using no padding or boundary. The application of different window shapes was implemented through the detrending function supplied to the STFT. The detrender first applied the desired window before z-scoring the result through mean-subtraction followed by scaling by its standard deviation. This detrending was performed so that the coefficients of the decomposition of two signal into trigonometric coefficients could be easily combined as in Eqn 68 to yield the components of the Pearson correlation contributed by each harmonic. The conversion of Fourier coefficients to their trigonometric equivalents is completely standard so we have provided the derivation in the Supporting Information. The derivation shows that a signal *x*[*n*] of length *L* with discrete Fourier transform coefficients *X*[*k*] = *u*_*k*_ + *jv*_*k*_ can be expressed in terms of sines and cosines as

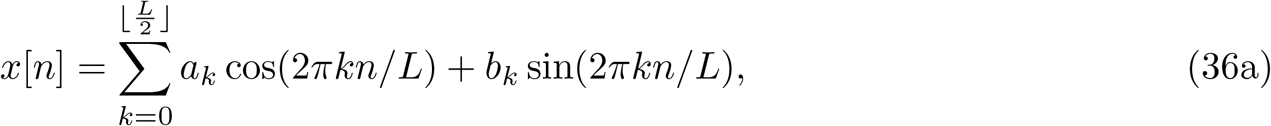

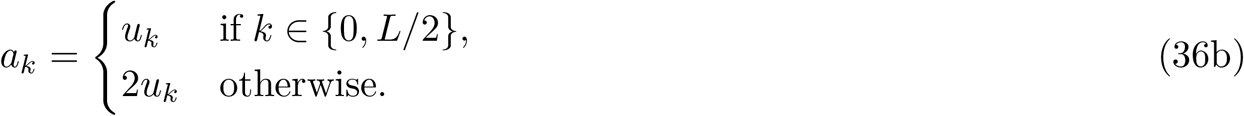

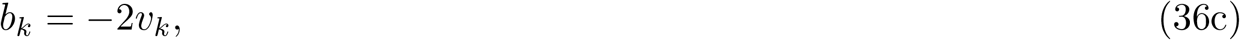

where ⌊·⌋ is the floor function.

##### Computing correlations from Fourier coefficients

The component of the correlation between two plumes for a given time window in a given harmonic was computed by combining the sine and cosine coefficients from each source according to Eqn 3. To compute the full Pearson correlation we summed the harmonic correlations over all harmonics as in Eqn 2. To compute the distribution of correlations at a given intersource distance we pooled all correlations for all time windows computed for all pairs of sources separated by the given intersource distance. Cumulative distribution functions for some of this data are plotted in the top panels of Fig 7.

#### 5.2.2 Fitting the distribution of correlations

We fit intermittent and non-intermittent versions of three different models to the correlation data. The models differed in the probability distributions they used to fit the correlations. Each model was based on a probability distribution on non-negative values. To extend the domain to cover negative values, the same base distribution was used but applied to the absolute value of the negative values, and with new set of parameters to cover these values, and the overall distribution normalized to one. That is, for a base distribution on non-negative values with parameters *θ*_+_,

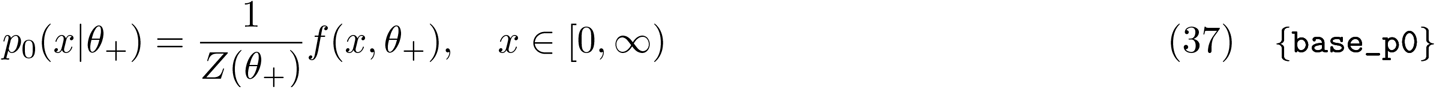

the extended distribution was defined as

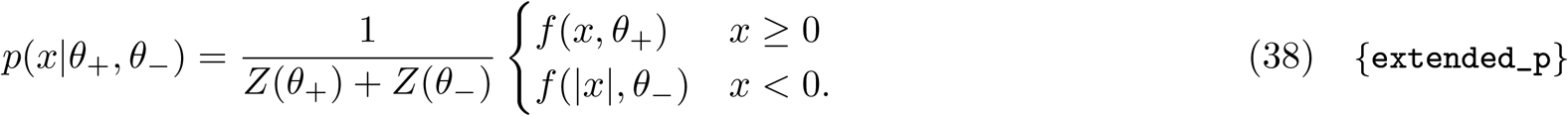

For example, the asymmetric Laplacian distribution in Eqn 24 corresponds to a base Exponential distribution, with parameters *λ* = (1 + *ρ*_*n*_(*s*))*/*2 covering the positive correlations, and *μ* = (1− *ρ*_*n*_(*s*))*/*2 covering the negative correlations.

**Note:** In what follows we will frequently refer to the extended models by their base distributions, so e.g. Exponential when referring to the asymmetric Laplacian distribution.

##### Evaluating models by comparing CDFs

We evaluated correlation models by comparing their predicted cumulative distribution functions (CDFs) to those we observed.

To compute the empirical CDF for a set of *N* observed correlations, we first sorted the values in ascending order, to yield

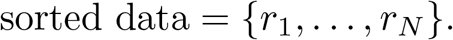

We then computed the empirical CDF evaluated at each data point *r*_*i*_ as

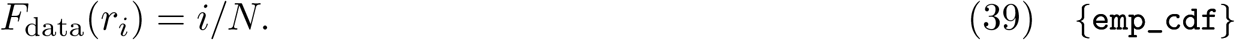

The predicted CDF can be derived using the probability density function of Eqn 25, which is

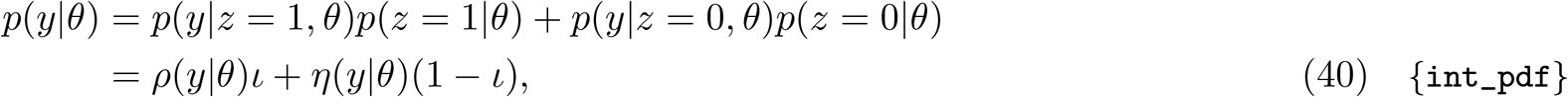

where *ρ* and *η* are the correlation and noise distributions, respectively (see Eqn 29 for a particular case). The CDF is then a linear combination of the CDFs for the correlation and noise distributions:

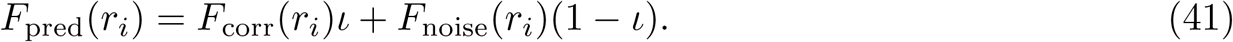

When comparing CDFs the *R*^2^ value can be overly optimistic since both CDFs start at zero and increase monotonically to 1. Therefore, we compared CDFs more stringently by computing the largest absolute difference in their values, which is bounded below by 0 and above by 1. That is, we computed

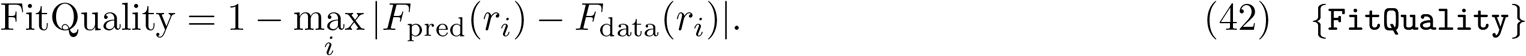

When numerically computing the predicted CDFs we sometimes encountered values outside of the valid [0,1] range. This was particularly the case when using scipy.stats.geninvgauss to compute the CDF of the generalized inverse Gaussian. Therefore, when computing fit quality we only used the subset of points for which the CDF values were valid.

##### Fitting models using expectation-maximization

To fit models with intermittency we had to determine which of a set of observed correlations {*y*_1_, …, *y*_*N*_} at a given harmonic and intersource distance were attributed to noise. From Eqn 25 this requires determining the values of the binary latent variables {*z*_1_, …, *z*_*N*_} corresponding to each of the observations. To fit the parameters *θ* of a model while accounting for these latent variables, we used the Expectation-Maximization algorithm [53]. Initialized with an initial guess of the model parameters, this algorithm consists of repeatedly estimating the values of the latent variables (the ‘E’-step), then using those estimates to update the estimates of the model parameters (the ‘M’-step). Below, we summarize the parameter updates. Derivations are supplied in the Supporting Information Sec S4.

The E-step update for all models at the *t* + 1’st iteration has the simple form

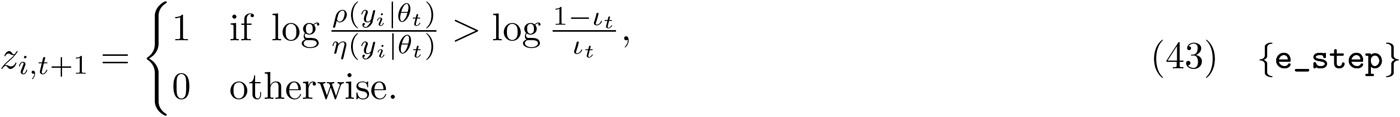

Here *ρ*(*y*_*i*_|*θ*_*t*_) and *η*(*y*_*i*_, *θ*_*t*_) are the probabilities of the observation *y*_*i*_ under the correlation, and noise, distributions, respectively, *θ*_*t*_ is the current estimate of the model parameters, and *ι*_*t*_ is the current estimate of the intermittency parameter. The E-step therefore declares an observation *y*_*i*_ to have been a correlation rather than noise if its probability according to the correlation distribution is greater than its probability according to the noise distribution by a threshold that depends on the current estimated intermittency level. For the non-intermittent models, all observations were marked as being correlations.

During the M-step model parameters were updated using the current estimate of the latent variables {*z*_1,*t*_, *z*_2,*t*_, …, *z*_*N,t*_} . All models had two parameters that related to intermittency: the intermittency level *ι*, and the standard deviation *σ* of the noise distribution. These parameters were updated the same way for all models. For the non-intermittent models, *ι* was fixed at 1, and *σ* at 0. Otherwise, these parameters were updated as described below.

To update the intermittency level *ι*, it is helpful to define the data intermittency at time *t* as

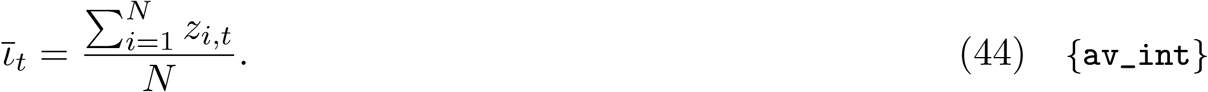

The intermittency level is updated by balancing the data intermittency against a prior on intermittency, 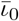,

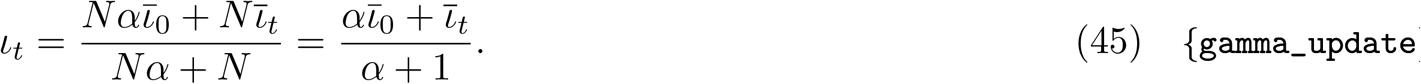

This can be interpreted as weighting *Nα* observations with an intermittency of 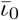against *N* observations at the data intermittency 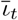. The hyperparameter 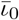was fitted using grid search (see below). The hyperparameter *α*, which sets the relative strength of the prior, was fixed at 1.

We updated the noise level by setting its variance to be the mean sum of squares of the observations attributed to noise in the E-step. Intuitively, this is because under our assumption that the noise distribution has mean zero, the mean sum of squares of the noise observations is an estimate of its variance. If the E-step did not flag any observations as noise we set the estimated noise variance to a small fraction of the variance of the observed correlations. We did this because setting the noise level to exactly zero would mean that the estimated noise variance would remain at zero for all future iterations. This is because the probability of any non-zero observations would be 0 under this degenerate noise distribution, so all observations in the next E-step would also be flagged as correlations and not noise, leaving the noise variance stuck at zero.

Letting 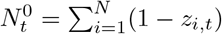 be the number of observations attributed to noise after the current E-step,

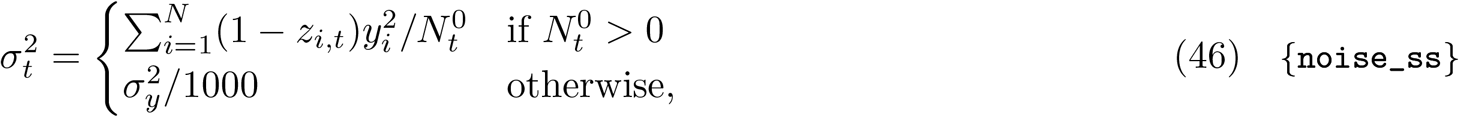

where 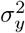 is the variance of the observed correlations.

The rest of the parameters varied by model so their updates will be described for each model separately.

###### Exponential model

In addition to the intermittency parameters, the Exponential model had two parameters, *λ* and *μ*, describing the decay rate of the positive, and negative, correlations, respectively. The M-step updates for these parameters had the closed form

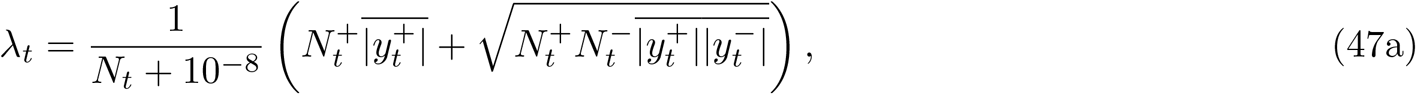

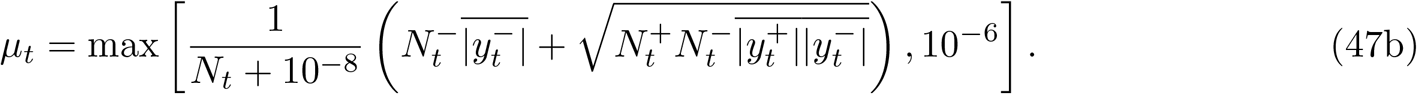

Here 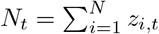 is the number of observations that were designated as correlations in the E-step. These can be split into 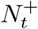 positive and 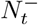 negative correlations, and 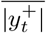 and 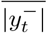 are the averages of the absolute values of the corresponding observations. The maximum operations prevents *μ*_*t*_ from being set to zero when the E-step did not designate any observations as noise. The added factor of 10^−8^ in the denominators prevents division by zero when *N*_*t*_ = 0 i.e. all observations were estimated to be noise.

###### Gamma model

The Gamma model has all the parameters of the Exponential model, plus two shape parameters, *k* and *m*, to describe the positive and negative correlations. To update the parameters during the M-step we minimized the negative log likelihood of the observations that were marked as correlations in the E-step:

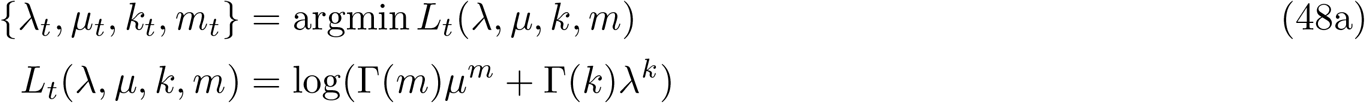

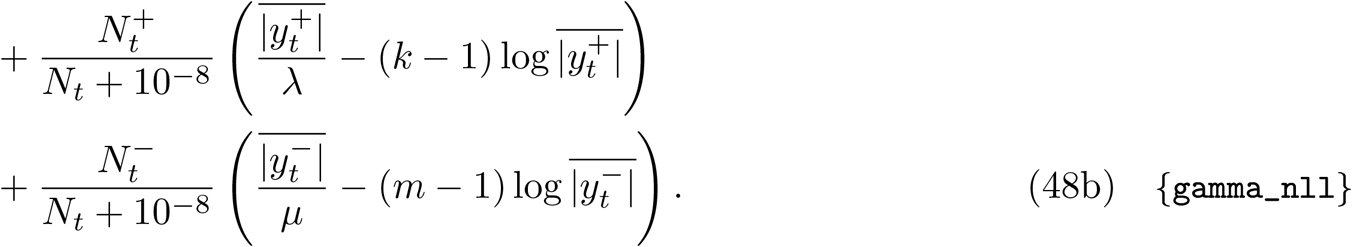

The minimization was performed using Nelder-Mead method as implemented by the scipy.minimize[52] function. The bounds of the search were [10^−6^, 10] for *λ* and *μ*, and [0, 10] for *k* and *m*. The search was initialized at the parameter values from the previous iteration, clipped to lie within these bounds.

###### Generalized inverse Gaussian

The Generalized inverse Gaussian model has all the parameters of the Gamma model, plus two additional shape parameters, *α* and *β*, for the positive and negative correlations, respectively. During the M-step we performed a damped update of the parameters towards the minimum of the negative log likelihood of the observations that marked as correlations in the E-step:

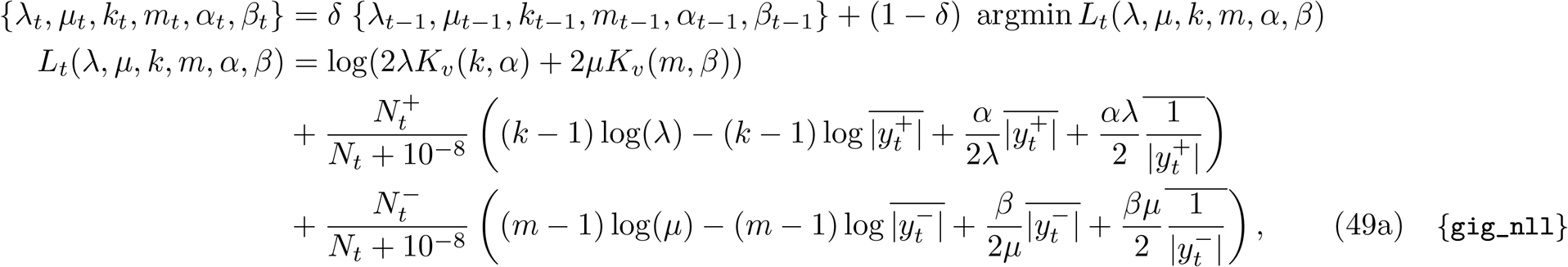

where *K*_*v*_(*a, b*) is the modified Bessel function of the second kind of real order *a* evaluated at *b*. Damping was used to aid convergence, with the damping factor *δ* fixed at 0.5. The minimization was performed the same way as for the Gamma model, with the new parameters *α* and *β* bounded to [10^−6^, 10]. The search was initialized at the parameter values from the previous iteration, clipped to lie within these bounds.

##### Ancestral initialization

The Gamma distribution subsumes the Exponential distribution, because the former reduces to the latter when the shape parameter is set to 1. In turn, the generalized inverse Gaussian distribution subsumes the Gamma distribution, because the former reduces to the latter when its *β* parameter (the coefficient of 1*/x* in the argument of the exponential) is set to zero. Because models with fewer parameters can be easier to fit, to aid the fitting process we fit our models ancestrally. That is, to fit a model using the Gamma distribution we first fit the data using an Exponential distribution, and then initialized the Gamma fit with the parameters of the Exponential, initializing the shape parameter of the Gamma to zero. Similarly, when fitting a generalized inverse Gaussian, we initialized its *β* parameter at zero, and initialized the remaining parameters from those of the best fit Gamma. The Gamma in turn was initialized with the parameters of the best fit Exponential.

##### Evaluating models using nested cross-validation

We fit a range of models to the correlation data by testing all combinations of the following hyperparameters:

- Whether the model was intermittent or not;
- If intermittent, the mean of the prior Beta distribution on the intermittency parameter, for the values {0.1, 0.2, …, 0.9}.
- The base probability for the correlations, whether Exponential, Gamma, or Generalized Inverse Gamma.

The remaining hyperparameters, listed below, were held constant at the stated values

- The minimum (0) and maximum (10) shape parameters for the Gamma and Generalized Inverse Gaussian distributions.
- The strength (1) of the Beta prior on intermittency.

We characterized these models in two ways. First, we estimated the performance of each model in fitting the correlation data. To do so, we repeatedly split the data into training (67%) and test (33%) sets. Each model was fitted to the training set and evaluated on the test set. Its performance was estimated as its average performance over three such random splits.

Secondly, we estimated the rank of each model, so that we could say e.g. which model was best overall, or which was the best among those with intermittency. One way to do this would be simply by ranking the performances we computed above. However, doing so would risk overfitting. To see why, note that by ranking the models we are in effect fitting the hyperparameters. Ranking the models using the same data used to evaluate their performance would then mean estimating performance with the same data used to fit the (hyper)parameters, risking overfitting.

To estimate the rank of each model while avoiding making this estimation on the same data that was used to estimate performance, we performed an inner loop of cross validation for each iteration of the outer loop.

In each iteration of the inner loop, the training set provided by the outer loop was further split at random into a training subset (67 %) and a validation set (33%). Models for every setting of the hyperparameters were fit on the training subset, and evaluated on the validation set. Models were ranked by their mean validation performance over three such random splits. The resulting model ranks were then averaged over iterations of the outer loop.

To see why this nested cross-validation procedure avoids overfitting, observe that each iteration of the outer loop yields, first, an estimate of model performance as computed on the test data, and second, model rank as determined from the training data by the inner cross-validation loop. Therefore, in each outer loop iteration, model rankings and model performance are estimated using different data. We then reduce the noise in these estimates by averaging over iterations of the outer loop.

#### 5.2.3 Modeling the dependence of Fourier coefficients on intersource distance

To compute the empirical distribution of the Fourier coefficients from two sources at a fixed intersource distance *s* and for a given harmonic *n*, we first listed all pairs of sources that were the desired distance apart. Our source locations only differed in the *y*-coordinate, and we used a signed distance, so dist(*y*_*i*_, *y*_*j*_) ≜ *y*_*i*_ − *y*_*j*_. Therefore, a desired intersource distance *d* yielded a list of ordered pairs of sources (*y*_*i*_, *y*_*j*_) such that *y*_*i*_− *y*_*j*_ = *d*. We then concatenated the sine and cosine coefficients in the given harmonic for all time windows across all sources in the first coordinate of each pairing. This yielded a vector of length

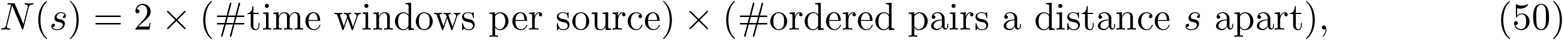

where the factor of 2 is because of the concatenation of both sine and cosine coefficients. The length is a function of intersource distance *s* because the number of pairs a given distance apart was distance-dependent. We then stacked this vector on top of the same concatenation of data applied to the sources in the second coordinate of each pairing. This yielded a 2 × *N* (*s*) matrix **Data**_*n*_(*s*),

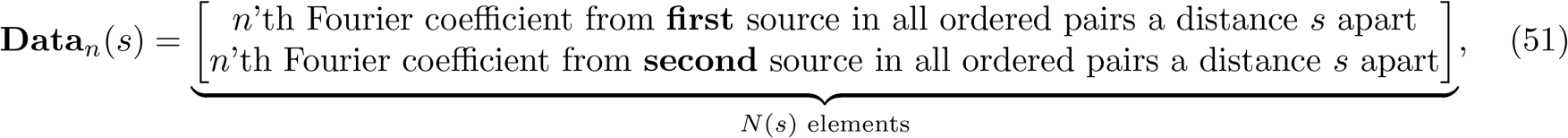

where each element of the first row was one Fourier coefficient computed in one time window from one source, and the corresponding element in the second row was the corresponding coefficient for another source a distance *s* apart. We plotted some of this data in Fig 6.

The columns of **Data**_*n*_(*s*) represent samples from the joint distribution *p*(*a*_*n*_, *c*_*n*_|*s*) of Fourier coefficients for the *n*’th harmonic, from sources a distance *s* apart. To fit a zero-mean bivariate Gaussian to **Data**_*n*_(*s*) as in Eqn 17b we computed the principle variances *λ*_*n*_(*s*) and *μ*_*n*_(*s*) along the major (*y* = *x*) and minor (*y* = −*x*) axes, respectively. To determine these we computed the variances of the projections of the data on the unit vectors 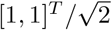 and 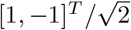, respectively,

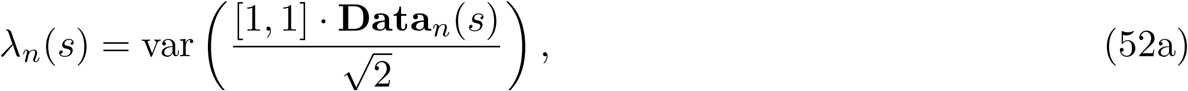

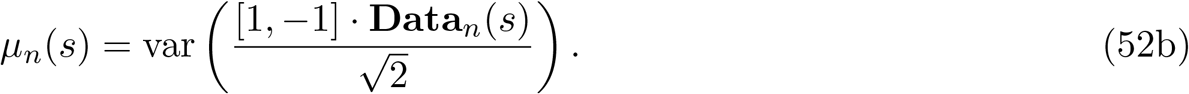

We computed the variance 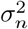 as the mean of the principle variances at *s* = 0,

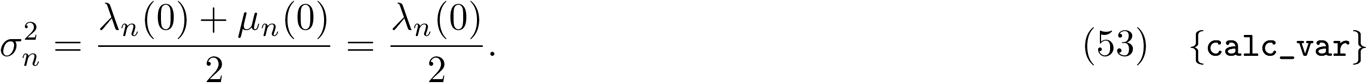

This is equivalent to computing the pooled variance of all coefficients from all sources for the given harmonic.

#### 5.2.4 Parametric fits to *ρ*_*n*_(*s*) as a function of distance

We next fit each *ρ*_*n*_(*s*) to the parametric form in Eqn 33 by non-linear least squares using `curve_fit` from the `scipy.optimize` package[52]. The parameters to be learned were the length scale *ι* and the offset *b*. We constrained the parameters to be non-negative, and initialized *ι* to 1 and *b* to 0. All other parameters to `curve_fit` were kept at their default values. Some example fits and parameters learned are in Fig 8.

#### 5.2.3 Regressing information on frequency

To produce the results in Fig 12 we regressed, for each value of intersource distance, Fisher information on frequency. The Fisher information at some intersource distances exhibited outliers that overly influenced the results of a standard linear regression, as judged by visual inspection. In addition to filtering out data points for which the computed Fisher information had NaN, infinite, or negative values, we took two measures to make the results more robust to these outliers. First, we limited the range of frequencies considered to be 1 Hz to ∼ 15 Hz. This was because some of the data, particularly in the supporting simulations, showed sudden drops in Fisher information above ~15 Hz. Secondly, we replaced ordinary linear regression with robust regression. We used Huber regression, as implemented in scikit-learn’s `linear_model.HuberRegressor`. We used the default parameters, except for `max_iter` which we set to 10,000.

#### 5.2.6 Bootstrapping procedure

To estimate the variability of the various statistics used in our approached we computed their bootstrap distributions. These statistics were computed from the trigonometric form of the Fourier decompositions computed for each plume in each time window. Therefore we first computed these coefficients for all time windows, and then created each bootstrap dataset by sampling time windows with replacement until we had as many time windows as the original dataset. We generated 50 bootstrap datasets in this way. We then computed our various statistics using the data for the time windows chosen for each bootstrap dataset. This then gave us as many point estimates of each statistics as bootstrap datasets, from which we computed e.g. the 5th, 50th and 95th percentiles of Fisher information shown in Fig 10A.

#### 5.2.7 Generating surrogate data

In brief, we generated surrogate data consisting of artificial plumes with covariance structure similar to real plumes but for which we could adjust the amount of spatial information. To generate these plumes we randomly generated coefficients according to the bivariate normal model of Eqn 17b using covariance kernel *K*(*n, s*) to relate the coefficients from plumes a distance *s* apart. Specifically,

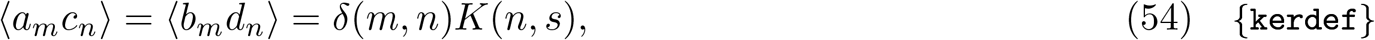

where as usual, *a*_*m*_ and *c*_*n*_ are the cosine coefficients for the *m* and *n*’th frequency components, generated by two plumes a distance *s* apart, *b*_*m*_ and *d*_*n*_ are the corresponding sine coefficients, and expectations are taken over time windows, and the *δ* function *δ*(*m, n*) ensures that coefficients for different harmonics are uncorrelated. We combined these coefficients with their corresponding sine and cosine waveforms to generate the surrogate signals.

In detail, we generated signals for *M* sources, each of length 2 ⌊*L/*2⌋ + 1, by first creating the kernel relating the trigonometric coefficients of their Fourier decompositions. The signals had zero-mean, therefore for each source, we needed to specify ⌊*L/*2⌋ sine and cosine coefficients, corresponding to the normalized frequencies 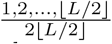, covering all positive harmonics of the fundamental up to the Shannon limit. The kernel needed to specify how each harmonic at every source was correlated with each harmonic at every other source, a total of *M* ^2^⌊*L/*2⌋^2^ values. We organized these values into a symmetric kernel matrix of size *M* ⌊*L/*2⌋ × *M* ⌊*L/*2⌋. The elements of this matrix were determined as

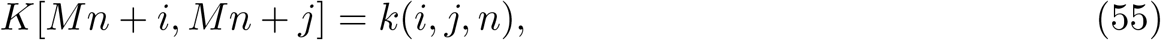

where *k*(*i, j, n*) was the desired covariance of the trigonometric coefficients for the *n*’th harmonic for sources *i* and *j* (see below). We assumed coefficients at different harmonics were uncorrelated.

To compute cosine coefficients with covariance *K*, we first computed the Cholesky decomposition *K* = *LL*^*T*^. We then generated *M* ⌊*L/*2⌋ i.i.d. samples from a standard normal, *u*, and determined the cosine coefficients as *c* = *Lu*. We generated a second set of samples, *v*, from the standard normal in the same way, and used those to produce the sine coefficients *s* = *Lv*. Indexing these coefficients by source *m* and harmonic *n*, we produced the signals at each source by combining the coefficients with their corresponding trigonometric waveforms,

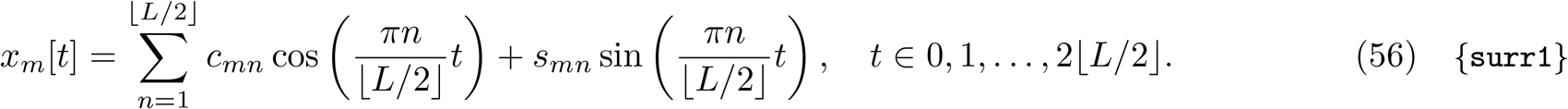

##### Kernel functions

We used several different kernel functions *k*(*i, j, n*) to generate the surrogate data in the text. Each of these kernel functions was the product of a term *S*(*n*) determining the power at the *n*’th harmonic, and a function *G*(*i, j, n*) determining the correlation between sources *i* and *j*, that in some cases depended on the harmonic. In every case, this correlation function was a function only of the absolute difference *s* = |*i* − *j*| of the sources, so it simplified to *G*(*s, n*). Thus our kernels were of the form

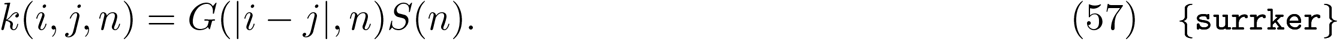

Our kernels differed by whether they used a flat power spectrum or one similar to that of the simulations. We captured both cases by using a 1*/f* power spectrum:

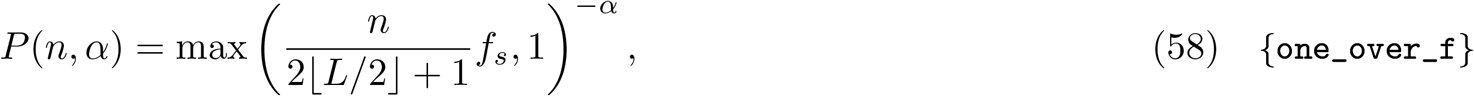

where *f*_*s*_ was the sample rate in Hz. The maximum operation provides a frequency cutoff at 1 Hz below which the power is set to 1. To achieve a ‘white’ spectrum, we set *α* = 0. To achieve a ‘pink’ spectrum similar to our CFD simulations, we set *α* = 4.

Our surrogate data also differed in the correlation functions used. For the data where all frequencies were equally informative, we set

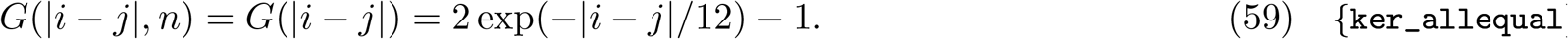

For the data where the high frequencies were more informative for sources close together than the lower frequencies, we set

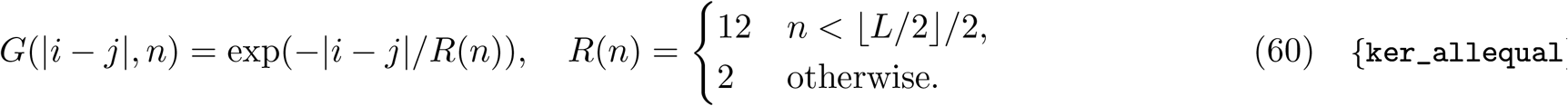

Thus the length scale of decay was 6 times shorter for high harmonics (those in the upper half of the range) than for low harmonics.

The table below summarizes the surrogate datasets and the power and correlation functions used in each.

#### 5.2.8 Computing integral length scales

We computed integral length scales from velocity autocorrelation functions as defined in [54]. For example, to compute the integral length scale in the *y*-direction at a location (*x, y*) of interest, we first computed the *y*-velocity autocorrelation function in the *y* direction at that location by computing the time average of the product of the *y*-velocity at the location of interest, *u*_*y*_(*t*; *x, y*) and that at a fixed positive displacement *r* in the *y*-direction, *u*_*y*_(*t*; *x, y* + *r*). That is, we computed

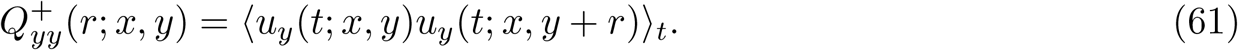

We also computed the symmetric average in the negative *y*-direction,

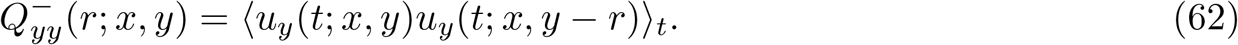

We then averaged these two to arrive at the autocorrelation for a single, positive value of *r*

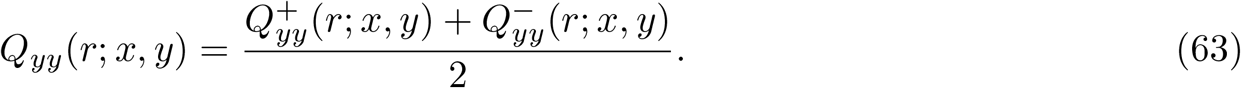

If e.g. (*x, y* + *r*) was not in the domain, then 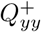 would not be computed and *Q*_*yy*_ took the value of 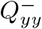, and similarly if (*x, y* − *r*) was not in the domain.

We then computed the longitudinal velocity correlation function by normalizing the velocity correlation function by its value at *r* = 0

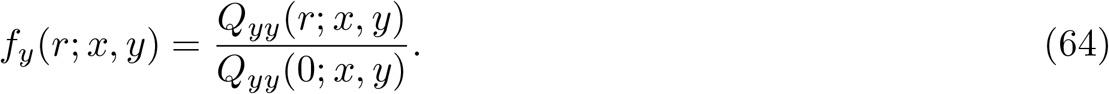

Finally, we computed the integral length scale by integrating this function from 0 to ∞:

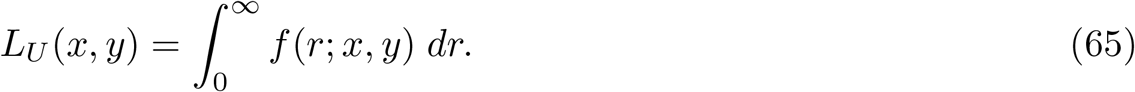

The *r* values we used were discrete and spaced Δ*r* = 0.02*ϕ* apart, so we approximated the integral above as

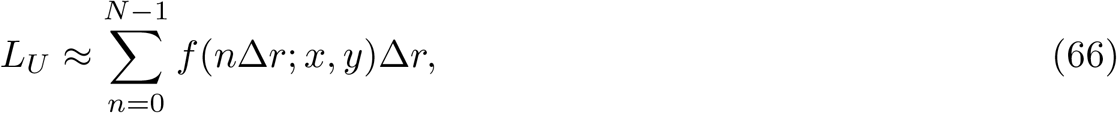

where *N* was the number of *r* values used.

In Fig S11 we have plotted the longitudinal velocity autocorrelation functions in the *x* and *y* directions, and the corresponding integral length scales, for two locations of interest.

### 5.3 Analytical methods

#### 5.3.1 Decomposing Pearson correlation

We can express two concentration profiles *x*(*t*) and *y*(*t*) observed over a window of width *T* in terms of their Fourier decompositions,

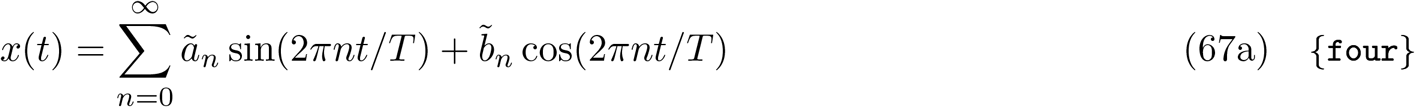

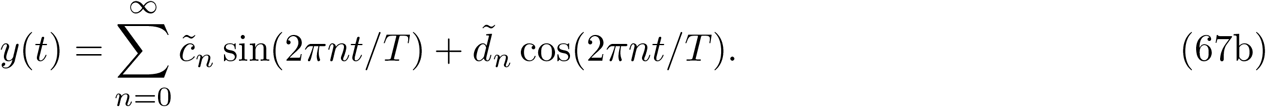

Using the orthogonality of different harmonics of the fundamental frequency we can express the Pearson correlation in terms of the Fourier components as

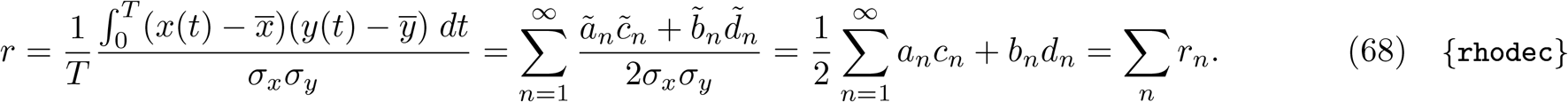

#### 5.3.2 Relating the coefficients at one source to those at another

To determine the uncertainty about the coefficients *c*_*n*_, *d*_*n*_ of a component waveform at a second source,

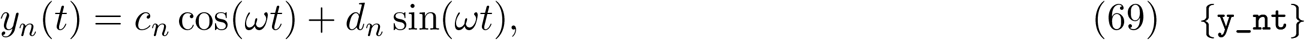

given the coefficients *a*_*n*_, *b*_*n*_ of a first,

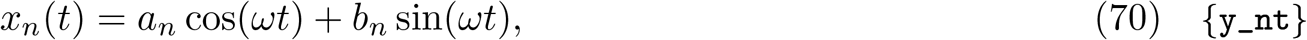

we express the former as the sum of a deterministic component formed of a scaled and phase-shifted version of the component waveform from the first source, plus noisy residual. That is,

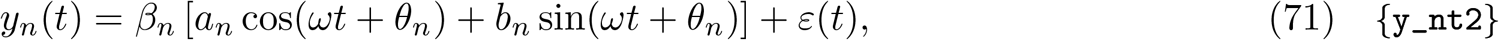

where *β*_*n*_ and *θ*_*n*_ are the best-fit scaling and phase-shift of the first source. These can be found to satisfy (see below)

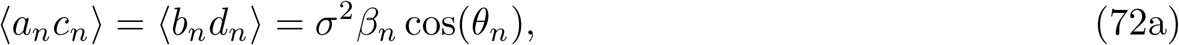

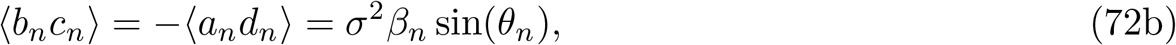

where as usual expectations are over time windows.

The residual *ε*(*t*) therefore captures the uncertainty remaining about the second component waveform, given knowledge of the first. To determine its statistical properties we express it by subtracting Eqn 71 from Eqn 70 as

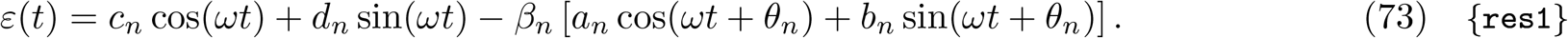

This is a linear combination of sinusoidal waveforms at the same frequency *ω*, so we can write it as

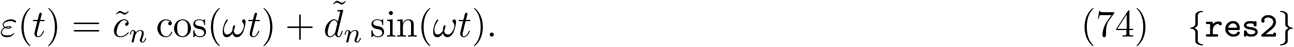

By performing trigonometric expansions, the coefficients are found to be

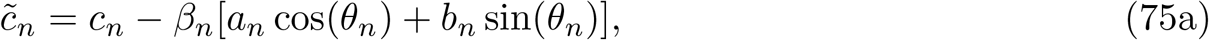

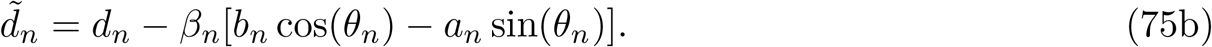

For *n >* 0, *a*_*n*_, *b*_*n*_, *c*_*n*_, and *d*_*n*_ have mean zero (Eqn 10). Therefore the coefficients above, and in turn *ε*(*t*), as linear combinations of zero-mean random variables, are also zero mean.

To determine the variance of the residuals, we first show in Supporting Information Sec S5.2 that the coefficients 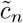 and 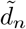 are uncorrelated. Therefore, from Eqn 74, the residual variance is the scaled sum of the variances of the coefficients. To compute these variances, we have

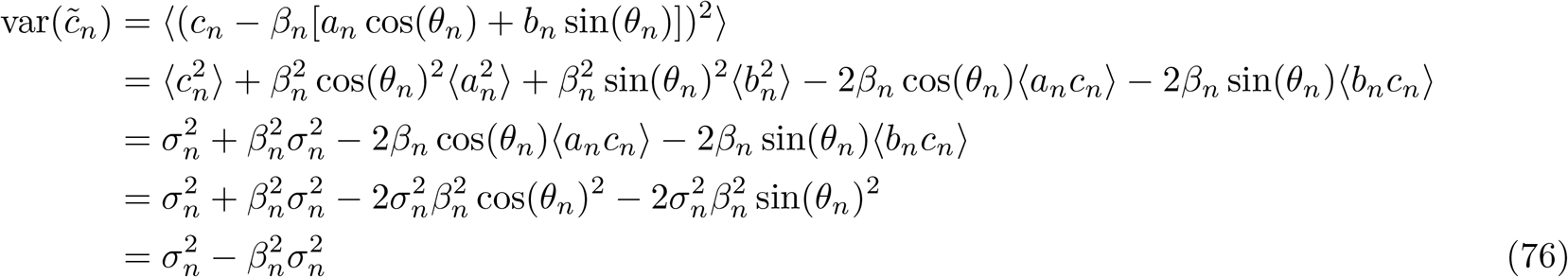

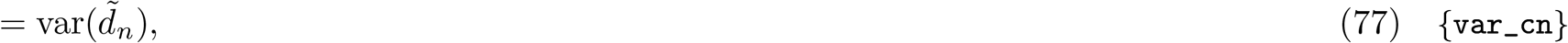

where we’ve used the relations in Eqn 75 to express coefficient correlations in terms of *β*_*n*_ and *θ*_*n*_, and the final equality follows by temporal stationarity. We then arrive at

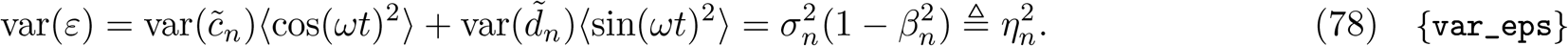

Knowledge of the mean and variance would completely specify the probability distribution of the residual if it were a Gaussian random variable. Although it can be expressed as a linear combination of the Gaussian random variables *a*_*n*_, *b*_*n*_, *c*_*n*_ and *d*_*n*_ (Eqn 73), these latter variables are not necessarily independent. Therefore, their combination is not necessarily Gaussian. To enforce this requirement we made the assumption 2.2.

To determine the scaling *β*_*n*_ and phase-shift *θ*_*n*_, we equate Eqn 71 an Eqn 71 and match coefficients of cos(*ωt*) and sin(*ωt*), to get

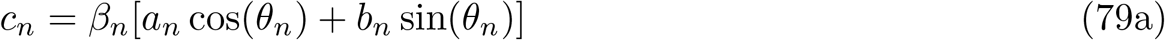

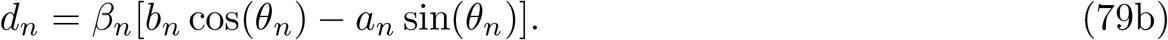

Correlating these equations against *a*_*n*_ and *b*_*n*_, respectively, and using Eqn S29 and the definition Eqn 12 of the coefficient variances 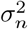, we get

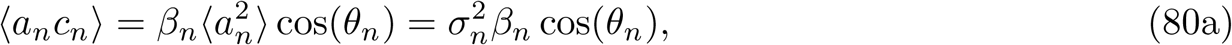

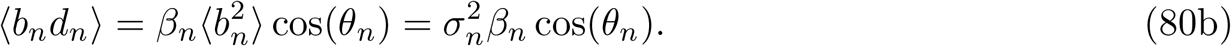

Correlating against *b*_*n*_ and *a*_*n*_, respectively, and using Eqn S30, we get

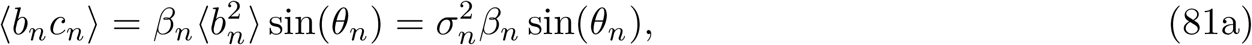

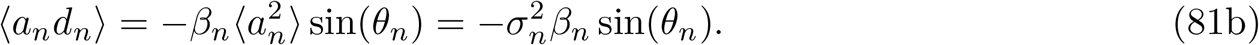

#### 5.3.3 Deriving the distribution of correlations

Here we derive the distribution *p*(*r*_*n*_|*s*) of the component correlations *r*_*n*_ for the *n*’th harmonic for sources separated by a distance *s*, given our assumptions about the distribution of Fourier coefficients, *a*_*n*_ and *b*_*n*_, from the first source and *c*_*n*_ and *d*_*n*_ from the second source. Dropping the *n* subscripts for clarity,

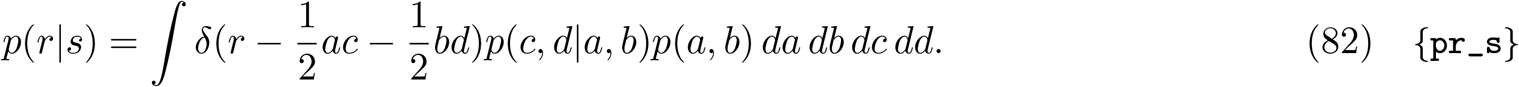

From Eqn 15 We have that

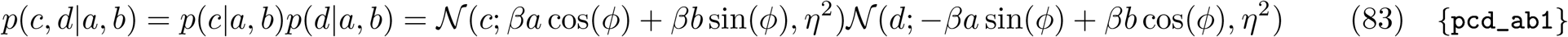

By defining **z** = (*a, b*), **x** = (*c, d*) we can write Eqn 83 as

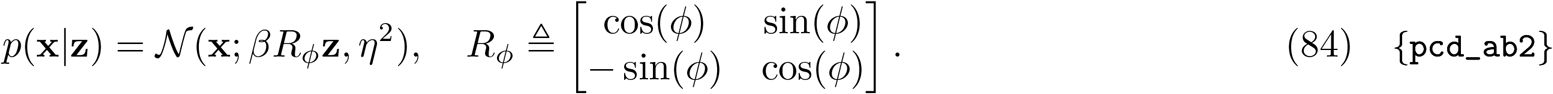

Combining this with Eqn 12 to get *p*(**z**) we have

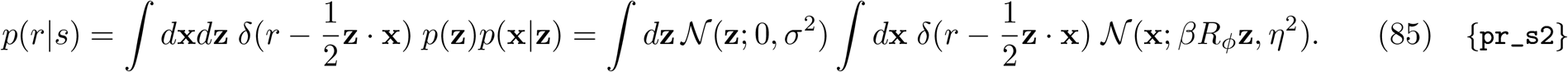

where we’ve used Eqn 12 to specify *p*(**z**). The integrand in Eqn 85 is rotationally invariant in **z** since any rotation in **z** is matched by **x** through the *δ* function. So we can compute Eqn 85 by computing it for one radial slice of **z**, say **z** = *z***e**_1_ = (*z*, 0), and multiplying the result by 2*π*. Switching **z** to polar coordinates **z** = (*z, θ*), for which

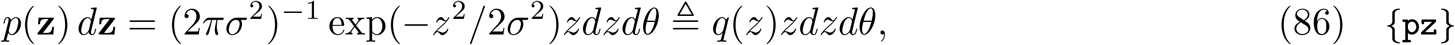

we have

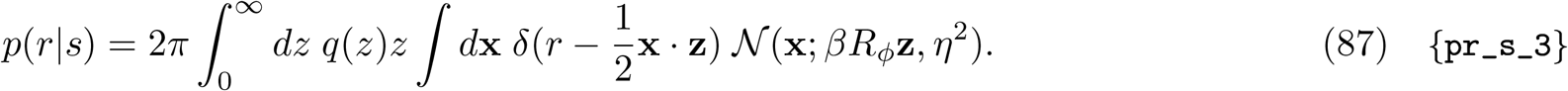

To compute the latter integral, we switch **x** from (*c, d*) to (*u, t*) coordinates, defined for **z** = (*z*, 0) as

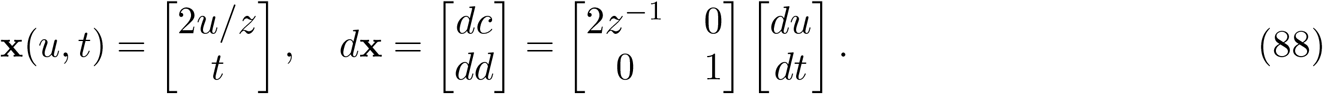

Then the area element *dc dd* becomes 2*z*^−1^*dudt* and we have for the latter integral in Eqn 87,

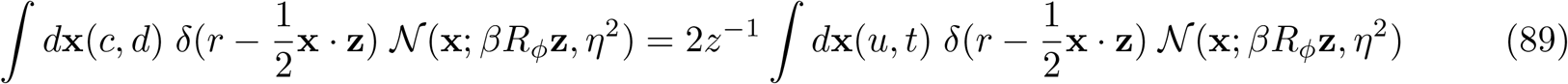

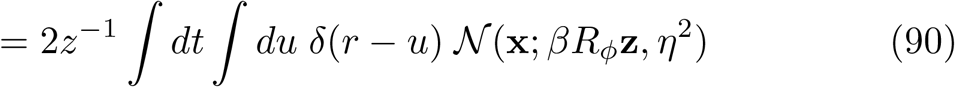

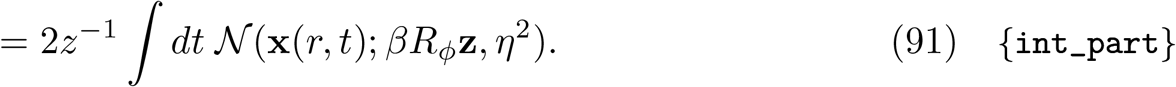

Now for our slice 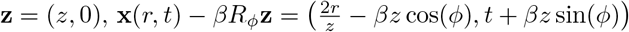, so

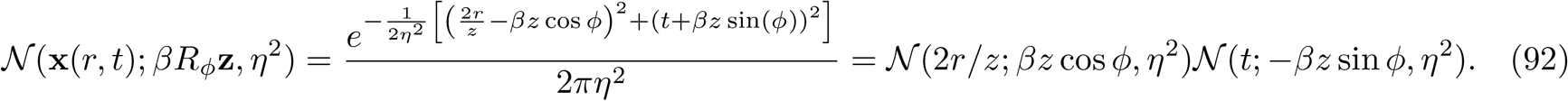

Hence Eqn 91 evaluates to

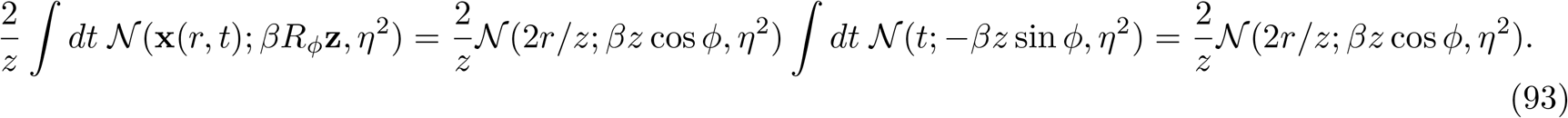

Substituting this in for the latter integral in Eqn 87, we have

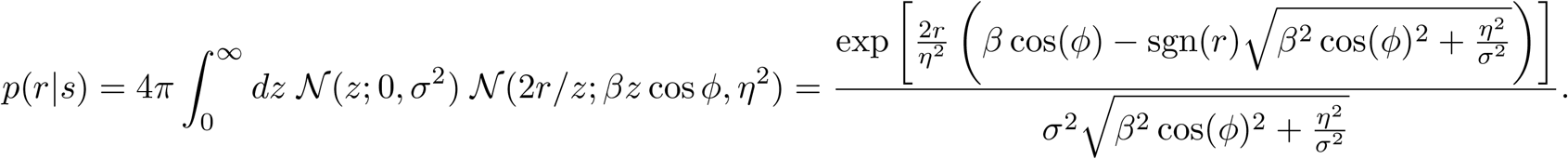

Finally, to arrive at Eqn 22, we simplify the length constant of the exponential by relating *η*^2^ in Eqn 78 to *Z* in Eqn 23 as *η*^2^ = −*σ*^−2^*Z*^2^ − *σ*^2^*β*^2^ cos(*ϕ*)^2^, and use the definition of *ρ* in Eqn 18 to get

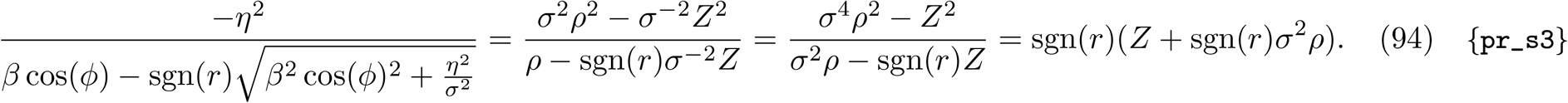

#### 5.3.4 Fisher information

To compute Fisher information for the distribution in Eqn 22, we need (dropping subscripts for clarity),

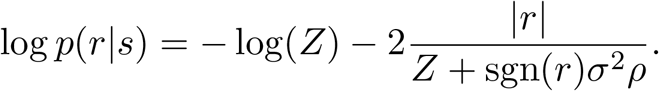

Then, defining *λ* = *σ*^2^*ρ*

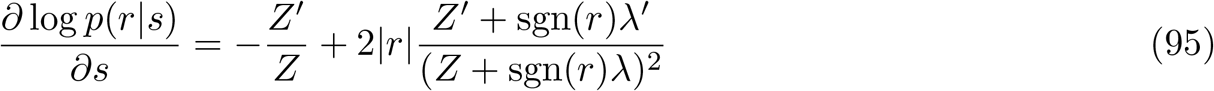

Now defining *λ*_⊥_ = *σ*^2^*ρ*_⊥_, we have 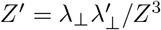 so

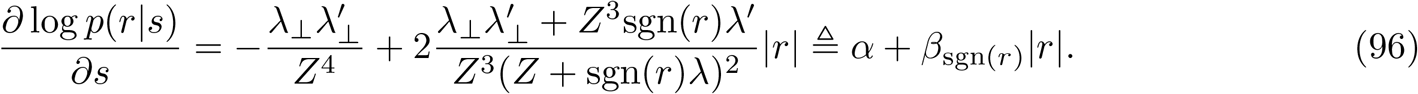

Then the square is

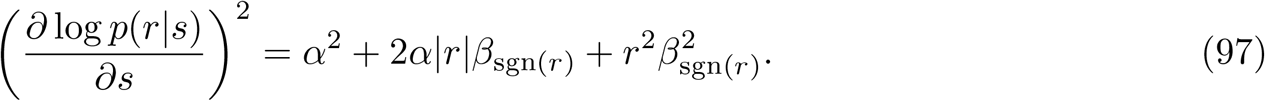

We need to compute the expectation of this with respect to *r* under *p*(*r*|*s*). We can split this into the contributions from each component.

##### Constant term

The expectation of the constant term is just

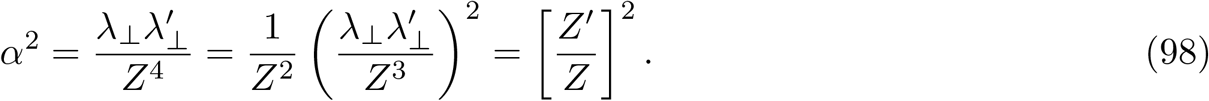

##### Linear term

Defining 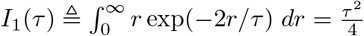 we have

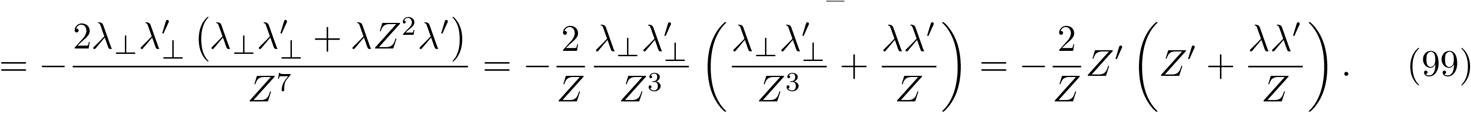

##### Quadratic term

Defining 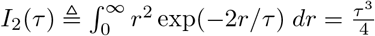we have

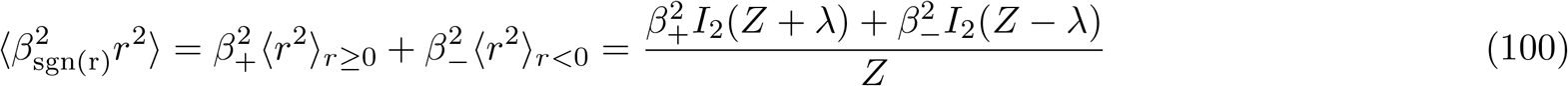

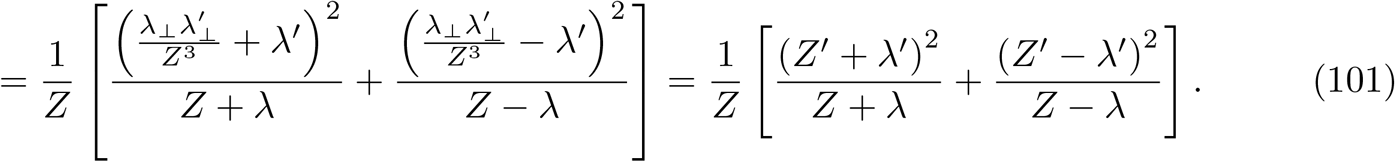

##### Fisher Information

Collecting terms, we arrive at

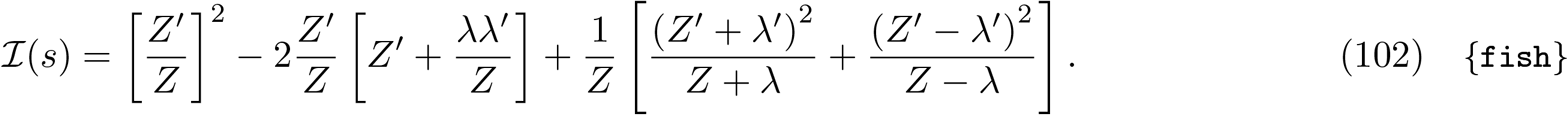

This expression simplifies significantly if we assume that *λ*_⊥_(*s*) ≈ 0, since then *Z* ≈ *σ*^2^, *Z*^′^ ≈ 0, and

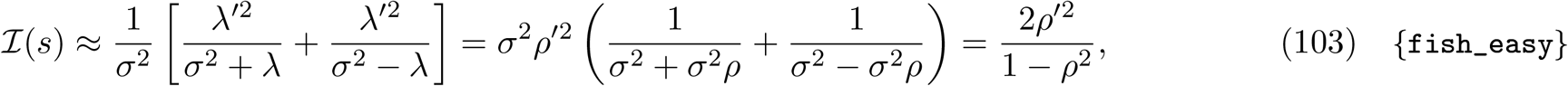

which is Eqn 32. For *ρ* = (1 − *b*)*e*^−*s/γ*^ + *b* as in Eqn 33 we get, in terms of normalized distances 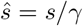,

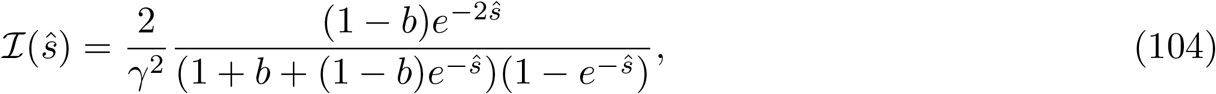

which is Eqn 35.

## 6 Acknowledgements

We thank the members of the Crimaldi and Schaefer labs, the Neuroscience Interest Group at the Francis Crick Institute, the Latham lab at the Gatsby Computational Neuroscience Unit, and the members of Odor2Action IRG3 for their feedback. We also thank Jonathan Victor for his detailed comments on this work, in particular for pointing out the need to consider the influence of out-of-phase correlations, and for helping to clarify and simplify our model assumptions.

## Supporting Information

### S1 Using other windows

To show that our results hold for other leakage-reducing windows, in Fig S1 and Fig S2 below we plot two panels from the Main Text but computed for a Kaiser window with parameter set to 16.

### S2 Governing equations and parameterization of plumes

The 2D turbulent non-dimensional flow field **u**\* = [*u*\*, *v*\*] with streamwise (*x*\*) and cross-stream (*y*\*) velocity components *u*\* and *v*\* is governed by the non-dimensional, incompressible, Navier-Stokes and continuity equations

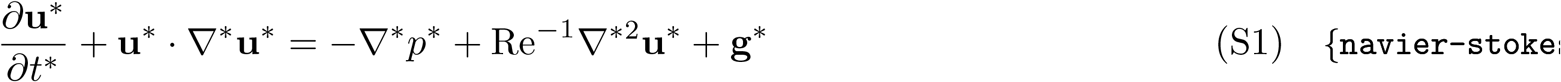

and

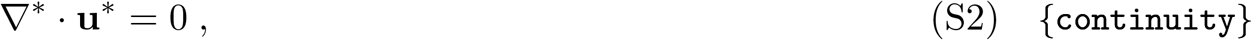

where length, time, velocity, and pressure have been non-dimensionalized by *ϕ, ϕ/U*_*o*_, *U*_*o*_, and 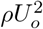, respectively. *U*_*o*_ is the mean ambient flow speed, *ϕ* is the characteristic length scale of the mixing grid (introduced below), and **g**\* = *ϕ***g***/U* ^2^. Here and elsewhere we denote vector quantities in bold face, scalar quantities in plain face, and non-dimensional quantities with an asterisk. Note the Reynolds number Re describes the relative importance of inertial and viscous flow effects and parameterizes the flow field governed by Eqns. S1 and S2,

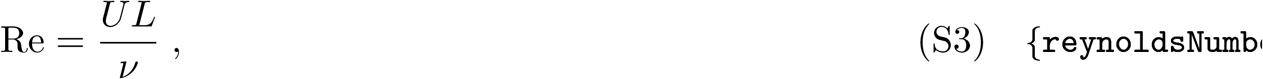

based on a characteristic velocity *U* (mean ambient flow speed *U*_*o*_ here), length scale (*L*), and the fluid kinematic viscosity *ν*. We take the mixing grid length scale *ϕ* (*pitch*, same order of magnitude as both cylinder diameters) as an appropriate characteristic length scale. The non-dimensional passive scalar concentration fields *c*\* is governed by the coupled non-dimensional, non-reacting advection-diffusion equation

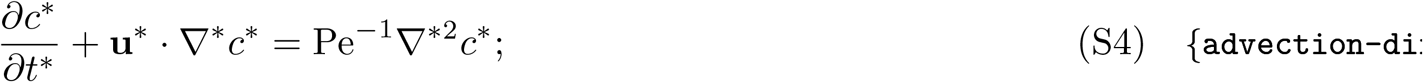

where scalar concentration *c* has been normalized by the source concentration *c*_*o*_. Velocity and time are nondimensionalized as in Eqns. S1 and S2. The Péclet number Pe parameterizes the scalar concentration field governed by Eq. S4, representing the relative importance of advective and diffusive odour mass transport as

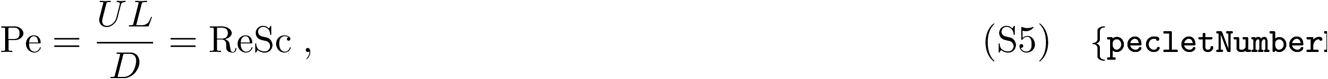

where the Schmidt number Sc is the ratio of the momentum diffusivity of the ambient fluid *ν* (i.e. kinematic viscosity) to the mass diffusivity of the given odour species in the ambient fluid *D*. Note that the instantaneous fluid velocities **u**\* and odour concentrations *c*\* are decomposed into mean 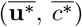 and fluctuating (turbulent) components 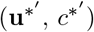 for analysis (the Reynolds decomposition).

### S3 A second set of CFD simulations

Here we provide the details the second set of computational fluid dynamics simulations that we ran to further test our approach.

The domain schematic is shown in Fig S3. We simulated the flow of a fluid inside a two-dimensional wind-tunnel 1.2 m long and 1 m, uniformly meshed with cells of diameter ∼ 2 mm. The fluid parameters were the same as those of the simulations in the Main Text. Specifically, the fluid flowed at a velocity of 0.1 m/s, with kinematic viscosity of 1.8 × 10^−5^ m^2^*/*s.

Vorticity was introduced into the fluid by twelve cylindrical obstacles of 38 mm diameter, equally spaced vertically with a gap of 38 mm at a horizontal distance of 20 cm from the inlet. Taking the gap between obstacles as the length scale yielded a Reynolds number of ∼ 200.

Point sources of odour (actual diameters were that of the cells of the mesh, ∼ 2 mm) were placed downstream of the obstacles at a horizontal distance of 40 cm from the inlet at various vertical locations. Just like the simulations in the Main Text, we simulated 16 odour sources, located symmetrically around the midline and spaced 8 mm apart, similar to the 7.5 mm spacing used in the Main Text. Like those simulations, the molecular diffusivity of odours was 1.5× 10^−5^ m^2^*/*s. Fluid flow over these odours sources carried odour downstream where their concentration profiles were detected by simulated probes.

The simulations in the Main Text were carried out using COMSOL. The supplementary simulations were performed in OpenFOAM [55], using the PISO algorithm for transient incompressible flow to solve the Navier-Stokes equations. Odours were modeled as passive scalars. To simulate multiple odour sources we found it more efficient to run multiple single-source simulations. As our simulations were deterministic and our scalars passive, this produced equivalent results to simulating multiple sources simultaneously.

The OpenFOAM code for our simulations are provided in the code repository associated with our manuscript at https://github.com/stootoon/fisher-plumes.

The plots in Fig S4, Fig S5 and Fig S6 below demonstrate that our results in the Main Text qualitatively hold for this second set of simulations.

### S4 Derivations of expectation-maximization updates

We fit models to a set *Y* = {*y*_1_, …, *y*_*N*_} of observed correlations at a given harmonic and intersource distance. The parameters for each model can be written as *θ* = {*ι, σ, θ*^+^, *θ*^−^} . The intermittency level *ι* and noise distribution standard deviation *σ*, are common to all the models. The remaining parameters, grouped into two sets *θ*^+^ and *θ*^−^, are specific to each model and are used to fit the positive and negative correlations, respectively.

To fit model parameters *θ* to the correlation data *Y*, we assumed that the observations they were independent and identically distributed according to Eqn 25 above. We then used maximum-*a posteriori* (MAP) estimation to estimate the values of the model parameters. That is, we maximized

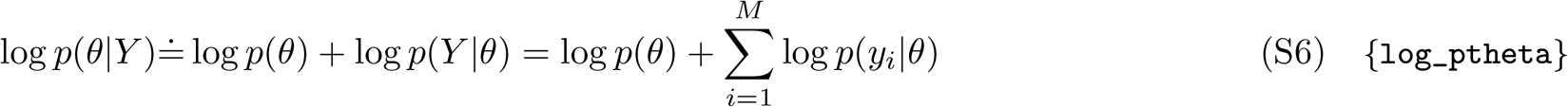

over *θ*. The dot over the first equal sign indicates that the equality is up to the additive constant, namely − log *p*(*Y*) which does not depend on *θ*. The expansion into the sum in the last term is due to our assumption of independence of the observations.

To maximize Eqn S6 for *θ* we must account for the variables *z*_1_ … *z*_*N*_ upon which the observations depend. Because these variables are latent, we use the expectation maximization algorithm [53] to iteratively maximize a lower bound to Eqn S6. At each step of the iteration, we estimate the values of the latent variables *Z* = {*z*_1_, …, *z*_*N*_} (the E-step), and then use those values to estimate the parameters *θ* of the model. We iterate until the change in parameters is less than a predefined tolerance.

Expectation maximization can be seen as coordinate ascent on the *negative variational free energy* [40]. This quantity relates a distribution *q*(*Z* | *Y, θ*) on the latents to their posterior distribution *p*(*Z*|*Y, θ*). It has two equivalent forms

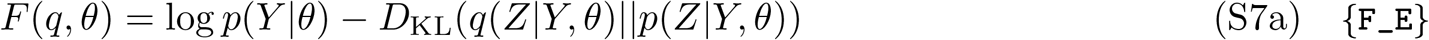

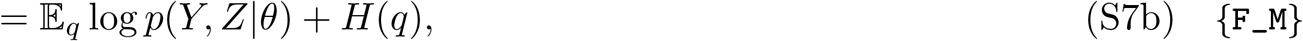

where *H*(*q*) is the entropy of the proposed distribution. From Eqn S7a and the non-negativity of the KL divergence we can see that the free energy is a lower bound on the log likelihood. During the E-step, we maximize this lower bound with respect to *q*. During the M-step, we use the form of the free energy in Eqn S7b to maximize it with respect to *θ*.

To perform expectation maximization we require a distribution *q*(*Z*| *Y, θ*) to model the posterior probability of the latent variables. We will assume that this distribution factorizes, so that

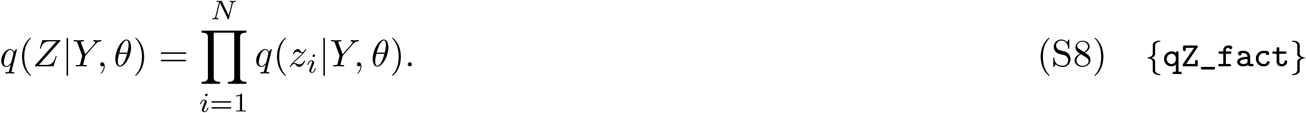

To specify *q*(*z*_*i*_|*Y, θ*) we will use a distribution that puts all the probability mass for each variable on the most likely value, whether 0 or 1.

#### S4.1 E-step

Maximizing Eqn S7a with respect to *q* is equivalent to minimizing *D*_KL_(*q*(*Z*| *Y, θ*) || *p*(*Z*| *Y, θ*)). We can write this as

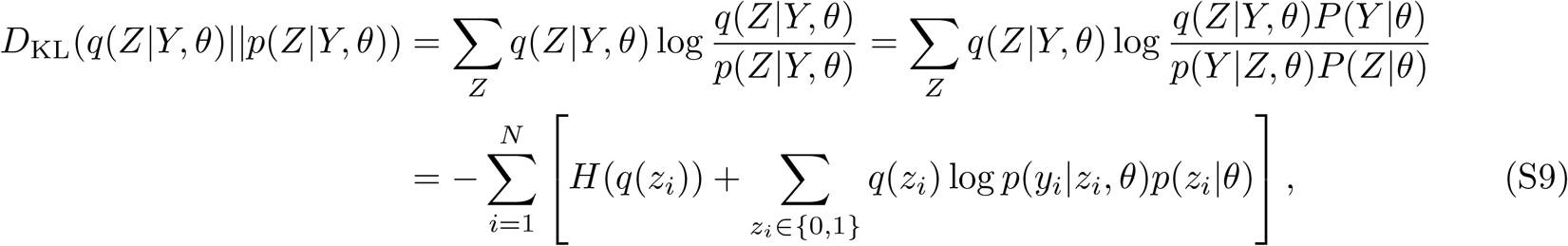

where we’ve shortened *q*(*z*_*i*_ *Y, θ*) to *q*(*z*_*i*_). Minimizing this term over *z*_*i*_ means maximizing each of the terms in square brackets. The entropy terms are all zero because *q* puts all the probability mass on zero or one. To determine whether *z*_*i*_ should be set to zero or one, we maximize the second term, which yields

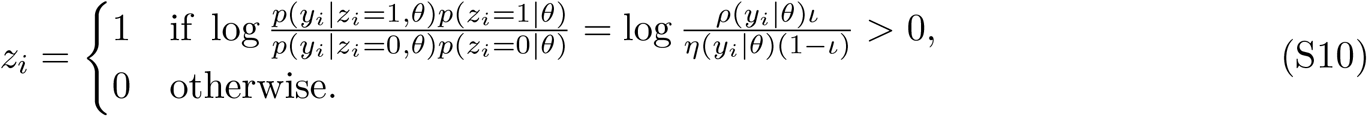

After rearranging, this is Eqn 43 in the Main Text.

#### S4.2 M-step

During the M-step we maximize Eqn S7b plus the log prior with respect to the parameters *θ*. For our model the entropy term is always zero so we need only maximize the joint distribution of observations and latents given the parameters. Also, we only use a prior distribution for the intermittency parameter *ι*. So,

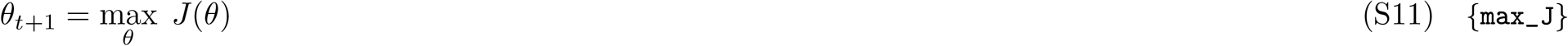

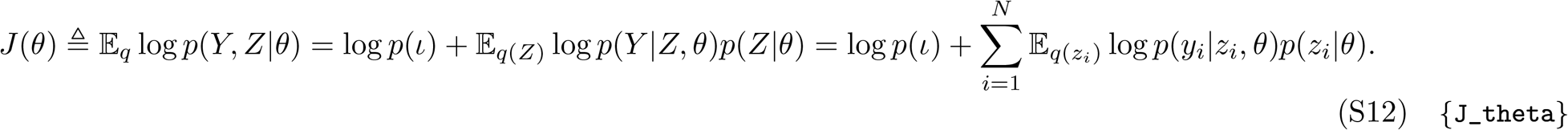

We can split the sum into components over those observations marked as correlations in the E-step, indexed by the set *I*^1^, and those marked as noise, indexed as *I*^0^:

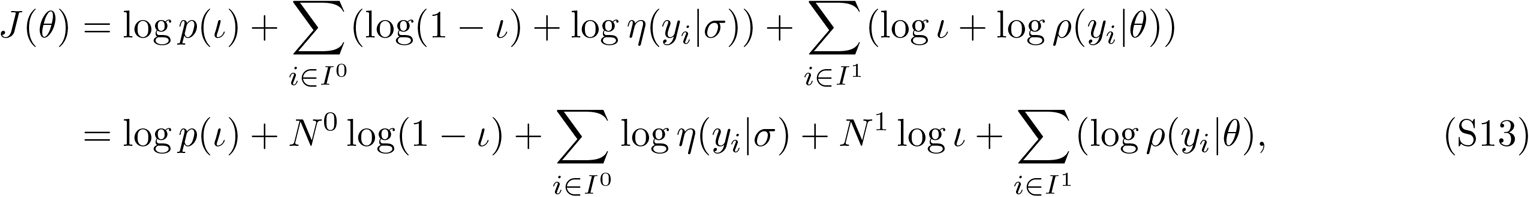

where *N* ^0^ = | *I*^0^| is the number labeled as noise, and *N* ^1^ = | *I*^1^| is the number of observations labeled as correlations. We can split this further by the sign of the correlations

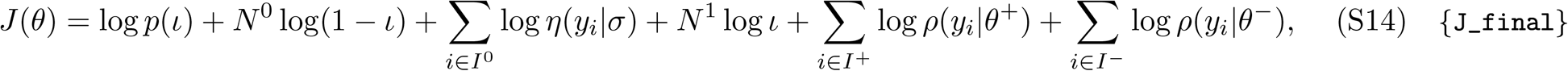

where *I*^+^ ⊂ *I*^1^ indexes the subset of the correlations that are positive, and *I*^−1^ ⊂ *I*^1^ indexes the subset that are negative. Letting *N* ^+^ = | *I*^+^| and *N* ^−^ = | *I*^−^| be the sizes of these sets, we have *N* ^+^ + *N* ^−^ = *N* ^1^.

We now maximize Eqn S14 with respect to the parameters.

#### S4.2.1 Intermittency parameters

*ι* **update**. We put a Beta prior on *ι* with mean and sample size hyperparameters, respectively, so that

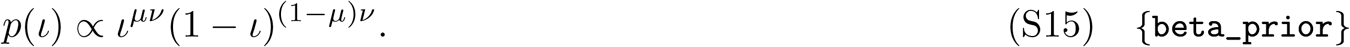

Then setting the gradient of *J*(*θ*) with respect to *ι* to zero gives

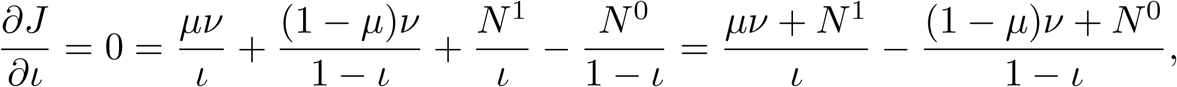

Solving for *ι* yields

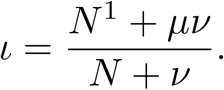

Substituting 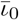for *μ* and *Nα* for *ν* gives Eqn 45.

*σ*^2^ **update**. To update the standard deviation *σ* of the noise distribution we compute

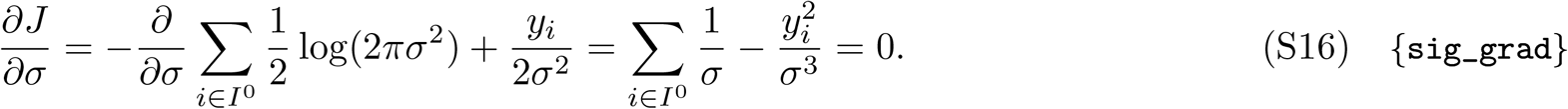

Solving for *σ*^2^ yields Eqn 46.

##### S4.2.2 Correlation parameters

From Eqn S14, maximizing the correlation parameter *θ*^+^ and *θ*^−^ can be done separately by maximizing the likelihood of the positive, and negative observations, respectively, that were marked as correlations in the E-step. This was done analytically for the Exponential model and numerically for the remaining two models.

###### Exponential model

The Exponential model parameters are *θ*^+^ = *λ* and *θ*^−^ = *μ*, with correlation distribution

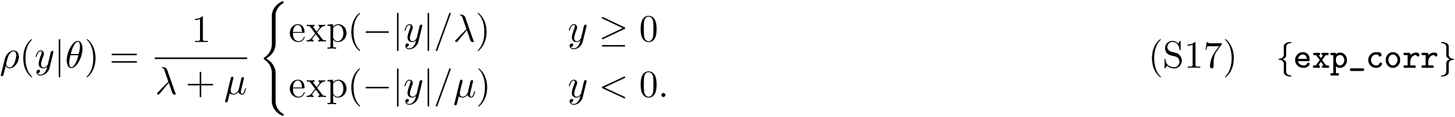

To determine *λ*, we consider a positive correlation *y*_*i*_ *>* 0. Then

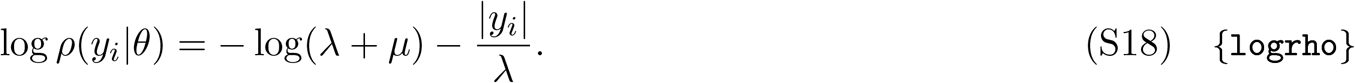

Taking the gradient of Eqn S14,

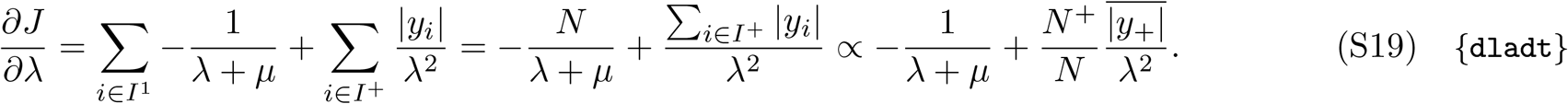

Here 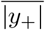 is the average positive correlation. Similarly,

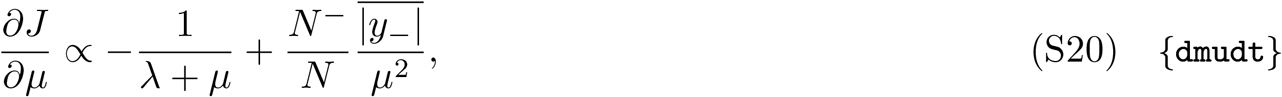

where 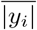 is the average negative correlation. Setting Eqn S19 and Eqn S20 to zero, we get

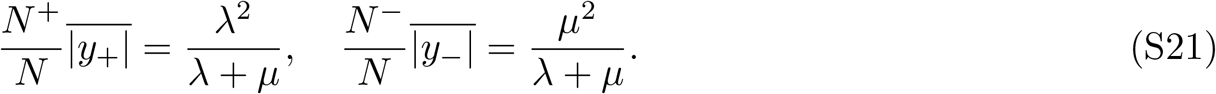

Solving for *λ* and *μ* gives the updates in Eqn 47.

###### Gamma model

The model parameters are *θ*^+^ = {*λ, k*} and *θ*^−^ = {*μ, m*}, with correlation distribution

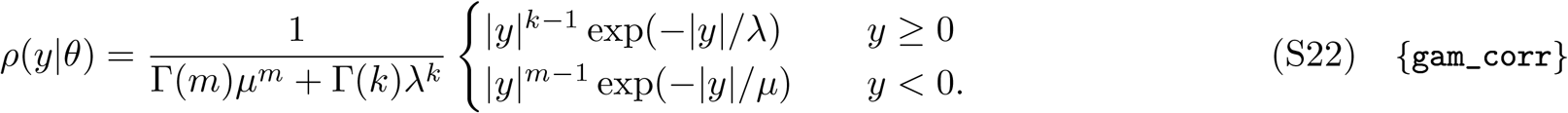

Taking the negative logarithm and summing over correlations by sign gives Eqn 48b, which we numerically minimize.

###### Generalized inverse Gaussian model

The model parameters are *θ*^+^ = {*λ, k, α*} and *θ*^−^ = {*μ, m, β*}, with correlation distribution

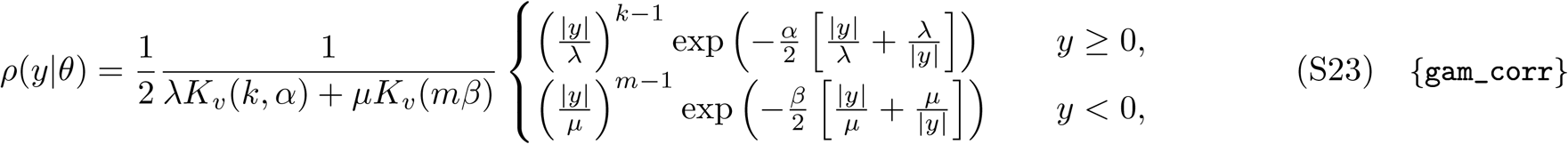

where *K*_*v*_(*a, b*) is the modified Bessel function of the second kind of real order *a* evaluated at *b*. We use the negative logarithm of this distribution, summed over correlations by sign, in the damped update of Eqn 49a.

### S5 Miscellaneous

#### S5.1 Implications of temporal stationarity

Our assumption that odour concentration profiles are temporally stationary has several important implications. First, it implies that the marginal distribution of the sine and cosine coefficients must be the same,

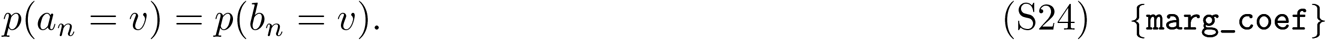

To see why, consider computing coefficients *a*_*n*_ and *b*_*n*_ at two time points *t*_2_ *> t*_1_ separated by one-quarter the period of the *n*’th harmonic. Note that a sine waveform starting at the earlier time point becomes a cosine waveform at the later one. This means that the statistics of the sine coefficients at the earlier time point are the same as those of the cosine coefficients at the later time point, that is 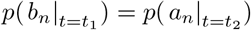. By stationarity we know that the statistics of the sine coefficients at the two time points are the same, that is 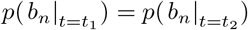. Combining these equalities yields Eqn S24.

A second implication is that the coefficients at non-zero frequency have mean zero. Intuitively, this is because any other value would introduce a non-stationarity into the plume signal that could not be canceled by any combination of the other harmonics due to their linear independence. Formally, by the linearity of Fourier decomposition the coefficients for the mean plume signal are the means of the coefficients for the plume. By stationarity, the mean plume signal is constant in time. Therefore, the mean value of the coefficients at non-zero frequencies must be zero. That is,

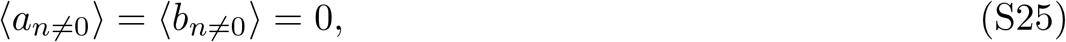

where expectations are over time windows.

A third implication is that the sine and cosine coefficients are uncorrelated. Intuitively, this is because the sine and cosine coefficients together determine the phase of the sinusoid at a given harmonic, and any correlation between the coefficients would produce non-stationarity in the phase distribution of the signal. Formally, consider computing correlations at time points *t*_2_ *> t*_1_ separated by a quarter period as before. As we previously noted, a sine waveform at *t*_1_ becomes a cosine waveform at *t*_2_. A cosine waveform at *t*_1_, however, becomes a minus sine at *t*_2_. Therefore *b*_*n*_ at *t*_1_ maps to *a*_*n*_ at *t*_2_, while *a*_*n*_ at *t*_1_ maps to − *b*_*n*_ at *t*_2_. We thus have

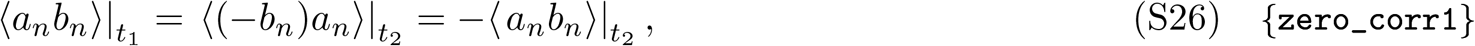

where the expectations are over *t*_2_ *> t*_1_ pairs separated by a quarter cycle. By stationarity the statistics at *t*_1_ and *t*_2_ must be the same. Therefore,

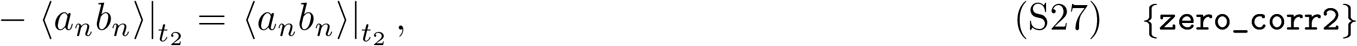

and we arrive at

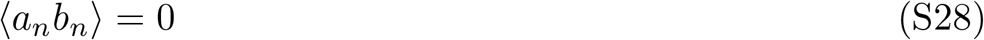

where expectation is over time. Related to this point, a final implication involves the correlations of coefficients from different sources. Applying the same considerations of the two time points *t*_1_ and *t*_2_ to the coefficients *c*_*n*_ and *d*_*n*_ from a second source, we find that

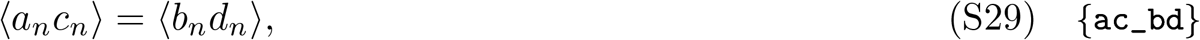

And

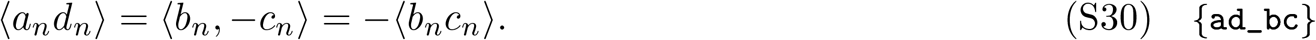

#### S5.2 Residual coefficients ar e uncorrelated

The correlation of the residual coefficients in Eqn 75 is

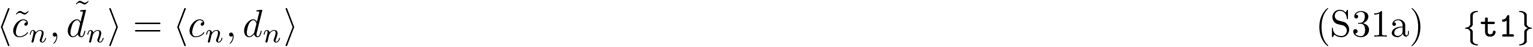

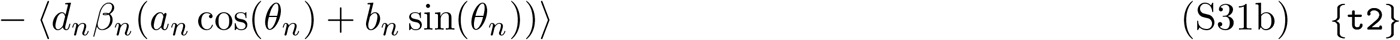

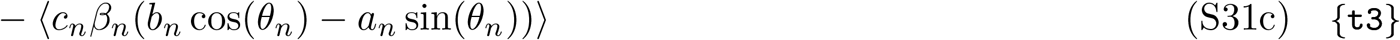

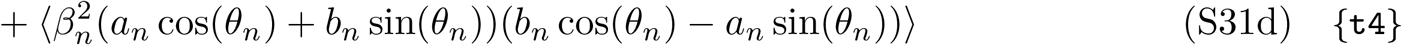

To see that this correlation is zero, note first that the right-hand side of Eqn S31a is zero since *c*_*n*_ and *d*_*n*_ are uncorrelated (Eqn 11). Next, dropping the 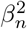 term from Eqn S31d and collecting terms we have

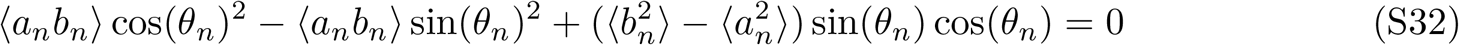

since *a*_*n*_ and *b*_*n*_ are uncorrelated, and have the same variance. Finally, combining Eqn S31b and Eqn S31c and dropping *β*_*n*_, we have

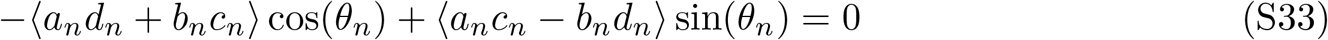

by the implications of temporal stationarity in Eqn S29 and Eqn S30. Therefore, the terms in Eqn S31 are all either individually or in combination zero, and we conclude that their sum is zero.

#### S5.3 Fits at very low frequencies

The data in Fig 7G suggests that the fits at 1 Hz are poor. We’ve highlighted those fits in Fig S8 below. These poor fits are likely due to the window size, as expanding it to 2 seconds improves the 1 Hz fits; see Fig S9 below.

#### S5.4 Converting Fourier coefficients to trigonometric coefficients

The STFT performed on plumes provides (discrete) Fourier transform coefficients. To convert the Fourier coefficients *X*[*k*] of a length *L* signal *x*[*n*] to the equivalent trigonometric re<u>present</u>ation we used the fact that the (discrete) Fourier transform of a real signal of length *L* satisfies *X*[*k*] = *X*[*L* − *k*], where the bar indicates complex conjugation. If we write each Fourier coefficient as *X*[*k*] = *u*_*k*_ + *jv*_*k*_ where *j* is the imaginary unit, we have

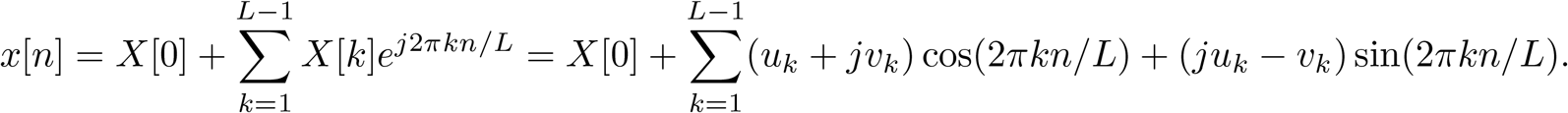

Then, if the signal is of odd length, we have

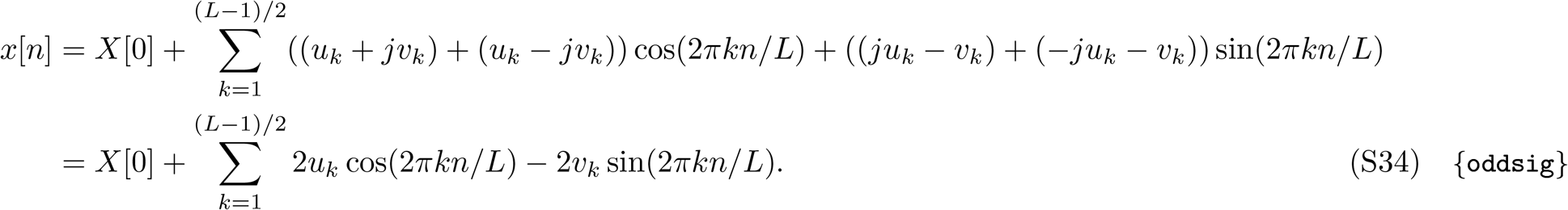

If the signal is of even length, we have

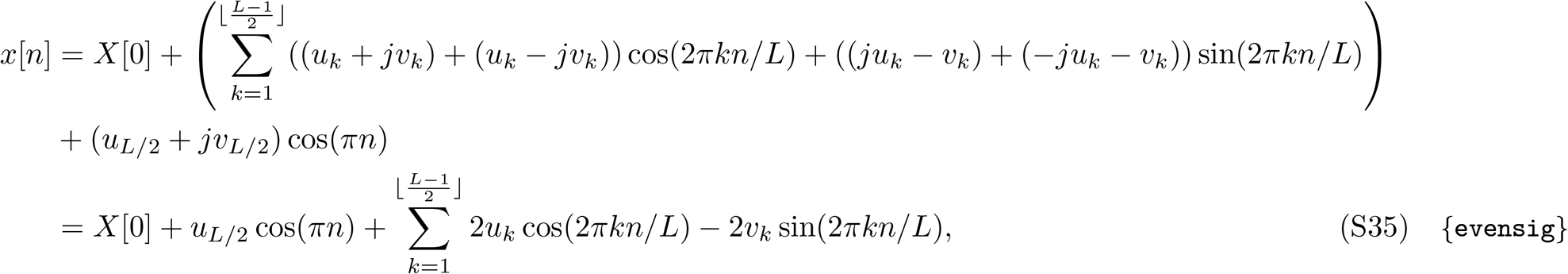

where ⌊·⌋ is the floor function. The last equality holds because *v*_*L/*2_ = 0 since *X*[*L/*2] is the correlation of the signal with exp(*jπn*) = cos(*πn*), which is purely real. We can then combined Eqn S34 and Eqn S35 to write the signal in terms of sine and cosine coefficients as in Eqn 36.

#### S5.5 Effect of windowing on covariances

We will perform the computation in continuous time for simplicity. Let two plumes generated at locations *p* and *q* be *x*_*p*_(*t*) and *x*_*q*_(*t*). We will take the full duration of these plumes to be the time interval [0, *T* ], where we’ll set *T* = 1 for convenience. Performing Fourier decompositions we have

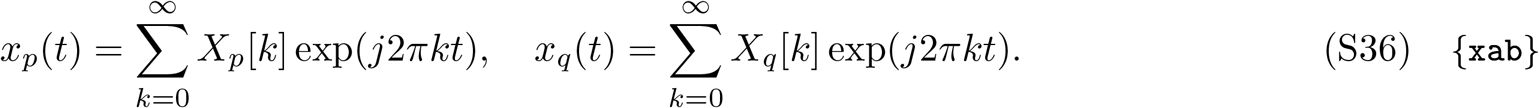

Plumes are stochastic so *X*_*p*_[*n*] and *X*_*q*_[*n*] are random variables. The covariance kernel *K*(*p, q, n*) determines how these variables are related,

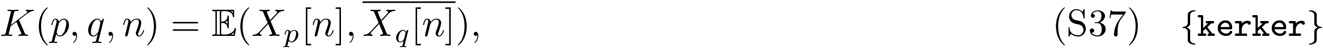

where the expectation is taken over time windows and the overline means complex conjugation. For convenience we will use this definition of the covariance kernel, relating complex Fourier coefficients, rather than Eqn 54 used in the Main Text, relating the expression of the Fourier coefficients as trigonometric functions. The animal is unable to observe the entire plume and only sees a windowed version of it. Using tilde to indicate a windowed quantity, the animal observes the plume from location *p* as

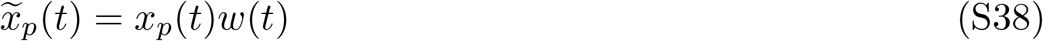

where *w*(*t*) is the windowing function that truncates the plume to a smaller time window, applies exponential filtering etc. Windowing a signal convolves its spectrum with that of the window,

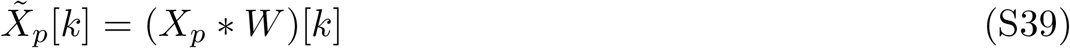

making the resulting kernel

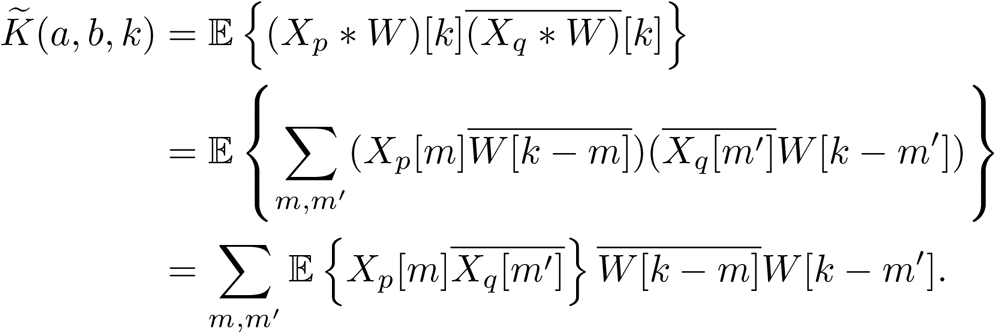

The expectation in the expression above is the covariance of coefficients from plumes at two different locations for two frequency components *k* and *k*^′^. When comparing the coefficients of a plume to itself, the covariance of coefficients at different frequency components will tend towards 0 if the interval over which the coefficients are computed is large enough [56]. We will assume that this condition is satisfied. Since the covariance between plumes from different sources will be weaker than that of a plume with itself, we can take this covariance to be 0 when comparing different frequency components. That is,

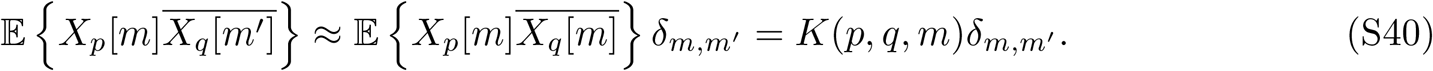

We then have

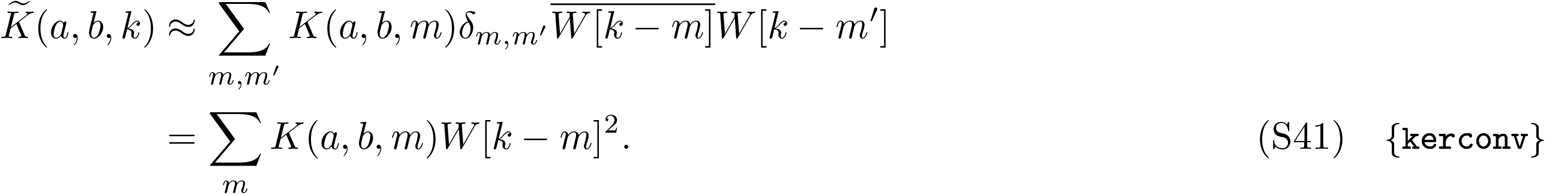

Thus the kernel 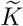 of the windowed signal is approximately the convolution of the true kernel *K* with the squared amplitude spectrum of the windowing function. Since the relevant timescale here is that of an entire plume (10s of seconds), rather than the window (*O*(1 second)), the approximation is likely to be very good in practice and we will treat the approximate equality as an equality in what follows. If the true kernel is decomposable as a product of a spatial and a frequency kernel,

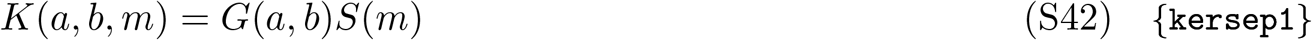

then the windowed kernel is also decomposable

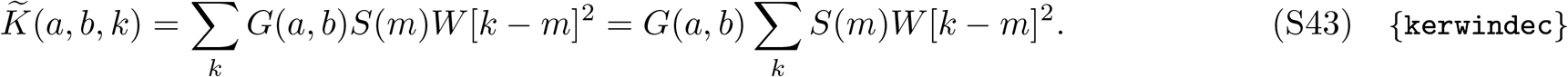

From Eqn 32 the spatial information in the kernel is related to its derivative with respect to the spatial separation |*a* − *b*|, which is

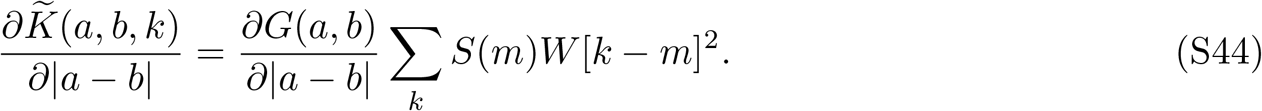

The effect of the window is thus a scaling of the kernel by the convolution of the frequency kernel *S*(*m*) with the squared spectrum of the window. Since the information content only depends on the shape of the kernel as a function of distance and not its absolute scaling, the spatial information in the windowed kernel is the same as in the un-windowed case.

If, however, the kernel is not decomposable as Eqn S42, then its shape, and thus its information content, can be changed by windowing. For example, consider a kernel which exhibits spatial decay with spaceconstant *L*_0_ for all frequencies components except at *k*_1_, for which the space constant is *L*_1_:

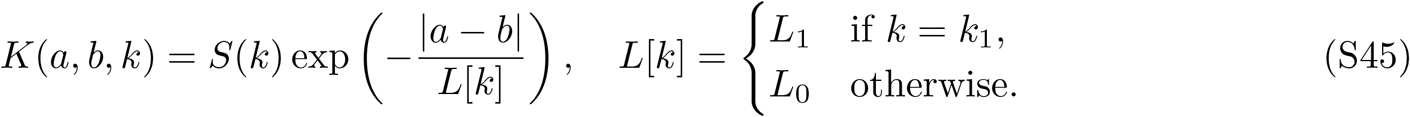

The windowed kernel is

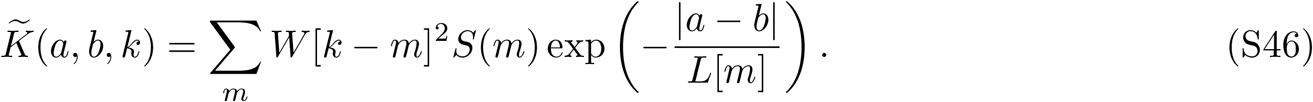

The derivative with respect to the spatial separation |*a* − *b*| is

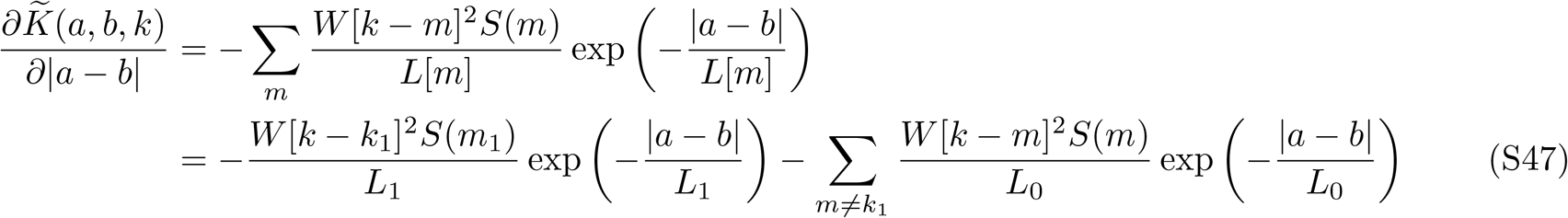

By adding and subtracting terms we can write this as a correction added to the derivative of the windowed kernel that has a single space constant *L*_0_

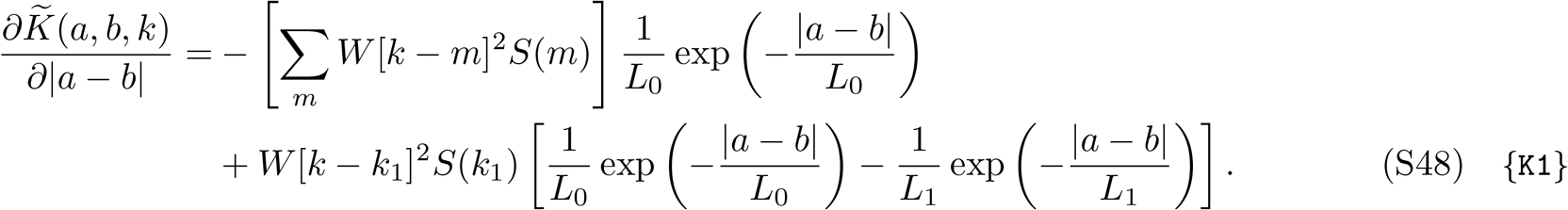

If, for example, the power *S*(*k*_1_) at the frequency component with the space-constant *L*_1_ is sufficiently low, then the correction term will be dominated by the derivative of the kernel with constant space constant. To see this, we can evaluate the derivative in Eqn S48 at *k* = *k*_1_. Doing so for the true kernel we get,

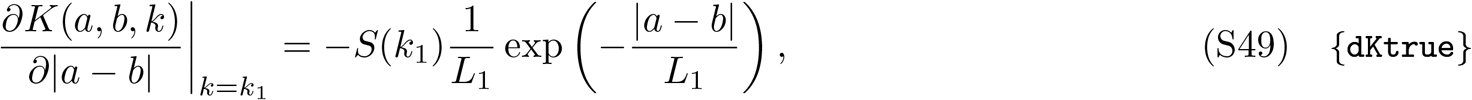

where we’ve assumed *W* [0]^2^ = 1. For the windowed kernel,

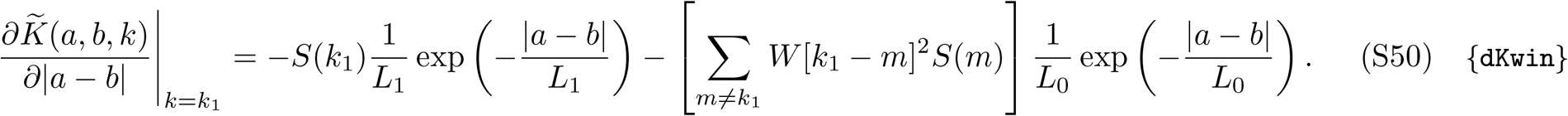

This is just the value for the true kernel in Eqn S49 corrupted by the values for neighbouring frequencies through the spectral leakage (the term in square brackets).

Thus when the true kernel is not decomposable as a product of space-only and frequency-only parts, the spatial information available in the windowed plume can differ from that in the full plume. This can result in frequencies that are particularly informative in the true signal being masked by spectral leakage in the windowed signal.

#### S5.6 Relating covariance kernels to Fisher information

How does the kernel *K*(*n, s*) used to generate a surrogate dataset determine Fisher information? In Eqn 32 we have expressed it in terms of the average in-phase correlation *ρ*_*n*_(*s*). To express Fisher information in terms of the covariance kernel *K*(*n, s*), we multiply the numerator and denominator by *σ*^4^, to get

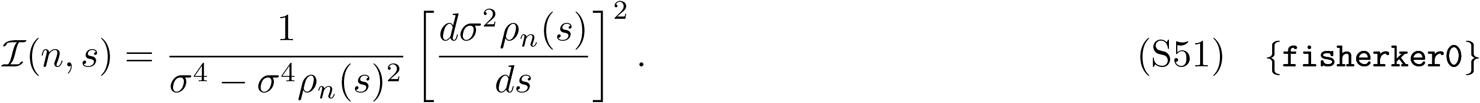

The variance corresponds to the value of the kernel at an intersource separation of 0. Therefore,

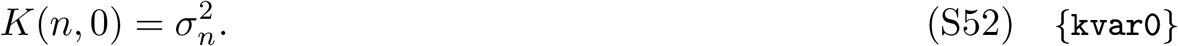

The values of the kernel at *s >* 0 determine the covariance of coefficients from different sources. Therefore, from Eqn 17b we have that

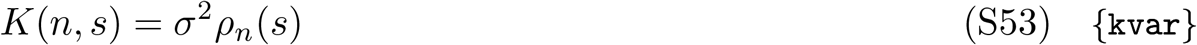

Substituting Eqn S52 and Eqn S53 into Eqn S51 gives the Fisher information interms of the kernels as

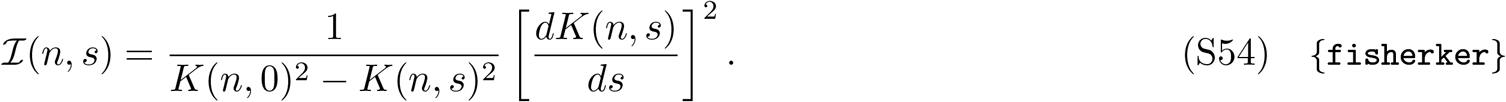

Thus we can change the amount of spatial information present in the correlations by changing the shape of the kernel, as expected. For example, we can consider a kernel that does not change with intersource distance. In that case, *dK*(*n, s*)*/ds* = 0. Substituting this into Eqn S54 (taking appropriate care of the zero in the denominator) reveals the Fisher information to be zero, as expected.

A more interesting kernel is one that can be decomposed into a product of a term *S*(*n*) that depends only on the frequency component *n*, and a term *G*(*s*) that depends only on the spatial separation:

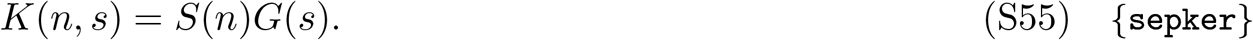

The first term in the product, *S*(*n*), reflects the variance of the coefficients of the *n*’th frequency component. The second term, *G*(*s*), reflects the correlations at intersource separation *s*. Because this term is not a function of the frequency component *n*, correlations at all frequencies change the same way with intersource separation, as observed in Fig 4F. For such ‘separable’ kernels, all frequencies are equally informative. Indeed, substituting Eqn S55 into our new expression for Fisher information in Eqn S54 yields

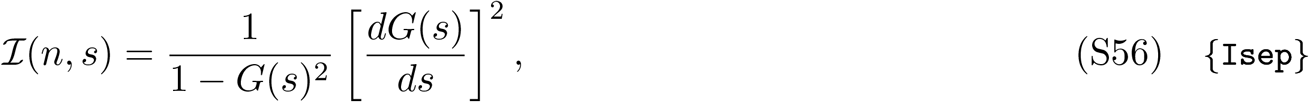

which depends only on the intersource separation *s*, and not the harmonic *n*.

### S6 Figures

**Fig S1.** Frequency decomposition of correlations for the Main Text data, using a Kaiser-16 window. Compare to Fig 9.

**Fig S2.** Fisher information vs. frequency for the Main Text data, using a Kaiser-16 window. Compare to Fig 12.

**Fig S3.** Description of the second set of computational fluid dynamics simulations. **(A)** Domain schematic and simulation parameters. Flow was from left-to-right. Vorticity was introduced into the flow by twelve cylindrical obstacles, vertically spaced evenly at a horizontal distance of 20 cm from the inlet (see also panel B). Point odour sources were placed at various vertical locations on the dashed line 40 cm horizontally from the flow inlet. Flow over these sources carried the odours to downstream probes. **(B)** Details of the obstacles and the simulation mesh. The cylindrical obstacles were 38 mm in diameter and evenly spaced 38 mm apart. The resulting center-to-center distances of 76 mm defined the pitch. The domain was discretized using a mesh with 2 mm resolution.

**Fig S4.** Example plumes and concentration profiles for the supplementary set of simulations. Compare to Fig 3.

**Fig S5.** Frequency decomposition of correlations for the second set of simulations, using a 1s Hann window. Compare to Fig 9.

**Fig S6.** Fisher information vs. frequency for the data in the second set of simulations, using a 1s Hann window. Compare to Fig 12. The fits are at for intersource distances of 0.1, 0.4 and 0.7 *ϕ*. The intersource distances for the supplementary data start at lower pitch values than for the data in the Main Text because the pitch for the supplementary simulations is *∼*3*×* larger.

**Fig S7.** Amplitude spectra of odour concentration profiles from all odour sources measured at the probe location, for the two simulated flows and the three surrogate datasets used in the text. Discrete Fourier transforms were computed for consecutive 1-second windows that overlapped by 500 msec, amplitudes were averaged and scaled to have the same value at 1 Hz. Surrogate datasets are indexed by their information content (‘all=‘: all frequencies equally informative; ‘high*>*low’: high frequencies more informative than low frequencies). The surrogate datasets (all =) and (high *>* low) were used in Fig 11 panels B, and C, respectively.

**Fig S8.** Modeling the distribution of observed correlations. As in Fig 7 but showing the fits to the 1 Hz data, highlighting the poor fits to the data.

**Fig S9.** Modeling the distribution of observed correlations. As in Fig 7 but showing the fits to the 1 Hz data, and when computing all statistics over 2-second Hann windows instead of the 1-second windows used in Fig 7.

**Fig S10.** Coupling of concentration profiles from two sources at each frequency as a function of intersource separation, expressed in terms of **(A)** phase and **(B)** strength of the coupling. **(C)** Strength (saturation) and phase (hue) together. **(D)** Out-of-phase correlations, computed as *β* sin(*θ*).

**Fig S11.** Velocity autocorrelation functions and corresponding integral length scales (*L*_*U*_) for the simulations in the Main Text, evaluated along the midline (*y* = 0) at *x* = 0 (the *x* location of the odour sources, labeled ‘origin’) and at the probe location (labeled ‘probe’). **(A)** Velocity in the *x*-direction (parallel to the mean flow) autocorrelated along the same direction. There are fewer data points at the ‘probe’ location since the largest *x*-displacement at that location is to the origin, while the largest *x*-displacement for the ‘origin’ extends past the probe location. **(B)** Velocity in the *y*-direction (perpendicular to the mean flow), autocorrelated along the same direction. The maximum displacement is approximately half that of panel A because the data in that panel spans the entire width of the simulation domain, while the data in this panel only extends from the midline to the upper and lower boundaries.

## Notes

### Competing Interest Statement

The authors have declared no competing interest.

